# Transcriptional co-activator regulates melanocyte differentiation and oncogenesis by integrating cAMP and MAPK/ERK pathways

**DOI:** 10.1101/2020.07.14.202697

**Authors:** Jelena Ostojić, Tim Sonntag, Billy Ngyen, Joan M. Vaughan, Maxim Shokirev, Marc Montminy

## Abstract

The cyclic AMP pathway promotes melanocyte differentiation in part by triggering gene expression changes mediated by CREB and its coactivators (CRTC1-3). Differentiation is dysregulated in melanomas, although the contributions of different cAMP effectors in this setting is unclear. We report a selective differentiation impairment in CRTC3 KO melanocytes and melanoma cells, due to downregulation of OCA2 and block of melanosome maturation. CRTC3 stimulated OCA2 expression via binding to CREB on a conserved enhancer, a regulatory site for pigmentation and melanoma risk in humans. Response to cellular signaling differed between CRTC3 and its family members; CRTC3 was uniquely activated by ERK1/2-mediated phosphorylation at Ser391 and by low levels of cAMP. Phosphorylation at Ser391 was constitutively elevated in human melanoma cells with hyperactivated ERK1/2 signaling; knockout of CRTC3 in this setting impaired anchorage-independent growth, migration and invasiveness while CRTC3 overexpression supported cell survival in response to MAPK inhibition by vemurafenib. Human melanomas expressing gain of function mutations in CRTC3 were associated with poorer clinical outcome. Our results suggest that CRTC3 inhibition may provide benefit in the treatment of hyperpigmentation and melanoma, and potentially other disorders with deregulated cAMP/MAPK crosstalk.

## Introduction

Increased melanoma risk in humans is correlated with pigmentation variants, light skin and poor tanning ability, although the molecular mechanisms are not fully understood^1,2^. Regulators of melanocyte homeostasis are frequently involved in melanoma development, as these tumors subvert core signaling pathways present in normal melanocytes during their progression^3–5^.

Elevations in intracellular cAMP upregulate melanin synthesis, DNA repair, and survival pathways in melanocytes^6^. cAMP signaling is required for both basal (constitutive) and stimulus-induced pigmentation, depending in part on the intracellular cAMP levels^7,8^; these reflect the relative activities of cAMP-generating adenylyl cyclases and hydrolyzing phosphodiesterases (PDEs). Although the underlying mechanisms are unclear, aberrant cAMP signaling has been reported in a variety of human cancers; indeed, this second messenger can have both positive and negative impact on oncogenesis^9^.

cAMP stimulates the PKA-mediated phosphorylation of CREB leading to the expression of target genes including MITF, the master melanocyte regulator and a melanoma oncogene^10–12^. In addition to stimulating the expression of genes that promote growth and survival, MITF also regulates the expression of melanogenic enzymes such as tyrosinase (TYR) and dopachrome tautomerase (DCT)^13^.

Superimposed on the effects of cAMP, the MAPK pathway also modulates melanocyte proliferation and differentiation, in part by regulating MITF activity^14–16^. MAPK and cAMP pathways show a nonlinear interaction which can be synergistic or antagonistic, depending on the signaling environment^17–20^.

The MAPK pathway is commonly hyperactivated in melanomas, due to mutations in RAS and BRAF that lead to downstream induction of the Ser/Thr kinases ERK1/2^21^. BRAF inhibitors are effective in the treatment of melanomas, but rates of acquired treatment resistance and relapse are also high^22^, pointing to the involvement of other signaling pathways that can circumvent BRAF inhibition. Within this group, the CREB pathway has been reported to confer resistance to BRAF blockers^23^.

CREB Regulated Transcription Coactivators (CRTC1-3) function as effectors of cAMP signaling. In the basal state, salt-inducible kinases (SIKs) sequester CRTCs in the cytoplasm via their phosphorylation at 14-3-3 binding sites. Increases in cAMP promote PKA-mediated phosphorylation and inhibition of the SIKs, leading to CRTC dephosphorylation, nuclear migration and recruitment to CREB binding sites^24^. In keeping with their considerable sequence homology, CRTCs have been found to exert overlapping effects on CREB target gene expression; yet knockout studies also reveal distinct phenotypes for each family member^25–28^. In melanocytes, CRTCs have been proposed as targets for treatment of hyperpigmentary disorders by virtue of their ability to mediate effects of CREB on MITF expression^29–32^.

Here we show that deletion of the CRTC3 but not CRTC1 or CRTC2 genes affects pigmentation and melanocyte fitness. In line with overlapping effects of CRTC family members, MITF expression is only marginally affected in CRTC3 KO animals. Melanosome maturation is defective in CRTC3 mutant melanocytes, reflecting decreases in the expression of the melanosomal transporter OCA2, a key regulator of pigmentation and a melanoma susceptibility gene. CREB and CRTC3 were found to stimulate OCA2 expression by binding to cAMP Responsive Elements (CREs) on a conserved distal enhancer. The specificity of the CRTC3 phenotype reflects in part the ability of this coactivator to integrate signals from both cAMP and MAPK/ERK pathways. By profiling transcriptomes of CRTC3-depleted murine and human melanoma cells, we found that CRTC3 regulates genes involved in cAMP signaling, pigmentation and cell motility. CRTC3 activity is upregulated in subsets of human melanomas, where it is associated with reduced survival. Our results suggest therapeutic benefit of targeting CRTC3 in the treatment of pigmentary disorders and cutaneous melanoma, and potentially other conditions characterized by dysregulated cAMP/MAPK crosstalk.

## Results

### Absence of CRTC3 in mice impairs melanocyte differentiation

We noticed that mice with a knockout of CRTC3 but not CRTC1 or CRTC2 had decreased fur pigmentation, (65% of wild-type (WT)) (Fig. 1a, b). Knowing the importance of the CREB pathway in regulating MITF expression and melanogenesis, we considered that CRTC3 expression in skin could exceed those of CRTC1 and CRTC2. However, all three CRTCs were comparably expressed in both whole skin and in primary melanocytes (Fig. 1c, d, Suppl. Fig. 1a). CRTC3 KO skin did not show defects in protein accumulation of MITF, or several of its target genes (TYR, DCT, PMEL), likely reflecting compensation by CRTC1 and CRTC2 (Fig. 1d). To identify changes that might explain the CRTC3 KO phenotype, we performed RNA-sequencing on WT and CRTC3 KO neonatal skins. Consistent with protein accumulation data, mRNA levels for MITF and major melanogenic enzymes were not significantly altered in the knock-out (Suppl. Table 1). Transporters were amongst the most significant down-regulated gene categories in this analysis; indeed, three of these were melanocyte-specific (Fig. 1e, 1f, 1g). Although melanocytes account for less than 10% of cells in the skin, a melanocyte-specific gene OCA2 (oculo-cutaneous albinism II) scored as the most significantly down-regulated gene in CRTC3 KOs (Suppl. Table 1). OCA2 encodes a transmembrane anion transporter that regulates melanosomal pH, and that is essential for the activity of melanogenic enzymes, in particular TYR^33,34^. Alterations in OCA2 underlie natural pigmentation variants, albinism and melanoma risk in humans^35–39^.

**Fig. 1:**
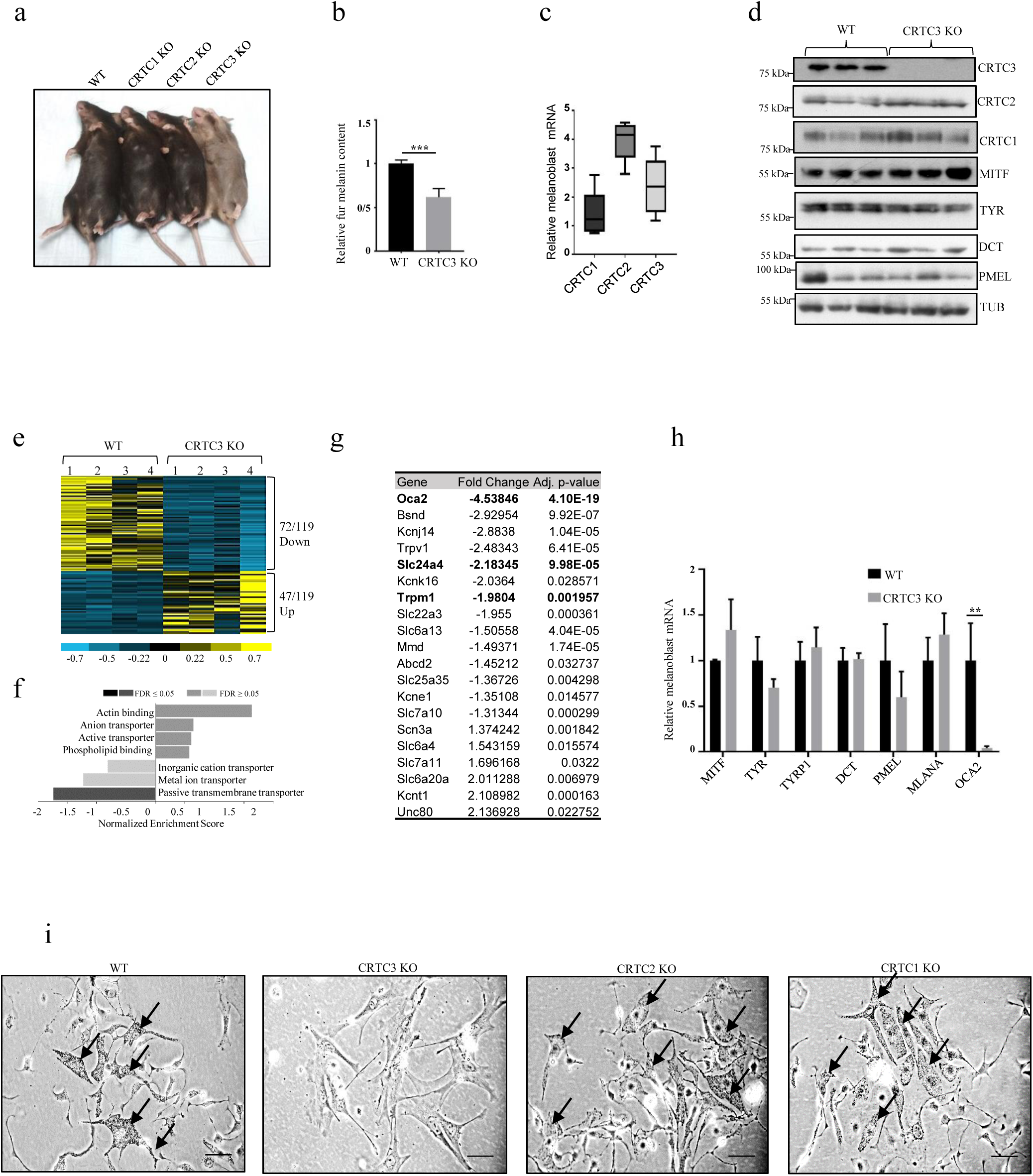
Hypopigmentation in CRTC3 knockout mice. **a)** Fur color comparison between mice with individual knockouts of CRTC family members. **b)** Quantification of melanin content in dorsal hair from WT and CRTC3 KO mice (N=3 per group). Significance determined by Welch’s t-test. **c)** RT-qPCR quantification of CRTC expression levels in primary melanoblasts from WT mice (N=5). **d)** Western blots from whole skin of WT and CRTC3 KO mice (P2) showing expression of CRTCs, melanogenic enzymes and structural proteins. **e)** Heat map of significant differentially expressed genes in RNA-sequencing experiments of WT and CRTC3 KO whole skins (P2, N=4 per group). **f)** Clustering of significant differentially expressed genes in RNA-sequencing experiments shown in panel e) **g)** List of transporters from RNA-seq experiments shown in panel e). Melanocyte-specific transporters are in bold. **h)** RT-qPCR data from sorted melanoblasts of WT and CRTC3 KO mice (P2, N=4-5 per group). Significance determined by Welch’s t-test. **i)** Primary melanocytes isolated from skins of WT, CRTC1 KO, CRTC2 KO and CRTC3 KO mice (P2), cultured for 2 weeks in differentiation media. Arrows point to intracellular melanin granules that are absent from CRTC3 culture. Bar is 50µm.

We used DCT promoter-Green Fluorescent Protein (GFP) reporter, to evaluate neonatal melanoblast populations in skins of WT and CRTC3 KO animals^40,41^. No significant differences in the percentage of GFP positive melanoblasts were seen between the two genotypes (Suppl. Fig. 1b). Similar to whole skin, OCA2 mRNA amounts were also reduced in CRTC3 KO melanoblasts, while MITF or other core melanogenic enzymes were unaffected (Fig. 1h).

We evaluated the differentiation of WT and CRTC3 KO melanoblasts following exposure to cholera toxin (CTX) and phorbol ester 12-o-tetradecanoylphorbol-13-acetate (TPA), activators of cAMP and MAPK cascades. CTX and TPA induced melanin synthesis in WT, CRTC1 KO, and CRTC2 KO melanoblasts, but not in CRTC3 KO (Fig. 1i). Rather, surviving CRTC3 KO cells expressed fibroblast markers, but not melanocyte markers (Suppl. Fig. 1c). We considered that *ex-vivo* culture may exacerbate the phenotype of CRTC3 KO mouse melanocytes due to increases in reactive oxygen species^42^. Supplementing the primary culture with catalase, an antioxidant cocktail or co-culturing with feeder keratinocytes did not rescue differentiation of CRTC3 KO melanocytes, however. These results indicate that the selective loss of CRTC3, but not CRTC1 or CRTC2 impairs melanocyte differentiation.

### CRTC3 promotes pigmentation by regulating the expression of OCA2

To further characterize the role of CRTC3 in pigmentation, we used mouse melanoma cells B16F1, which produce melanin following exposure to cAMP agonists^43^. CRISPR or RNAi-mediated depletion of CRTC3 reduced B16F1 cell pigmentation, which was rescued by CRTC3 re-expression (Fig. 2a, Suppl. Fig. 1d, e). Despite profound pigmentation defects, CRTC3 KO cells expressed comparable amounts of MITF and TYR proteins to control cells (CTRL); they were able to accumulate both proteins efficiently following FSK exposure (Suppl. Fig 1f). TYR DOPA oxidase activity was decreased in CRTC3-depleted cells, by in-gel and whole lysate analyses; re-expression of CRTC3 restored TYR activity (Fig. 2b, c). TYR activity depends on correct maturation of melanosomes^44^, prompting us to assess melanosome populations in CTRL and CRTC3 KO cells. Using transmission electron microscopy (TEM), we noted that maturation of melanosomes was severely impaired in CRTC3 KO cells as evidenced by a high number of early (I and II) versus melanized late stage (II and IV) melanosomes (Fig. 2d, Suppl. Fig. 1g). These results indicate that depletion of CRTC3 reduces pigmentation by altering an essential step in melanosome maturation, which is necessary for TYR enzyme activity.

**Fig. 2:**
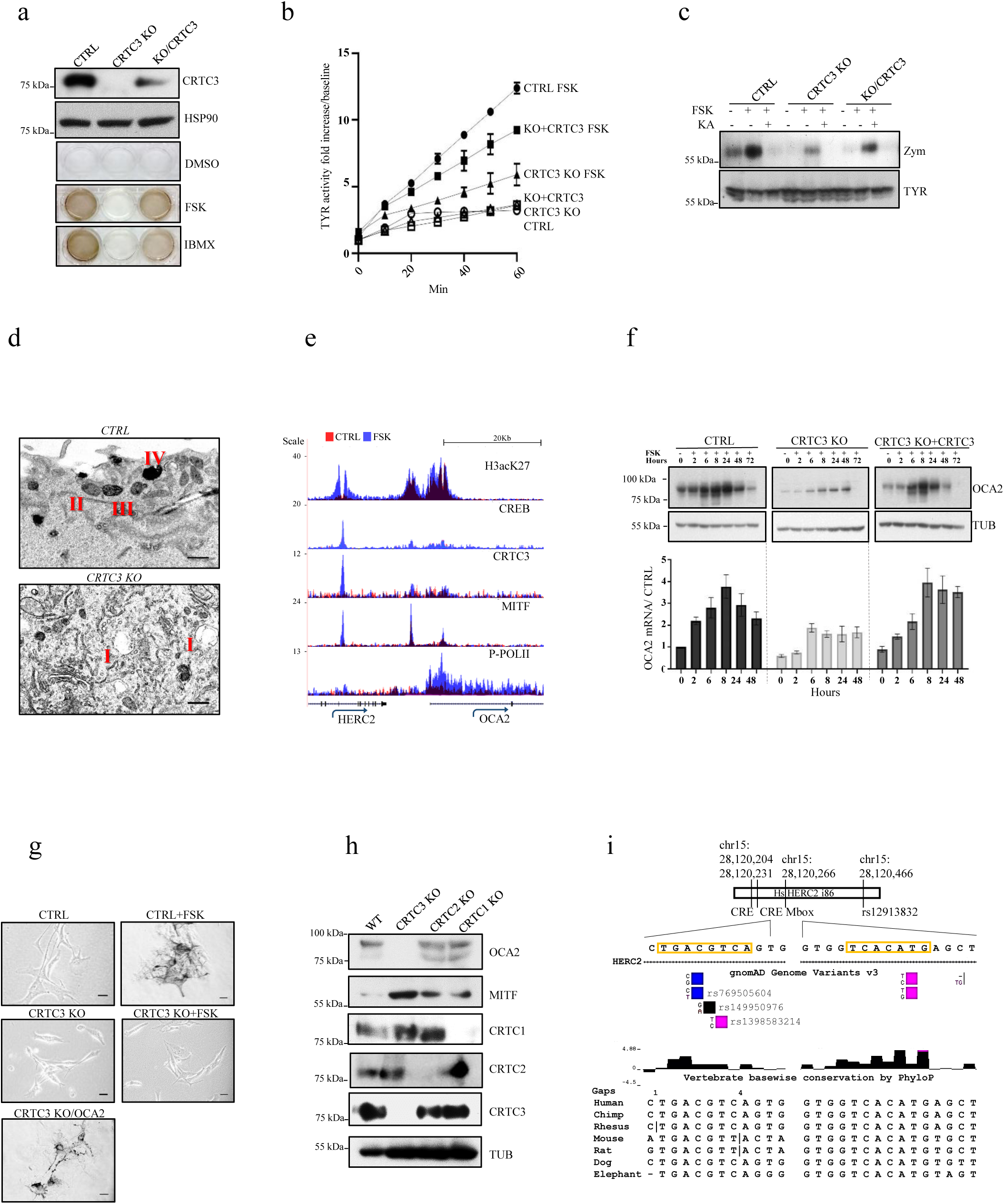
Loss of CRTC3 impairs OCA2 expression and melanosome maturation. **a)** Western blot and melanin quantification assay in CTRL, CRTC3 KO and CRTC3-rescued B16F1 cells following differentiation stimulus with 5µM FSK or 100µM IBMX for 60h. **b)** Tyrosinase activity in whole lysates of B16F1 cells with indicated genotypes and treatments (5µM FSK, 48h). **c)** Tyrosinase in-gel activity (zymography) and western blot showing TYR protein accumulation levels in B16F1 cells of indicated genotypes. Cells were treated with 5µM FSK or 500µM TYR inhibitor kojic acid for 48h. **d)** Representative TEM image of melanosome maturation stages in CTRL and CRTC3 KO B16F1 cells after 48h of 5µM FSK stimulation. Observed melanosome maturation stages are indicated with Roman lettering. Bar is 400nm. **e)** Browser plot of genomic region containing OCA2 enhancer showing occupancy of acetylated histone H3K27, CREB, CRTC3, MITF, and phospho-POLII in B16F1 cells treated with vehicle or 5µM FSK for 1h. Scale indicates normalized tag enrichment. **f)** Time course for OCA2 mRNA and protein induction upon 5µM FSK treatment, assayed through RT-qPCR and western blotting. **g)** Baseline rescue of melanogenesis in CRTC3 KO B16F1 cells transiently transfected with OCA2. CTRL and CRTC3 KO cells were treated with 5µM FSK for 60h to induce melanin synthesis. Bar is 25µm. **h)** Representative western blot of adult dorsal skin from mice of indicated CRTC genotypes showing accumulation of OCA2 and MITF. i) Schematic representation of the conserved region of intron 86 of HERC2 gene, containing the SNP rs12913832, relevant for pigmentation and melanoma risk in humans. Positions and conservation of the full CRE site and the nearest Mbox site are indicated. Validated SNP variants in the CRE site are shown and colored to indicate allelic prevalence.

To identify direct targets of CRTC3 that contribute to melanin production, we performed ChIP-seq and RNA-seq studies in B16F1 cells. Exposure to FSK triggered CRTC3 recruitment to regulatory regions for genes involved in pigmentation, vesicular trafficking, and MAPK signaling. CRTC3 was primarily recruited to CREB binding sites and in close proximity to MITF-bound M-box sites (Suppl. Fig. 1h, i). Pointing to compensation between CRTC family members, knock-out of CRTC3 decreased the expression of canonical CREB targets and MITF only modestly (Suppl. Table 2, 3). Down-regulated genes were clustered in processes involved in substrate interaction and pigmentation; most of these genes were efficiently rescued by re-expression of CRTC3 (Suppl. Fig. 1j, k).

OCA2 was prominent within the subset of CRTC3-regulated genes (Suppl. Table 3). Exposure to FSK simulated binding of CRTC3, CREB and MITF to a conserved OCA2 enhancer located in intron 86 of the upstream HERC2 gene (Fig. 2e)^45,46^. Concurrently, exposure to FSK increased occupancy of phosphorylated polymerase II over the OCA2 gene body (Fig. 2e). OCA2 mRNA amounts were increased after 2h of FSK stimulation, followed by increases in protein amounts as determined by immunoblotting with a validated OCA2 antiserum (Fig. 2f, Suppl. Fig. 2a-d). Induction of OCA2 was severely blunted in CRTC3 KO and recovered with CRTC3 re-expression (Fig. 2f). The effects of CRTC3 on OCA2 expression appear to account for reduced pigmentation since overexpression of OCA2 in CRTC3 KO cells rescued melanin production even under basal conditions (Fig. 2g). Having seen the selective loss of pigmentation in CRTC3 KO mice, we wondered if the effects on OCA2 were also CRTC3-specific. We generated a CRISPR KO of CRTC1 and assessed its melanin production and OCA2 status in B16F1 cells. While depletion of CRTC3 impacted both basal and FSK-induced OCA2 amounts, CRTC1 KO reduced FSK-induced OCA2 protein amounts only modestly. Extracellular melanin output was largely unchanged in CRTC1 KO, indicating that CRTC1 is dispensable for efficient melanin production in this setting (Suppl. Fig. 2f). OCA2 protein amounts are substantially reduced in CRTC3 KO skin compared to WT as well as CRTC1 and CRTC2 KOs by immunoblot assay (Fig. 2h). OCA2 expression in melanocytes is primarily regulated by a distal enhancer in intron 86 of the HERC2 gene. Within this enhancer, we noticed two CREB binding sites that are positioned in the conserved region of intron 86, which is also the site of long-distance chromatin loop formation with the OCA2 proximal promoter^36,45^ (Fig. 2i). The CRE site contains validated low frequency SNP variants, all of which are predicted to decrease CREB binding based on previously reported fluorescence anisotropy assays of physiologically relevant CRE motifs^47^ (Fig.2i).

### cAMP and ERK1/2 regulate CRTC3 activity

We hypothesized that CRTC3 may be selectively activated in melanocytes by signals other than cAMP or by low levels of cAMP that are otherwise insufficient to activate CRTC1 and CRTC2. ERK1/2-mediated phosphorylation of CRTC3 at Ser^391^ increases its interaction with protein phosphatase 2 (PP2A), leading to the dephosphorylation of CRTC3 at inhibitory 14-3-3 binding sites^48^. Exposure of B16F1 cells to several ERK1/2 activators (SCF, HGF, TPA) promoted the phosphorylation of CRTC3 at Ser^391^ (Fig. 3a). In line with previous observations, Ser^391^-phosphorylated CRTC3 was predominantly nuclear-localized (Suppl. Fig. 3a). Correspondingly, loss of CRTC3 blocked melanin production and re-expression of CRTC3 rescued these effects, but the overexpression of CRTC2 did not. Notably, over-expression of the hybrid CRTC2 protein containing the PP2A binding domain of CRTC3^48^ was competent to rescue melanin synthesis (Fig. 3b). Mutation of Ser^391^ to Ala reduced CRTC3 effects on melanin production relative to WT CRTC3 (Suppl. Fig. 3b).

**Fig. 3:**
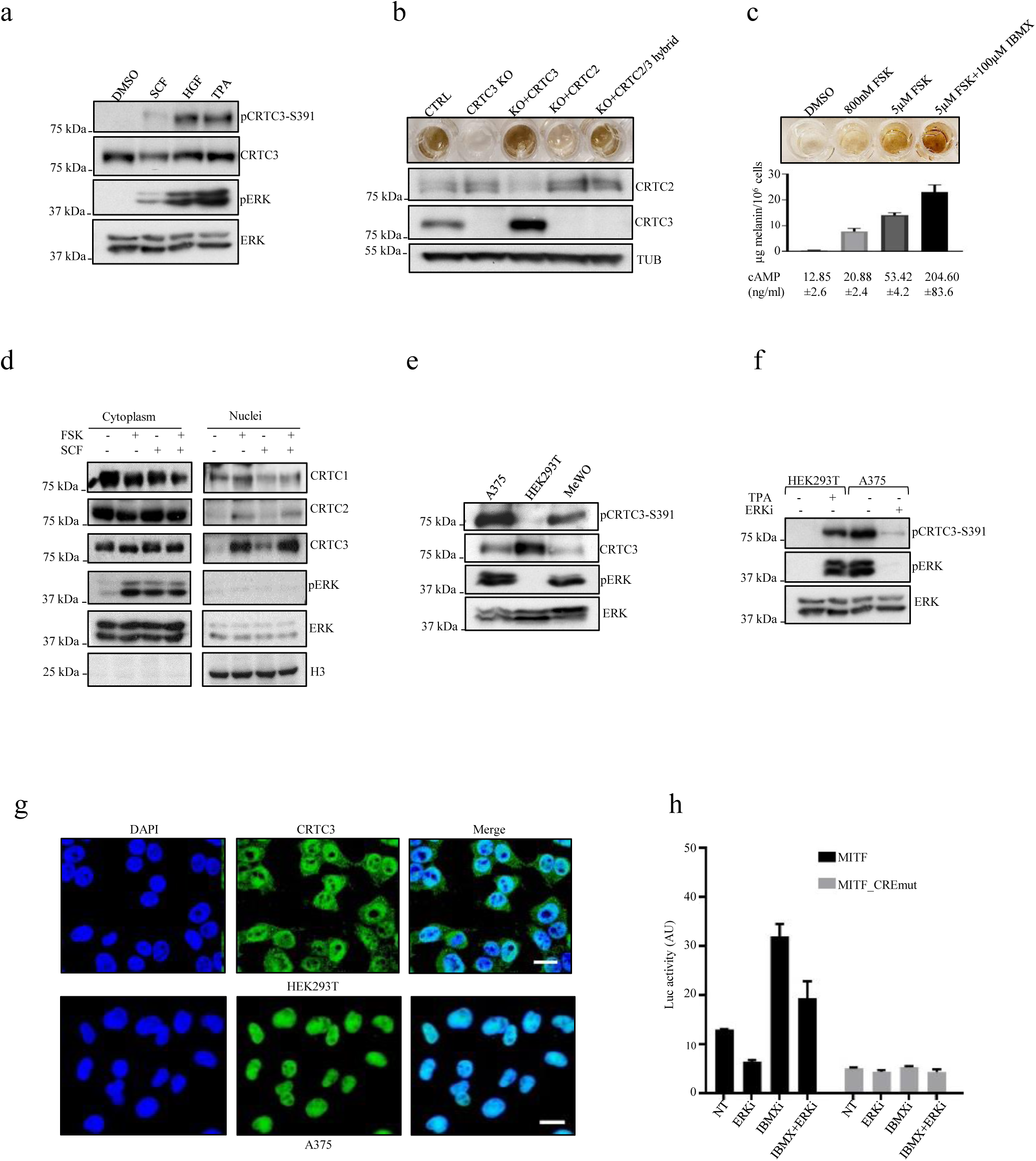
Selective induction of CRTC3 in response to ERK1/2 activation. **a)** Western blot showing phosphorylation of CRTC3 at Ser^391^ upon stimulation of ERK1/2 in B16F1 cells (SCF – stem cell factor (30ng/ml); HGF – hepatocyte growth factor (20ng/ml), TPA (200nM) **b)** Rescue of melanogenesis in CRTC3 KO B16F1 cells after transfection with indicated constructs. Cells were treated with 5µM FSK for 60h, 48h after transfection **c)** Melanin production related to cAMP content in B16F1 cells stimulated with indicated compounds for 60h. **d)** Western blot of a representative sub-cellular fractionation experiment after treatment with 800nM FSK or 30ng/ml SCF for 20min. **e) f)** Western blots of indicated human cell lines in the basal state and treatment with 200nM ERK1/2 inhibitor SCH772984 for 8h. **g)** Immunofluorescence of HEK293T and A375 cells stained with hCRTC3(414-432) antibody and DAPI. Bar is 20µM **h)** Luciferase reporter assay in A375 cells transfected with MITF proximal promoter construct. Treatments and mutations of regulatory elements are indicated. Cells were pretreated with 200nM ERK1/2 inhibitor SCH772984 for 6h before adding 100µM IBMX for 5h.

To test whether CRTC3 is activated at lower cAMP levels than other CRTCs, we exposed B16F1 cells to a range of FSK concentrations; at 800nM, FSK modestly increased intracellular cAMP levels and melanin synthesis (Fig. 3c). Low FSK (800nM) increased nuclear amounts of CRTC3 to a greater extent than CRTC1 or CRTC2 (Fig. 3d, Suppl. Fig. 3c). Low FSK treatment also stimulated ERK1/2 activation to a similar degree as treatment with SCF, which promoted nuclear translocation of CRTC3, but not CRTC1 or CRTC2 (Fig. 3d). Collectively, these results indicate that CRTC3 has increased sensitivity to both cAMP and ERK1/2 signals.

### CRTC3 contributes to oncogenic properties of transformed melanocytes by modulating intracellular cAMP

MAPK hyperactivation is found in over 90% of human melanomas^21,49^, prompting us to test whether corresponding increases in ERK1/2 activity up-regulate CRTC3. Amounts of phosphorylated CRTC3-Ser^391^ correlated with the extent of ERK1/2 activity in several human cell lines. HEK293T cells had low basal ERK1/2 activity and correspondingly low CRTC3 phospho-Ser^391^; by contrast, A375 and MeWO cells had high basal ERK1/2 activity and increased levels of phospho-Ser^391^ (Fig. 3e, 3f).

A375 human melanoma cells express an oncogenic gain of function BRAF (V600E) mutation and low levels of cAMP signaling due to increased expression of phosphodiesterases^50^, which provided us with a window to evaluate selective effects of high ERK1/2 signaling on CRTC3 activity. CRTC3 was predominantly nuclear-localized in A375 but not HEK293T cells, consistent with their relative ERK1/2 and CRTC3-Ser^391^ phosphorylation profiles (Fig. 3g). Treatment of A375 cells with ERK1/2 inhibitor decreased CRTC3-pSer^391^ amounts and correspondingly reduced CREB activity over the MITF promoter (Fig. 3f, 3h).

We tested the regulatory contribution of CRTC3 to oncogenic properties of melanoma cells by generating CRISPR-derived CRTC3 KO A375 cells (Suppl. Fig. 3d). Loss of CRTC3 had no effect on proliferation, but it impaired anchorage-independent growth in both A375 and B16F1 melanoma cells (Suppl. Fig. 3e, f, g, h). Anchorage-independent growth was also reduced in cells expressing phosphorylation-defective CRTC3-Ser^391^Ala mutant relative to WT CRTC3 (Suppl. Fig. 3i).

To determine transcriptional effects of CRTC3 in a human melanoma model, we performed RNA-sequencing on CTRL and CRTC3 KO A375 cells. Loss of CRTC3 down-regulated genes involved in cAMP signaling, cell migration and chemotaxis (Fig. 4a, Suppl. Table 4). A number of canonical CREB targets were significantly down-regulated in the KO cells, amongst these the phosphodiesterases PDE4B and PDE4D (Fig. 4a, Suppl. Table 4). PDE4 has been found to promote melanoma progression by reducing inhibitory effects of cAMP on activation of the MAPK pathway^20,50–52^. Based on its ability to stimulate PDE4 expression, we reasoned that loss of CRTC3 should increase intracellular cAMP. Supporting this idea, CRTC3 KO cells had higher basal concentrations of cAMP, higher PKA activation and greater response to intermediate concentrations of FSK (Fig. 4b, Suppl. Fig. 4a).

**Fig. 4:**
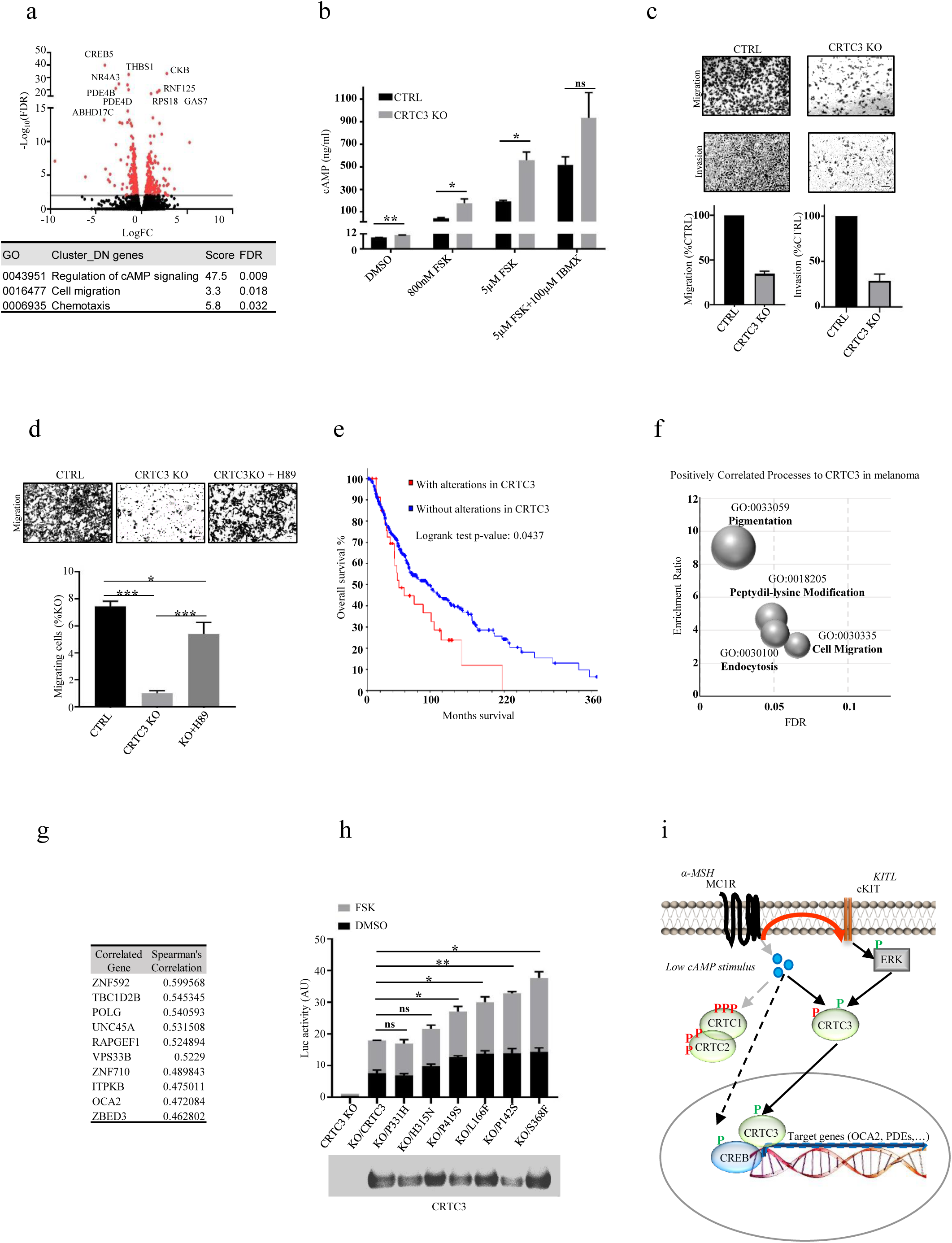
CRTC3 activity is associated with increased tumorigenic potential and decreased survival. **a)** Volcano plot of RNA-seq data showing differentially expressed genes between CTRL and CRTC3 KO A375 cells. Significance determined as adjusted p-value of ≤0.05 and Log_2_ fold changes of ≤-1 or ≥1. Table shows significant cluster enrichment of downregulated genes, listed in Suppl. Table 4. **b)** Quantification of cellular cAMP in CTRL and CRTC3 KO A375 cells treated with indicated compounds for 15 min. Significance determined by Welch’s t-test. **c)** Representative images showing migration (24h) and invasion (72h) of CTRL and CRTC3 KO A375 cells. 2% FBS was used as a chemotactic agent. Bar is 100µm **d)** Representative images and quantification of migration (24h) of CRTC3 KO A375 cells treated with DMSO or PKA inhibitor H89 (10µM). Bar is 50µm. Significance determined by one-way ANOVA and Tukey’s multiple comparisons tests. **e)** Kaplan-Meyer graph of overall survival of melanoma patients from TCGA, Firehose Legacy with and without alterations in CRTC3 (N=367). Graph and statistical analyses were obtained from cbioportal.org. **f)** CRTC3 correlation analysis in human melanoma patients from TCGA, Firehose Legacy (N=367). Bubble chart shows top positively enriched processes distributed by enrichment ratio and significance. **g)** Top 10 genes positively correlated with CRTC3 from the cohort in g). **h)** Luciferase reporter assay in B16F1 cells transfected with EVX-luc-2xCRE reporter and WT or mutated CRTC3 constructs. Cells were treated with 5µM FSK for 6h, 48h after transfection. Assayed patient mutations are indicated and shown on sequence alignment in Suppl. Fig. 4g. The experiment was run five different times overall. Data represents two replicas run in parallel, each assayed in technical duplicates and normalized by expression of transfected CRTC3. Both replicas had comparable expression of CRTC3 constructs, one of which is shown in the western blot insert. Differences in expression of the reporter constructs were compared to WT CRTC3 and significance determined by Welch’s t-test. **i)** Model for joint regulation of CRTC3 by cAMP and ERK1/2 in low cAMP state.

Increases in cAMP signaling have been shown to inhibit cancer motility, depending on the signaling context^53^. Treatment of A375 cells with cAMP agonists inhibited cell migration in a dose-dependent manner; these effects were reversed following exposure to the PKA inhibitor H89 (Suppl. Fig. 4b, c). Consistent with this observation, migration and invasiveness were reduced in CRTC3 KO cells; addition of H89 also rescued migration in this setting (Fig. 4c, d).

Small molecule inhibitors of BRAF constitute effective treatment options for patients with BRAF-mutant melanomas. Up-regulation of the CREB pathway has been found to confer resistance to BRAF inhibition^23^, prompting us to evaluate the potential role of CRTC3 in modulating the treatment response of A375 cells to BRAF inhibition by vemurafenib (PLX4032). Exposure to PLX4032 decreased cell viability comparably in CTRL and CRTC3 KO A375 cells. Notably, transient over expression of CRTC3 increased viability in cells exposed to PLX4032 but not to cisplatin, a DNA cross-linking compound (Suppl. Fig. 4d, e).

### Spontaneous CRTC3 mutations in human melanomas modulate its activity

To determine if CRTC3 activity is altered in human melanoma, we performed CRTC3 profiling on a cohort of 367 melanoma patients from the TCGA database. Alterations of CRTC3 were present in 10% of samples, most of which increased its expression; these were associated with reduced patient survival (Fig. 4e, Suppl. Fig. 4f). We evaluated the transcriptional signature of melanomas with alterations in CRTC3. To this end, we performed a co-expression analysis in the TCGA patient cohort and compared gene groups showing high unique positive correlation with each CRTC family member (Suppl. Table 5). In line with our molecular data, transcript signatures related to CRTC3 showed enrichment in pigmentation and cell migration (Fig. 4f, g, Suppl. Fig. 4h).

Based on public data^54,55^ we compiled a list of spontaneous CRTC3 mutations in melanoma and noticed that 20% of these occur in or near regulatory 14-3-3 binding sites (Suppl. Fig. 4g, Suppl. Table 6). Phosphorylation of conserved serines within SIK consensus motifs (LXBS/TX<u>S</u>XXXL) sequesters CRTC family in the cytoplasm by 14-3-3 binding; disruption of these sites would be expected to increase CRTC activity^24,56^. Based on their location on CRTC3 protein sequence we chose six patient mutations for further analysis in a CRE-luciferase reporter assay. Three of these mapped to 14-3-3 binding sites (L166F, P331H, S368F), one flanks a CRTC3 sumoylation site^57^ (P142S), while two are outside of discrete domains (H315N, P419S). Almost all mutant CRTC3 proteins stimulated reporter activity to a greater extent than WT, suggesting that CRTC3 is indeed upregulated in subsets of melanoma tumors (Fig. 4f).

## Discussion

CRTC family members share extensive sequence homology within their CREB-binding, regulatory and trans-activation domains. Correspondingly, loss of any one family member can be readily compensated by other family members, as we observed for MITF expression in primary melanoblasts and B16F1 cells. By contrast, OCA2 was significantly down-regulated following CRTC3 KO, accounting for the deficiencies in melanin synthesis without corresponding reductions in protein amounts of melanogenic enzymes. We found that the induction of OCA2 is mediated in part by recruitment of CREB/CRTC to CREB binding sites within a conserved enhancer. Knockout of CRTC3 reduced basal expression of OCA2, leading to lower net induction of pigmentation even in presence of high cAMP. Consistent with its substantial baseline activity, CRTC3 is the only CRTC family member that modulates constitutive pigmentation in mice.

By contrast with other CRTCs, CRTC3 also appears to stimulate CRE-dependent transcription in response to growth factors that activate ERK1/2. Phosphorylation of Ser^391^ primes CRTC3 for subsequent activation by modest increases in cAMP that are sufficient to dephosphorylate and fully activate CRTC3, but not CRTC1 or CRTC2. cAMP and ERK1/2 pathways function cooperatively in this setting, explaining in part the higher baseline activity of CRTC3 in melanocytes and melanoma cells. By contrast with the indiscriminant activation of all CRTCs in response to high cAMP levels, ERK1/2 and cAMP activate CRTC3 cooperatively under low cAMP signaling conditions to promote CREB-target gene expression (Suppl. Fig. 4i; Fig. 4i). In melanocytes and melanoma cells, several communication points between cAMP and MAPK/ERK cascades have been described, including transactivation of c-Kit by MC1R, activation of ERK1/2 by cAMP, inhibition of CRAF by PKA and regulation of MITF^17–20^. Our results suggest that CRTC3 functions as another communication point between these two pathways, thereby increasing specificity in signal transduction.

BRAF inhibitors represent a mainstay for the treatment of melanoma; reducing resistance to these inhibitors appears critical in improving patient survival. Knockout of CRTC3 in a BRAF-mutant melanoma model increased intracellular concentrations of cAMP as well as PKA activation and impaired chemotactic migration, invasion and anchorage-free growth. Conversely, up-regulation of CRTC3 was associated with improved cell viability during vemurafenib treatment.

Remarkably, spontaneous CRTC3 point mutants identified in a subset of human melanomas map to regulatory regions in the protein that normally sequester this coactivator in the cytoplasm. Our data suggest that the increased activity of CRTC3 in this setting contributes to melanoma progression by increasing the expression of CREB target genes involved in motility, invasiveness, or viability. The increased activity of these CRTC3 mutant proteins in melanomas may promote the constitutive upregulation of CREB target genes while avoiding potential inhibitory effects of PKA activation on tumor growth or metastasis. Future studies should provide further insight into this process.

## Author contributions

J.O. conceived the study, designed and performed experiments, analyzed data, and wrote the manuscript. T.S. performed IF experiments, analyzed data and provided reagents. N.N. provided experimental technical support under J.O. supervision. J.M.V. generated antisera for m-OCA2 (PBL #7431). M.S. provided suggestions and feedback for RNA-seq analysis and performed bioinformatics screen of the human intron 86 in HERC2 gene. M.M. conceived the study, provided feedback and reviewed/edited the manuscript.

## Acknowledgements

This work was supported by NIH grant R01 DK083834, the Leona M. and Harry B. Helmsley Charitable Trust, the Clayton Foundation for Medical Research and the Salkexcellerator Fund. We acknowledge the core facilities of the Salk Institute: Flow Cytometry Core Facility with funding from NIH-NCI CCSG: P30 014195 and Shared Instrumentation Grant S10-OD023689 (Aria Fusion cell sorter), the Waitt Advanced Biophotonics Core Facility with funding from NIH-NCI CCSG: P30 014195 and the Waitt Foundation and the NGS Core Facility of the Salk Institute with funding from NIH-NCI CCSG: P30 014195, the Chapman Foundation and the Helmsley Charitable Trust.

## Methods

### Animal studies

All animal procedures were approved by the Institutional Animal Care and Use Committee of the Salk Institute. Mice were housed in colony cages with a 12 hr light/12 hr dark cycle in a temperature-controlled environment. C57BL6 were purchased from Jackson Laboratories. Knock-outs of CRTC1, CRTC2 and CRTC3 were previously described^25,27,58^. iDCT-GFP mice were obtained from NCI Mouse Repository.

### Primary melanoblast isolation and melanocyte culture

Primary cells were isolated from whole epidermises of 2 days old WT, CRTC1 KO, CRTC2 KO and CRTC3 KO pups and cultured in differentiation media containing TPA (P1585, Sigma) and cholera toxin (C8052, Sigma), following previously described detailed protocol^59^. For skin melanoblast quantification, WT and CRTC3 KO mice were crossed with mice carrying iDCT:GFP transgene system. Pregnant females were given 1mg/ml doxycycline in water and epidermal cell suspensions were isolated from 2 days old pups (WT N=5, KO N=4). GFP positive cells were counted through fluorescence-activated cell sorting (Becton-Dickinson Influx™ cytometer). To account for differences in total amounts of cells isolated from each animal, quantification was expressed as percentage of GFP positive cells per epidermis. In experiments where XB2 feeder keratinocytes were used, cells were obtained from Wellcome Trust Functional Genomics Cell Bank and prepared using mitomycin c treatment (M4287, Sigma), as previously described^59^. In some experiments, catalase (LS001896, Worthington Biochemical) and an antioxidant supplement (A1345, Sigma) were added to the primary cultures in the attempt to improve survival and differentiation of CRTC3 KO melanoblasts.

### Cell line culture

B16F1, A375, MeWO and HEK293T cells were obtained from the American Type Culture Collection. Cells were tested for mycoplasma contamination with MycoAlert kit (LT07-418, Lonza) before use in experiments. A375, MeWO and HEK293T cells were consistently cultured in DMEM media (Gibco®, high glucose), supplemented with 10% fetal bovine serum (FBS, Gemini Bio-Products) in 5% CO_2_ at 37°C. Two different culture modes were tested for B16F1 in order to preserve their melanogenic potential. These were cultures in RPMI 1640 medium (Gibco®, high glucose) without phenol red supplemented with 10% FBS in 10% CO_2_ or DMEM without phenol red supplemented with 10% FBS in 5% CO_2_. In both cases cells were plated sparsely (1×10^4^) for regular maintenance. For induction of melanogenesis, cells were grown to 65% confluence, making sure to avoid cell clumping, and treated with compounds that stimulate melanin synthesis (Forskolin (FSK, F6886; Sigma) or IBMX (I5879, Sigma)). Cells grown in RPMI 1640 were switched to DMEM before stimulation. Maintenance plates were regularly tested (every 3 passages) to confirm the ability of cells to produce melanin. All cells were used for experiments between passages 4 and 12, after which they were discarded and a new low-passage batch thawed for the next experimental round. Under these conditions, there was no difference in melanin output of B16F1 cells between the two culture modes, so we used consistent DMEM for most shown results.

Silencing of CRTC3 in B16F1 cells was performed by transfecting cells with pSilencer 2.1-U6 puro plasmid (AM5762, Applied Biosystems) carrying CRTC3 shRNA constructs. 5’ phosphorylated double stranded CRTC3 oligonucleotides were cloned into BamHI and HindIII sites of the pSilencer 2.1-U6 puro plasmid, according to the manufacturer’s protocol. 0.8×10^6^ cells were transfected with 1.5µg plasmid DNA using Lipofectamine 2000 (11668019, Invitrogen). Selection with 2µg/ml puromycin (P8833, Sigma) was started 48h post transfection and maintained for at least 7 days before testing bulk cell populations for successful CRTC3 knock-down and melanin induction. Empty vector was used as a control.

Sequences of working constructs:

Sh1: 5’GATCCGCTTCAGCAACTGCGCCTTTTCAAGAGAAAGGCGCAGTTGCTGAAGTTTTTTGGAAA 3’

Sh3:5’GATCCGAAGCTCCTCTGGTCTCCATTCAAGAGATGGAGACCAGAGGAGCTTCTTTTTTGGAAA 3’

Sh4: 5’GATCCGCACATCAAGGTTTCAGCATTCAAGAGATGCTGAAACCTTGATGTGCTTTTTTGGAAA 3’

For CRISPR-Cas9 knockout of CRTC1 and CRTC3, B16F1 cells were separately transfected with two different guide RNAs (gRNAs) per gene, cloned into pSpCas9(BB)-2A-GFP (PX458) (Addgene plasmid # 48138)^60^. CTRL cells were transfected with the empty vector. Single clones were selected by fluorescence-activated cell sorting (Becton-Dickinson Influx™ cytometer) after 48 hours. The gRNAs target exons 1 and 3 of CRTC1 and exons 1 and 8 of CRTC3:

mC1g1: 5’ CACCGTGACAGGGGTCGACGGTGCG 3’ ex3

mC1g2: 5’ CACCGTCACCCGCGCGGCCCGCGTC 3’ ex1

mC3g1: 5’ CACCGCATCAAGCCGATAATGTTCG 3’ ex1

mC3g2: 5’ CACCGAGCCACTGCCTAAACACCTG 3’ ex8

Rescue experiments were performed by transfecting clonal B16F1 CRTC3 KO cells with pSelect-puro-mcs plasmid (psetp_mcs, Invivogen) carrying full length CRTC3, CRTC2, OCA2, CRTC3-Ser^391^A or CRTC2/3 hybrid constructs (CRTC2_328-449_ exchanged for CRTC3_326-402_, as previously decribed^48^), cloned into SalI and NheI sites. Transfected cells were selected with 2µg/ml puromycin for at least 7 days before initial testing. Empty vector was used to select for CTRL cells. Sequences of oligonucleotides used for cloning:

CRTC2 F: 5’ GATGGTCGACATGGCGACGTCAGGGGCGAACGGGCCGGGTTCC 3’

CRTC2 R: 5’ CATGCTAGCTCACTGTAGCCGATCACTACGGAATGAGTCCTC 3’

CRTC3 F: 5’ GATGGTCGACATGGCCGCCTCGCCCGGTTCGGGCAGCGCCAAC 3’

CRTC3 R: 5’ CATGCTAGCTCATAGCCGGTCAGCTCGAAACGTCTCCTCCAC 3’

OCA2 F: 5’ GATGGTCGACATGCGCCTAGAGAACAAAGACATCAGGC 3’

OCA2 R: 5’ CATGCTAGCTTAATTCCATCCCACCACAATGTGAGCAATCAGG 3’

Site directed mutagenesis of CRTC3-Ser391 to Ala residue was performed from pSelect-puro-mcs-CRTC3 construct with oligonucleotides:

S391A_F: 5’ GGCGCAGGCAGCCTCCAGTCGCCCCTCTCACGCTCTCTCCTGG 3’

S391A_R: 5’ CCAGGAGAGAGCGTGAGAGGGGCGACTGGAGGCTGCCTGCGCC 3’

Q5 high fidelity polymerase (M0492S, New England Biolabs) was used for the PCR reaction (95°C 3min, 95°C 30sec, 55°C 1min, 72°C for 7min, 18 cycles, 72°C for 7min final extension). Template plasmid was digested with DpnI for 2h at 37°C before amplification in Top10 competent cells (Invitrogen). Presence of S^391^A mutation was confirmed by sequencing.

CRTC3 knock-out and single cell selection in A375 cells was performed with the same CRISPR protocol used for B16F1 cells. The gRNAs targeted exons 1 and 11 of human CRTC3:

hC3g1: 5’ CACCGTGAGCGGCCCGTCTCGGCGT 3’ ex11

hC3g2: 5’ CACCGCGCGCTGCACACGCAGAGAC 3’ ex1

### Melanin quantification

1.5 µg of dorsal hair was plucked from 8 weeks old WT and CRTC3 KO animals (N=3 for each group) and solubilized in 1ml of 1M NaOH at 85°C for 4h under agitation. Samples were centrifuged at 12000 rpm for 5 minutes and the absorbance of supernatants was read at 475nm in the Synergy-H1 microplate reader (BioTek). Absorbance values were compared to the standard curve of synthetic melanin (155343, MP Biomedicals) dissolved in 1M NaOH. The standard curve was in a linear range of the experimental values.

For extracellular melanin quantification, B16F1 cells were plated in 6-well dishes at the density of 0.5⨯10^6^ cells/well in phenol red-free media and treated with FSK (F6886, Sigma) or IBMX (I5879, Sigma) for 60h, 24h after seeding. Final concentrations of small molecules used for treatment are indicated in figure legends. Melanin in the collected medium was determined by comparison of experimental absorbance values at 405nm with synthetic melanin standard curve diluted in the same media.

### Transmission Electron Microscopy (TEM)

B16F1 cells were fixed in a solution of 2% paraformaldehyde, 2.5% glutaraldehyde, and 2mM CaCl_2_ in 0.1M sodium cacodylate buffer (pH 7.4) for 2 h at room temperature, post-fixed in 1% osmium tetroxide for 40 and 1.5% potassium ferrocyanide in sodium cacodylate buffer for 1 hour at 4°C in the dark, stained in 1% aqueous uranyl acetate at 4°C in the dark, dehydrated in ethanol graded series, and embedded in Eponate12 resin (Ted Pella). Ultra-thin sections of 70 nm were obtained using a diamond knife (Diatome) in an ultramicrotome (Leica EM UC7) and placed on copper grids (300 mesh). Sections were imaged on Zeiss TEM Model Libra 120, operated at 120 kV and captured as 2048 ⨯2040 pixel tiffs using a Zemas camera system.

### Tyrosinase activity

B16F1 cells were plated in 6-well dishes at a density of 0.5⨯10^6^ cells/well and treated with FSK for 48h, 24h after seeding. Cells were lysed in 100 μl of 50 mM sodium phosphate buffer (pH 6.8) containing 1% Triton X-100 and 0.1 mM phenylmethylsulfonyl fluoride (PMSF, Thermo Fisher) and frozen at −80°C for 30 min. Thawed cellular extracts were centrifuged at 12,000g for 30 min at 4°C. For whole lysate activity, supernatants (80 μl) and 20 μl of 3,4-Dihydroxy-L-phenylalanine (L-DOPA, 2 mg/ml freshly dissolved in 50mM sodium phosphate buffer) were placed in a 96-well plate, and the absorbance at 492 nm was read every 10 min for 1 h at 37 °C using the Synergy-H1 plate reader. For in-gel activity, protein concentration of cell lysates was determined by micro BCA assay (Pierce), equal protein amounts were mixed with Laemmli sample buffer without β-mercaptoethanol and incubated at 37°C for 15 min with slight agitation. Proteins were resolved on 8% sodium-dodecyl-sulfate (SDS)-acrylamide gel by electrophoresis. Gels were equilibrated in 50 mM sodium phosphate buffer (pH 6.8) for 1h before adding 1mg/ml L-DOPA. Further incubation was monitored until clear colorimetric detection of tyrosinase activity. L-DOPA was obtained from Tocris Bioscience (3788).

### OCA2 antibody production

Female New Zealand white rabbits (10-12 weeks old; I.F.P.S. Inc., Norco, California, USA) were used for mOCA2 antibody production as previously described^48^. OCA2 synthetic peptide (Cys^27^ mOCA2(2-27)-NH_2_) was synthesized by RS Synthesis (Louisville, KY) and conjugated to maleimide activated Keyhole Limpet Hemocyanin (KLH) per manufacturer’s instructions (77610, Thermo Fisher). Antisera with highest titers against the synthetic peptides were tested for the ability to recognize endogenous OCA2 in B16F1 cell line and primary mouse melanocytes. Rabbit PBL #7431 anti-mOCA2 was purified using Cys^27^ mOCA2(2-27)-NH_2_ covalently attached to Sulfolink agarose (20401, Thermo Fisher). Validation of the purified antibody is available in Suppl. Fig. 2a-d.

### Soft agar, migration and invasion assays

Soft agar assays were carried out as previously described^61^. Briefly, 6-well dishes were coated with 0.5% noble agar in DMEM media and cells were seeded at 3×10^4^ cells/well in DMEM containing 0.3% noble agar and 10% FBS. Cells were allowed to form colonies for 28 days and were subsequently imaged on Zeiss VivaTome microscope. Diameters of imaged colonies were measured using ImageJ. For comparison between B16F1 KO cells carrying full length CRTC3 or CRTC3-S^391^A constructs, cells were subjected to initial 7 day selection with 2µg/ml puromycin and tested for comparable expression of CRTC3. Puromycin was also added to noble agar layers in these experiments. For migration and invasion assays, A375 cells were grown in serum-free DMEM media for 24h prior to seeding into 24-well permeable 8µM pore PET transwell inserts, with or without Matrigel coating (Corning 353097 and 354480, respectively). Inserts and companion plates were equilibrated in serum-free media at 37°C in 5% CO_2_ for at least 2h before starting the experiments. 3-7⨯10^4^ cells in serum-free media were added to the transwell inserts and 2% FBS was used as chemotactic agent in the bottom well. For certain experiments, cAMP elevating compounds were mixed with cells prior to seeding with concentrations indicated in Figures and Legends. Cells were allowed to migrate for 24h or invade for 72h. Transwell membranes containing migrating/invading cells were fixed in 3.7% paraformaldehyde for 2 min at RT, permeabilized in methanol for 15 min at RT and stained with a solution of 0.2% crystal violet/20%methanol for 20 min in the dark. Non-migrating cells were removed from the inside of membranes by swabbing with cotton tips before imaging on Zeiss VivaTome microscope. Cells were counted with ImageJ.

### cAMP Measurement

B16F1 or A375 cells were plated at 0.5⨯10^6^ cells/well in 6-well dishes and treated with cAMP inducing compounds for 15min, 24h after seeding. Different compounds and concentrations are indicated in Figures and Legends. Cellular cAMP levels were measured using an ELISA kit (581001, Cayman Chemical Company) according to manufacturer’s instructions.

### Immunofluorescence

HEK 293T and A375 cells were plated in Poly-D-Lysine coated glass bottom dishes (P35GC-0-10-C, MatTek Corporation). 24h post seeding, cells were fixed with 4% paraformaldehyde. After incubation with the primary hCRTC3(414-432) (PBL #7019) antibody, microscopy samples were incubated with secondary antibodies conjugated with Alexa Fluor®-568 (goat anti-rabbit). Counterstaining with DAPI (14285, Cayman Chemical Company) was performed before image acquisition (LSM 710, Zeiss).

### Real time qRT-PCR analysis

RNA was extracted from cultured cells, sorted primary melanoblasts or whole skin with TRIzol (15596026, Invitrogen)-chloroform. Skin samples were weighted, frozen in liquid nitrogen and equal amounts of frozen tissue were ground with mortar and pestle before extracting RNAs in TRIzol. RNA extracts were further purified and DNAse treated by using RNAeasy Qiagen columns (DNAse 79254, Qiagen). cDNA was synthesized from 400 ng (melanoblasts) or 1µg (cell lines and tissues) input RNA using Transcriptor first-strand cDNA synthesis kit (04897030001, Roche) according to the manufacturer’s instructions. Quantitative PCR was performed using LightCycler 480 SYBR green I master mix (04887352001, Roche) in a LightCycler 480 II (Roche). Relative mRNA levels were calculated using the 2−ΔΔCt method, normalized to L32. Primer sequences:

RPL32: F 5’ TCTGGTGAAGCCCAAGATCG 3’ R: 5’ CTCTGGGTTTCCGCCAGTT 3’

MITF: F 5’ GACTAAGTGGTCTGCGGTGT 3’ R: 5’ CTGGTAGTGACTGTATTCTA 3’

TYR: F 5’ TGGACAAAGACGACTACCACA 3’ R: 5’TTTCAGTCCCACTCTGTTTCC 3’

TYRP1: F: 5’ GGCATCAGGGGAAAAGCAGA 3’ R: 5’ GCTCAGATGAAAATACAGCAGTACC 3’

DCT: F: 5’ AACAGACACCAGACCCTGGA 3’ R: 5’ AAGTTTCCTGTGCATTTGCATGT 3’

PMEL: F: 5’ ACTGCCAGCTGGTTCTACAC 3’ R: 5’ CACCGTCTTGACCAGGAACA 3’

MLANA: F: 5’ GTGTTCCTCGGGGAAGGTGT 3’ R: 5’ CAGCAGTGACATAGGAGCGT 3’

OCA2: F: 5’ ATAGTGAGCAGGGAGGCTGT 3’ R: 5’ ACTGATGGGCCAGCAAAAGA 3’

COL11A1: F: 5’ CGATGGATTCCCGTTCGAGT 3’ R: 5’ GAGGCCTCGGTGGACATTAG 3’

S100A4: F: 5’ CCTCTCTCTTGGTCTGGTCTC 3’ R: 5’ GTCACCCTCTTTGCCTGAGT 3’

### Proliferation and viability assays

Cells were seeded in white 96-well plates with clear bottom (6005181, PerkinElmer) at 3500 cells/well and cell proliferation over 3-5 day periods was determined by measuring luminescence with CellTiter-Glo® assay kit (G7570, Promega), according to manufacturer’s instructions. For experiments measuring A375 cell viability after CRTC3 over-expression and treatment with inhibitors, 1×10^6^ cells (grown in 6-well dishes) were subjected to nucleofection with Amaxa® Cell Line Nucleofector® kit V (VCA-1003, Lonza), following the manufacturer’s protocol. Total of 2µg pSelect-puro-CRTC3 or empty vector were used per well. Cells were treated with 5µM vemurafenib (PLX4032, RG7204, S1267, Selleckchem) or 20µM cisplatin (S1166, Selleckchem) for 72h, 24h post nucleofection. Prior to experiments, cells were tested for sensitivity to vemurafenib and cisplatin (inhibitor range 0-20µM) and IC_50_ determined to be 0.0829µM and 6.99µM, respectively.

### Sub-cellular fractionation and western blotting

Cells were grown to 65% confluency on 10cm dishes, treated with compounds indicated and fractionated using Calbiochem® fractionation kit (539790), according to the manufacturer’s instructions. For some fractionation experiments a previously described protocol^62^ was used with some modifications. Briefly, cells were treated with indicated compounds for 20min, washed and scraped in cold phosphate-buffered saline (PBS), resuspended in buffer A (10mM HEPES-KOH pH 7.4, 1.5mM MgCl_2_, 10mM KCl, 0.2mM PMSF, protease inhibitor cocktail (P9599, Sigma), phosphatase inhibitors 2 and 3 (P5726 and P0044, Sigma), 0.1% NP-40) and allowed to swell on ice for 10min. Cell swelling was monitored through light microscope visualization. Cells were spun and supernatant used as cytosolic fraction. Nuclear proteins were extracted in high salt buffer C (20mM HEPES-KOH pH 7.9, 25% glycerol, 420mM NaCl, 1.5mM MgCl_2_, 0.2mM EDTA, 0.2mM PMSF, protease inhibitor cocktail, phosphatase inhibitors 2 and 3) for 20min on ice. Compounds used for cell treatments in these experiments were SCF (S9915, Sigma), HGF (H9661, Sigma), FSK (F6886; Sigma) and IBMX (I5879, Sigma). Whole cell or tissue extracts were prepared by collecting the samples in KB (Killer buffer, 2M urea, 4% sucrose, 5% SDS, 1mM EDTA, protease inhibitor cocktail, phosphatase inhibitors 2 and 3) and then passing through Qiashredder columns (79656, Qiagen). Tissue samples were weighted, frozen in liquid nitrogen and ground using mortar and pestle before lysis. Protein concentrations were determined with Pierce micro BCA protein assay kit (23235, Thermo Fisher). Samples were resolved by SDS-polyacrylamide gel electrophoresis and transferred onto PVDF membranes (IPVH00010, Millipore). Membranes were blocked in 5% milk before adding antibodies of interest. HyGlo HRP detection kit (E2500, Denville) was used to visualize proteins.

### Luciferase assays

OCA2 enhancer region was cloned from mouse genomic DNA into KpnI and XhoI sites of the pGL3 promoter vector (E1751, Promega). MITF proximal promoter was cloned into SacI and HindIII sites of the PGL4 promoter vector (E6651, Promega). EVX_2xCRE promoter cloned into PGL4^56^ was used to measure transcriptional activity of CRTC3 mutations found in melanoma patients. Mutations in CRE, E-box sites and CRTC3 were produced *via* site-directed mutagenesis. All constructs were sequenced to confirm the presence of mutations. Cells were seeded in 24-well dishes at 0.03×10^6^ cells/well and transfected with reporter constructs 24h post seeding, using Lipofectamine 2000 (11668019, Invitrogen) according to the manufacturer’s instructions. Total of 400ng of DNA was added in each well. In the case of CRTC3 melanoma mutations, EVX and pSelect_CRTC3 constructs were co-transfected at 200ng DNA each. Expression of CRTC3 constructs was visualized *via* western blotting and was used to normalize activity. Media was changed 24h after transfection. Various treatments that the cells were subjected to 48h after transfection are indicated in respective Figures and Legends. Cells were lysed in luciferase extraction buffer (25mM Gly-Gly, 15mM MgSO_4_, 4mM EGTA, 1mM DTT, 1% Triton X-100). 50μl of extract was added to 50µl of assay buffer (25mM Gly-Gly, 15mM MgSO_4_, 4mM EGTA, 15mM K_2_HPO_4_, pH 7.8, 2mM DTT, 2.5mM ATP) and 50µl of 0.1mM D-luciferin K^+^ salt before measuring luminescence in a GloMax® multi microplate reader (Promega).

Oligonucleotides used for cloning:

MITFprom_F: 5’CATGAGCTCAGACTCGGGTGCAAGATGAAG 3’

MITFprom_R: 5’GTGAAGCTTAGCAAGGTTTCAGGCAGCCCC 3’

OCA2Enh_F: 5’GATGGGTACCCAGTCCATTTCTGAATCAACAC 3’

OCA2Enh_R: 5’ CATCTCGAGCCAGTGAGACTCCTGGCTCCAAG 3’

Oligonucleotides used for site-directed mutagenesis:

MITFprom_CRE_F: 5’TATCTATGAAAAAAAGCATGAAATCAAGCCAGCAGGGAAACTGATATC 3’

OCA2Enh_CRE1_F: 5’ ATCCTGATGGTGATTACATGAAAAGAGTTTTTTTTTTCTG 3’

OCA2Enh_CRE2_F: 5’ TTATTTCCCCACCTTGGATGAAAAGCATTTTAAAGTTTAC 3’

OCA2Enh_EBOX_F: 5’ CTGAACCTTTGTATTAACAATTGATTTTTATCCTGATGGTGAT 3’

CRTC3-P142S_F: 5’ GAGCTGGCCACGGCAACAGTCTCCTTGGAAAGAAGAGAAGCA 3’

CRTC3-L166F_F: 5’ GGACCAATTCTGATTCTGCTTTTCACACGAGTGCTCTGAGCACC 3’

CRTC3-H315N_F: 5’ CAACCTTCCAGCTGCCATGACTAACCTGGGGATAAGAACCTCCTC 3’

CRTC3-P331H_F: 5’ CTCCAAAGTTCTCGAAGTAACCATTCCATCCAAGCCACACTCAGT 3’

CRTC3-S368F_F: 5’ CACCCCTCCCTCCGGCTCTTCTTCCTTAGCAACCCGTCTCTTTCC 3’

CRTC3-P419S_F: 5’ GACCAGCCCACTGAACCCGTATTCTGCCTCCCAGATGGTGACCTCA 3’

### Chromatin immunoprecipitation and sequencing (ChIP-seq)

ChIP was performed as previously described^63^. Briefly, cells were grown to 80% confluency in 150mm dishes (one per IP), treated with DMSO or 5µM FSK for 30min, fixed in 1% formaldehyde for 10min and quenched with 125mM glycine for 5min. Cells were washed and scraped in ice-cold phosphate-buffered saline (PBS) and then resuspended in buffer LB3 (10mM Tris-HCl (pH 8.0), 100mM NaCl, 1mM EDTA, 0.5mM EGTA, 0.1% Na-deoxycholate, 0.5% N-laurylsarcosine, protease inhibitor cocktail). Resuspended cells were mixed with 100mg glass beads and sonicated (Virtis Virsonic 100 sonicator) at power 9 for 9 pulses (20sec on, 1min off on ice). Sonication testing was performed before every ChIP experiment to determine the optimal shearing conditions. After sonication, 1% final Triton X-100 was added and the extract clarified by centrifugation at 17,000 × g for 5 min. Thirty microliters of protein A-agarose beads (20333, Thermo Scientific) were prepared for IP by washing in LB3 buffer, then washing twice with PBS/0.5% bovine serum albumin (BSA) and incubating with 5µg antibody in PBS/0.5% BSA for 3h at 4°C with rotation. For ChIP, beads were washed once in PBS/0.5% BSA and 500μl of clarified extract was added to antibody-coupled beads and incubated overnight at 4°C with rotation.

Beads were washed 3 times in 500μl wash buffer 1 (20mM Tris-HCl pH 7.4, 150mM NaCl, 2mM EDTA, 0.1% SDS, 1% Triton X-100) and three times in wash buffer 2 (20mM Tris-HCl (pH 7.4), 250mM LiCl, 1mM EDTA, 1% Triton X-100, 0.7% Na-deoxycholate). Elution was then performed by incubating beads in 50μl elution buffer 1 (Tris-EDTA, 1% SDS) for 15min at 50°C and 45μl elution buffer 2 (TE, 1% SDS, 300 mM NaCl) for 15min at 50°C with shaking. Both elutions were combined and cross-links reversed overnight at 65°C with shaking. Samples were incubated with RNase A for 1h at 37°C and proteinase K for 1h at 50°C. ChIP DNA was purified using Agencourt AMPure XP beads (A63881, Beckman Coulter) and DNA amounts determined by Qubit® 2.0 fluorometer (Invitrogen). All samples had more than 0.5ng/µl DNA, which was considered as the cutoff to proceed to library preparation. Libraries were prepared with NebNext® ChIP seq reagents for Illumina (E6240 and E7335, New England Biolabs), following the manufacturer’s protocol. High-throughput sequencing was performed on the HiSeq 2500 system (Illumina) at a run configuration of single read 50 bases. Image analysis and base calling were done with Illumina CASAVA-1.8.2.on HiSeq 2500 system and sequenced reads were quality-tested using FASTQC.

### ChIP-seq analysis

Reads were aligned to the mouse (mm10) genomes using STAR^64^. Tag directories of uniquely mapped reads were generated with HOMER^65^ and peaks were called and annotated using HOMER (findPeaks in “factor” mode for CREB, CRTC2, CRTC3, POLII and MITF and in “histone” mode for H3AcK27; annotatePeaks.pl, default parameters). Motif enrichment near ChIP-seq peaks was performed with HOMER using findMotifsGenome.pl. Overlapping peaks were visualized by generating BigWig files with HOMER (makeMultiWigHub.pl, default parameters) and uploading the hubs in the UCSC Genome Browser (http://genome.ucsc.edu). Enrichment analysis was preformed using WebGestalt^66^.

### DNA binding motif screen in intron 86 of H.sapiens HERC2 gene

HOMER annotatePeaks.pl was used to scan for CRE and Mbox motifs in intron 86 of the HERC2 gene in the hg38 human genome. Motifs of interest were visualized in the UCSC genome browser and overlapped with variants from gnomAD^67^, PhyloP^68^ conservation tracks.

### RNA-sequencing and analysis

RNAs were prepared as indicated in RT-qPCR protocol. The quality of the isolated total RNA was assessed using the Tape Station 4200 and RNA-seq libraries were prepared using the TruSeq stranded mRNA Sample Preparation Kit v2 (RS-122-2001) according to Illumina protocols. RNA-seq libraries were multiplexed, normalized and pooled for sequencing. High-throughput sequencing was performed on the HiSeq 2500 system (Illumina) at a run configuration of single read 50 bases. Image analysis and base calling were done with Illumina CASAVA-1.8.2.on HiSeq 2500 system and sequenced reads were quality-tested using FASTQC. FASTQ files were aligned to the mouse mm10 or human hg19 genome builds using STAR^64^. Independent biological replicates (2 for CTRL and CRTC3 KO B16F1 and A375 cell line samples, 1 for CRTC3 rescue sample in B16F1 cells, 4 for WT and CRTC3 KO skin samples) were used for differential expression analysis with HOMER (analyzeRepeats.pl with option –raw; getDiffExpression.pl using DESeq2^69^). Expression was compared between vehicle and FSK treatment or CTRL and KO genomes and differentially expressed genes were defined as having Log_2_ fold change of ≥1 or ≤-1 (1.3 and -1.3 for whole skin samples) and an adjusted p-value of ≤0.05. Rescued genes were defined with a Log_2_ fold change of ≥0.58 or ≤-0.58 with respect to their relative KO values. Enrichment analysis was preformed using WebGestalt^66^. Heat maps were generated with Cluster 3.0^70^ and Java TreeView (version 1.1.6r4)^71^.

### Data availability

Sequencing data sets are available under GEO accession number GSE154117.

### Antibodies

Rabbit anti-CREB serum (244, in-house), rabbit anti-CRTC1 (C71D11, 2587, CST), rabbit anti-CRTC2 serum (6865, in-house), rabbit anti-CRTC3 (C35G4, 2720, CST), rabbit anti-CRTC3-pSer^391^ (PBL #7408, in-house^48^), rabbit-anti-hCRTC3 (PBL #7019, in-house^48^), rabbit anti-mOCA2 (PBL #7431, in-house, this work), rabbit anti-H3AcK27 (ab4729, Abcam), Anti-RNA polymerase II (CTD repeat YSPTSPS (phospho S2) antibody - ChIP Grade (ab5095)), mouse anti-Tubulin (05-829; EMD Millipore), mouse anti-MITF (clone C5, MAB3747-1 Millipore; MITF ChIP grade ab12039 Abcam), rabbit anti-TYR (ab61284, Abcam), rabbit anti-DCT (ab74073, Abcam), rabbit anti-MLANA (NBP254568H, Novus), rabbit anti-PMEL (ab137078, Abcam) rabbit anti-ERK1/2 (4695, CST), rabbit anti-pERK1/2 (9101, CST), mouse anti-HSP90α/β (SC-13119, Santa Cruz Biotechnology), rabbit anti-Histone H3 (9715, CST).

## References

1. Lin, J. Y. & Fisher, D. E. Melanocyte biology and skin pigmentation. Nature 445, 843–850 (2007).

2. Scherer, D. & Kumar, R. Genetics of pigmentation in skin cancer — A review. Mutat. Res. Mutat. Res. 705, 141–153 (2010).

3. Shannan, B., Perego, M., Somasundaram, R. & Herlyn, M. Heterogeneity in Melanoma. Cancer Treat. Res. 167, 1–15 (2016).

4. Vandamme, N. & Berx, G. Melanoma Cells Revive an Embryonic Transcriptional Network to Dictate Phenotypic Heterogeneity. Front. Oncol. 4, (2014).

5. Tsoi, J. et al. Multi-stage differentiation defines melanoma subtypes with differential vulnerability to drug-induced iron-dependent oxidative stress. Cancer Cell 33, 890-904.e5 (2018).

6. Wolf Horrell, E. M., Boulanger, M. C. & D’Orazio, J. A. Melanocortin 1 Receptor: Structure, Function, and Regulation. Front. Genet. 7, (2016).

7. Sánchez-Más, J., Hahmann, C., Gerritsen, I., García-Borrón, J. C. & Jiménez-Cervantes, C. Agonist-independent, high constitutive activity of the human melanocortin 1 receptor. Pigment Cell Res. 17, 386–395 (2004).

8. Cui, R. et al. Central role of p53 in the suntan response and pathologic hyperpigmentation. Cell 128, 853–864 (2007).

9. Fajardo, A. M., Piazza, G. A. & Tinsley, H. N. The Role of Cyclic Nucleotide Signaling Pathways in Cancer: Targets for Prevention and Treatment. Cancers 6, 436–458 (2014).

10. Levy, C., Khaled, M. & Fisher, D. E. MITF: master regulator of melanocyte development and melanoma oncogene. Trends Mol. Med. 12, 406–414 (2006).

11. Buscà, R. & Ballotti, R. Cyclic AMP a key messenger in the regulation of skin pigmentation. Pigment Cell Res. 13, 60–69 (2000).

12. Bertolotto, C. et al. Microphthalmia Gene Product as a Signal Transducer in cAMP-Induced Differentiation of Melanocytes. J. Cell Biol. 142, 827–835 (1998).

13. Wellbrock, C. & Arozarena, I. Microphthalmia-associated transcription factor in melanoma development and MAP-kinase pathway targeted therapy. Pigment Cell Melanoma Res. 28, 390–406 (2015).

14. Price, E. R. et al. Lineage-specific Signaling in Melanocytes c-Kit stimulation recruits p300/CBP to microphthalmia. J. Biol. Chem. 273, 17983–17986 (1998).

15. Hemesath, T. J., Price, E. R., Takemoto, C., Badalian, T. & Fisher, D. E. MAP kinase links the transcription factor Microphthalmia to c-Kit signalling in melanocytes. Nature 391, 298–301 (1998).

16. Wu, M. et al. c-Kit triggers dual phosphorylations, which couple activation and degradation of the essential melanocyte factor Mi. Genes Dev. 14, 301–312 (2000).

17. Herraiz, C. et al. Signaling from the human melanocortin 1 receptor to ERK1 and ERK2 mitogen-activated protein kinases involves transactivation of cKIT. Mol. Endocrinol. Baltim. Md 25, 138–156 (2011).

18. Buscà, R. et al. Ras mediates the cAMP-dependent activation of extracellular signal-regulated kinases (ERKs) in melanocytes. EMBO J. 19, 2900–2910 (2000).

19. Niwano, T., Terazawa, S., Nakajima, H. & Imokawa, G. The stem cell factor-stimulated melanogenesis in human melanocytes can be abrogated by interrupting the phosphorylation of MSK1: evidence for involvement of the p38/MSK1/CREB/MITF axis. Arch. Dermatol. Res. 310, 187–196 (2018).

20. Dumaz, N. et al. In melanoma, RAS mutations are accompanied by switching signaling from BRAF to CRAF and disrupted cyclic AMP signaling. Cancer Res. 66, 9483–9491 (2006).

21. Wellbrock, C. & Arozarena, I. The Complexity of the ERK/MAP-Kinase Pathway and the Treatment of Melanoma Skin Cancer. Front. Cell Dev. Biol. 4, 33 (2016).

22. Kozar, I., Margue, C., Rothengatter, S., Haan, C. & Kreis, S. Many ways to resistance: How melanoma cells evade targeted therapies. Biochim. Biophys. Acta BBA - Rev. Cancer 1871, 313–322 (2019).

23. Johannessen, C. M. et al. A melanocyte lineage program confers resistance to MAP kinase pathway inhibition. Nature 504, 138–142 (2013).

24. Altarejos, J. Y. & Montminy, M. CREB and the CRTC co-activators: sensors for hormonal and metabolic signals. Nat. Rev. Mol. Cell Biol. 12, 141–151 (2011).

25. Altarejos, J. Y. et al. The Creb1 coactivator Crtc1 is required for energy balance and fertility. Nat. Med. 14, 1112–1117 (2008).

26. Blanchet, E. et al. Feedback inhibition of CREB signaling promotes beta cell dysfunction in insulin resistance. Cell Rep. 10, 1149–1157 (2015).

27. Song, Y. et al. CRTC3 links catecholamine signalling to energy balance. Nature 468, 933–939 (2010).

28. MacKenzie, K. F. et al. PGE(2) induces macrophage IL-10 production and a regulatory-like phenotype via a protein kinase A-SIK-CRTC3 pathway. J. Immunol. Baltim. Md 1950 190, 565–577 (2013).

29. Bang, S. et al. Novel regulation of melanogenesis by adiponectin via the AMPK/CRTC pathway. Pigment Cell Melanoma Res. 30, 553–557 (2017).

30. Yun, C.-Y. et al. Nuclear Entry of CRTC1 as Druggable Target of Acquired Pigmentary Disorder. Theranostics 9, 646–660 (2019).

31. Kim, Y.-H. et al. Therapeutic Potential of Rottlerin for Skin Hyperpigmentary Disorders by Inhibiting the Transcriptional Activity of CREB-Regulated Transcription Coactivators. J. Invest. Dermatol. 139, 2359-2367.e2 (2019).

32. Kim, J.-H. et al. JNK suppresses melanogenesis by interfering with CREB-regulated transcription coactivator 3-dependent MITF expression. Theranostics 10, 4017–4029 (2020).

33. Bellono, N. W., Escobar, I. E., Lefkovith, A. J., Marks, M. S. & Oancea, E. An intracellular anion channel critical for pigmentation. eLife 3, e04543 (2014).

34. Park, S. et al. Unrevealing the role of P-protein on melanosome biology and structure, using siRNA-mediated down regulation of OCA2. Mol. Cell. Biochem. 403, 61–71 (2015).

35. Jannot, A.-S. et al. Allele variations in the OCA2 gene (pink-eyed-dilution locus) are associated with genetic susceptibility to melanoma. Eur. J. Hum. Genet. 13, 913–920 (2005).

36. Sturm, R. A. et al. A Single SNP in an Evolutionary Conserved Region within Intron 86 of the HERC2 Gene Determines Human Blue-Brown Eye Color. Am. J. Hum. Genet. 82, 424–431 (2008).

37. Duffy, D. L. et al. Multiple Pigmentation Gene Polymorphisms Account for a Substantial Proportion of Risk of Cutaneous Malignant Melanoma. J. Invest. Dermatol. 130, 520–528 (2010).

38. Donnelly, M. P. et al. A global view of the OCA2-HERC2 region and pigmentation. Hum. Genet. 131, 683–696 (2012).

39. Hawkes, J. E. et al. Report of a Novel OCA2 Gene Mutation and an Investigation of Two OCA2 Variants on Melanoma Predisposition in a Familial Melanoma Pedigree. J. Dermatol. Sci. 69, 30–37 (2013).

40. Steel, K. P., Davidson, D. R. & Jackson, I. J. TRP-2/DT, a new early melanoblast marker, shows that steel growth factor (c-kit ligand) is a survival factor. Dev. Camb. Engl. 115, 1111–1119 (1992).

41. Zaidi, M. R., Hornyak, T. J. & Merlino, G. A genetically engineered mouse model with inducible GFP expression in melanocytes. Pigment Cell Melanoma Res. 24, 393–394 (2011).

42. Guyonneau, L., Murisier, F., Rossier, A., Moulin, A. & Beermann, F. Melanocytes and pigmentation are affected in dopachrome tautomerase knockout mice. Mol. Cell. Biol. 24, 3396–3403 (2004).

43. Bennett, D. C. Mechanisms of differentiation in melanoma cells and melanocytes. Environ. Health Perspect. 80, 49–59 (1989).

44. Watabe, H. et al. Regulation of Tyrosinase Processing and Trafficking by Organellar pH and by Proteasome Activity. J. Biol. Chem. 279, 7971–7981 (2004).

45. Visser, M., Kayser, M. & Palstra, R.-J. HERC2 rs12913832 modulates human pigmentation by attenuating chromatin-loop formation between a long-range enhancer and the OCA2 promoter. Genome Res. 22, 446–455 (2012).

46. Visser, M., Kayser, M., Grosveld, F. & Palstra, R.-J. Genetic variation in regulatory DNA elements: the case of OCA2 transcriptional regulation. Pigment Cell Melanoma Res. 27, 169–177 (2014).

47. Q, L. et al. Mechanism of CREB recognition and coactivation by the CREB-regulated transcriptional coactivator CRTC2. Proc. Natl. Acad. Sci. U. S. A. 109, 20865–20870 (2012).

48. Sonntag, T. et al. Mitogenic Signals Stimulate the CREB Coactivator CRTC3 through PP2A Recruitment. iScience 11, 134–145 (2019).

49. Cancer Genome Atlas Network. Genomic Classification of Cutaneous Melanoma. Cell 161, 1681–1696 (2015).

50. Delyon, J. et al. PDE4D promotes FAK-mediated cell invasion in BRAF-mutated melanoma. Oncogene 36, 3252–3262 (2017).

51. Marquette, A., André, J., Bagot, M., Bensussan, A. & Dumaz, N. ERK and PDE4 cooperate to induce RAF isoform switching in melanoma. Nat. Struct. Mol. Biol. 18, 584–591 (2011).

52. Watanabe, Y. et al. Phosphodiesterase 4 regulates the migration of B16-F10 melanoma cells. Exp. Ther. Med. 4, 205–210 (2012).

53. Howe, A. K. Regulation of actin-based cell migration by cAMP/PKA. Biochim. Biophys. Acta 1692, 159–174 (2004).

54. Gonzalez-Perez, A. et al. IntOGen-mutations identifies cancer drivers across tumor types. Nat. Methods 10, 1081–1082 (2013).

55. Tate, J. G. et al. COSMIC: the Catalogue Of Somatic Mutations In Cancer. Nucleic Acids Res. 47, D941–D947 (2019).

56. Sonntag, T. et al. Analysis of a cAMP regulated coactivator family reveals an alternative phosphorylation motif for AMPK family members. PLoS ONE 12, (2017).

57. Hendriks, I. A. et al. Site-specific mapping of the human SUMO proteome reveals co-modification with phosphorylation. Nat. Struct. Mol. Biol. 24, 325–336 (2017).

58. Wang, Y. et al. Targeted disruption of the CREB coactivator Crtc2 increases insulin sensitivity. Proc. Natl. Acad. Sci. U. S. A. 107, 3087–3092 (2010).

59. Godwin, L. S. et al. Isolation, culture, and transfection of melanocytes. Curr. Protoc. Cell Biol. 63, 1.8.1-20 (2014).

60. Ran, F. A. et al. Genome engineering using the CRISPR-Cas9 system. Nat. Protoc. 8, 2281–2308 (2013).

61. Borowicz, S. et al. The Soft Agar Colony Formation Assay. JoVE J. Vis. Exp. e51998 (2014) doi:10.3791/51998.

62. Andrews, N. C. & Faller, D. V. A rapid micropreparation technique for extraction of DNA-binding proteins from limiting numbers of mammalian cells. Nucleic Acids Res. 19, 2499 (1991).

63. Van de Velde, S. et al. CREB Promotes Beta Cell Gene Expression by Targeting Its Coactivators to Tissue-Specific Enhancers. Mol. Cell. Biol. 39, (2019).

64. Dobin, A. et al. STAR: ultrafast universal RNA-seq aligner. Bioinforma. Oxf. Engl. 29, 15–21 (2013).

65. Heinz, S. et al. Simple combinations of lineage-determining transcription factors prime cis-regulatory elements required for macrophage and B cell identities. Mol. Cell 38, 576–589 (2010).

66. Liao, Y., Wang, J., Jaehnig, E. J., Shi, Z. & Zhang, B. WebGestalt 2019: gene set analysis toolkit with revamped UIs and APIs. Nucleic Acids Res. 47, W199–W205 (2019).

67. Collins, R. L. et al. A structural variation reference for medical and population genetics. Nature 581, 444–451 (2020).

68. Pollard, K. S., Hubisz, M. J., Rosenbloom, K. R. & Siepel, A. Detection of nonneutral substitution rates on mammalian phylogenies. Genome Res. 20, 110–121 (2010).

69. Love, M. I., Huber, W. & Anders, S. Moderated estimation of fold change and dispersion for RNA-seq data with DESeq2. Genome Biol. 15, 550 (2014).

70. Eisen, M. B., Spellman, P. T., Brown, P. O. & Botstein, D. Cluster analysis and display of genome-wide expression patterns. Proc. Natl. Acad. Sci. 95, 14863–14868 (1998).

71. Saldanha, A. J. Java Treeview--extensible visualization of microarray data. Bioinforma. Oxf. Engl. 20, 3246–3248 (2004).

