## Supplementary Information for "Transcriptional co-activator regulates melanocyte differentiation and oncogenesis by integrating cAMP and MAPK/ERK pathways"

### Supplementary Figure Legends

**Supplementary Fig. 1: CRTC3 regulates pigmentation in mouse melanocytes and melanoma cells.**

**a)** Western blot showing CRTC expression levels in primary melanocytes from WT animals. **b)** Percentage of DCT:GFP-positive cells in WT and CRTC3 KO newborn animals (P2) isolated from skin *via* cell sorting. **c)** Quantitative PCR data from primary melanocytes of WT and CRTC3 KO animals differentiated for 14 days, showing enrichment for markers of melanocytes and fibroblasts. **d)** Melanin production and western blot in B16F1 cells transfected with different short hairpins directed against CRTC3. Cells were treated with 5µM FSK for 60h **e)** Melanin production and western blot in B16F1 cells transfected with control or CRTC3 CRISPR KO guides (KO1, KO2). Cells were selected as single clones and treated with 5µM FSK for 60h. **f)** Western blot showing basal and FSK-stimulated accumulation of MITF and Tyrosinase in CTRL and CRTC3 KO B16F1 cells (5µM FSK, 4h). **g)** Quantification of early and late stage melanosomes in TEM analyses of CTRL and CRTC3 KO B16F1 cells treated with 5µM FSK for 48h. N=10 cells per genotype. **h)** Enriched clusters for genes associated with CRTC3 peaks in ChIP-seq studies of B16F1 cells. **i)** Enrichment of motifs under CRTC3 peaks in ChIP-seq analysis of B16F1 cells treated with 5µM FSK for 1h. **j)** Enriched clusters for differentially downregulated genes in RNA-seq experiments of CTRL and CRTC3 KO B16F1 cells. **k)** Heat map of significant differentially expressed genes in RNA-sequencing experiments of B16F1 cells with indicated genotypes and treatments (5µM FSK, 1h).

**Supplementary Fig. 2: Regulation of OCA2 by CRTC family members.**

**a)** Binding titers of purified OCA2 antibody (see Methods for details). **b-d)** Validation of OCA2 antibody: b) Western blot showing sub-cellular fractionation of OCA2 in B16F1 cells treated with 5µM FSK for 24h. HSP90, MLANA and Histone H3 were used as controls for cytoplasmic, melanosomal and nuclear fractions, respectively. **c)** Expression of OCA2 in primary melanocytes isolated from skins of newborn WT animals (P2). **d)** Induction of OCA2 in B16F1 cells of indicated genotypes after treatment with 5µM FSK for 24h. **e)** Western blot and melanogenesis assay in CTRL, CRTC1 KO and CRTC3 KO B16F1 cells, treated with 5µM FSK O/N for western analyses and 60h for melanin production.

**Supplementary Fig. 3: CRTC3 is regulated by cAMP and MAPK cascades in melanoma cells.**

**a)** Sub-cellular fractionation of B16F1 cells treated with 100µM IBMX or 20ng/ml HGF for 1h showing localization of CRTC3-pSer$^{391}$. **b)** Extracellular melanin in CTRL and CRTC3 KO B16F1 cells transfected with empty vector, full length CRTC3 or CRTC3-S391A mutant constructs where indicated. Quantification was performed on 4 independent experiments and significance determined by one-way ANOVA and Tukey's multiple comparisons tests. Western blot insert shows representative expression of CRTC3 in samples used. Cells were treated with 1µM FSK for 60h. **c)** Quantification of CRTC nuclear localization from Fig. 3d **d)** CRTC3 KO analysis of several A375 clones transfected with control or CRTC3 KO CRISPR guides. Red rectangles show cell clones selected for further experiments. **e)** Five day proliferation assay of CTRL and CRTC3 KO A375 cells. **f)** Quantification of colony size of CTRL and CRTC3 KO A375 cells grown in soft agar for 28 days. Significance determined by Welch's t-test. **g)** Three day proliferation test of CTRL and CRTC3 KO B16F1 cells. **h)** Quantification of colony size of CTRL, CRTC3 KO and CRTC3 rescued cells grown in soft agar for 21 days. Significance determined by one-way ANOVA and Tukey's multiple comparisons tests. **i)** Quantification and representative images of colony size of CRTC3 KO B16F1 cells transfected with full length CRTC3 or CRTC3-S391A mutant constructs, grown in soft agar for 28 days. Transfected cells were selected with 2µg/ml puromycin for 2 weeks prior to the assay. Bar is 0.2mm

**Supplementary Fig. 4: Modulation of PKA and CRTC3 activity in human melanoma.**

**a)** Western blots showing activation of PKA in control and CRTC3 KO A375 cells. **b,c)** Representative images of migration (24h) of A375 cells treated with increasing concentrations of FSK, IBMX (100µM) or PKA inhibitor H89 (10µM), where indicated. Bar is 100µm in b) and 50µm in c). **d)** Viability of CTRL and CRTC3 KO A375 cells treated with 5µM vemurafenib or 20µM cisplatin for 72h. **e)** Measurement of A375 cell viability after over-expression of full length CRTC3 construct and treatment with vemurafenib (5µM) or cisplatin (20µM) for 72h. Western blot insert shows expression and $S^{391}$ phosphorylation of CRTC3. Bar graph represents data from 4 replicate measurements and significance determined by Welch's t-test. **f)** Graphical representation of CRTC3 genomic alterations in a melanoma patient cohort from TCGA, Firehose Legacy N=367. **g)** Enriched processes for genes that showed unique positive expression correlation with CRTC1 and CRTC2 in the TCGA, Firehose Legacy cohort of melanoma patients (N=367). Data was obtained from cbioportal.org. Genes are listed in Suppl. Table 5. **h)** Sequence alignment of human and mouse CRTC3 protein. CREB binding domain (1-50), PP2A binding domain (326-402) and transactivation domain (548-619) are highlighted in yellow, light gray and teal, respectively. Phosphorylated serines within 14-3-3 binding sites are highlighted in purple (full consensus LXBS/TX<u>S</u>XXXL). Three 14-3-3 binding sites and the sumo site containing tested human melanoma mutations are shown in solid and dashed rectangles, respectively. Missense melanoma mutations listed in Suppl. Table 6 are underlined on the sequences in bold red. **i)** Model for joint regulation of CRTC3 by cAMP and ERK1/2 in high cAMP state.

### Supplementary Figures

**Supplementary Fig. 1:** CRTC3 regulates pigmentation in mouse melanocytes and melanoma cells.

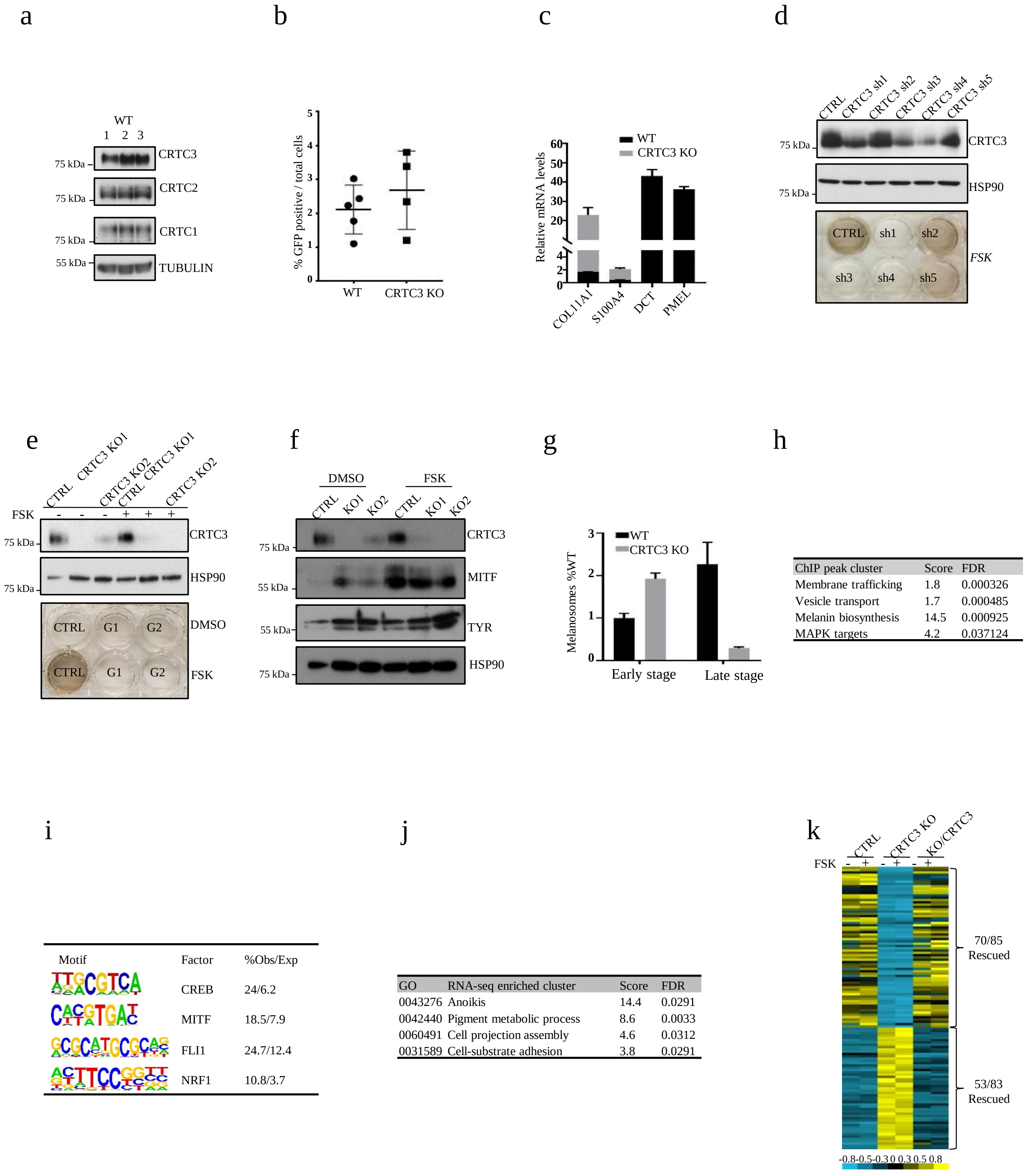

| ChIP peak cluster | Score | FDR |
|---|---|---|
| Membrane trafficking | 1.8 | 0.000326 |
| Vesicle transport | 1.7 | 0.000485 |
| Melanin biosynthesis | 14.5 | 0.000925 |
| MAPK targets | 4.2 | 0.037124 |

| Motif | Factor | %Obs/Exp |
|---|---|---|
| | CREB | 24/6.2 |
| | MITF | 18.5/7.9 |
| | FLI1 | 24.7/12.4 |
| | NRF1 | 10.8/3.7 |

| GO | RNA-seq enriched cluster | Score | FDR |
|---|---|---|---|
| 0043276 | Anoikis | 14.4 | 0.0291 |
| 0042440 | Pigment metabolic process | 8.6 | 0.0033 |
| 0060491 | Cell projection assembly | 4.6 | 0.0312 |
| 0031589 | Cell-substrate adhesion | 3.8 | 0.0291 |

**Supplementary Fig. 2:** Regulation of OCA2 by CRTC family members.

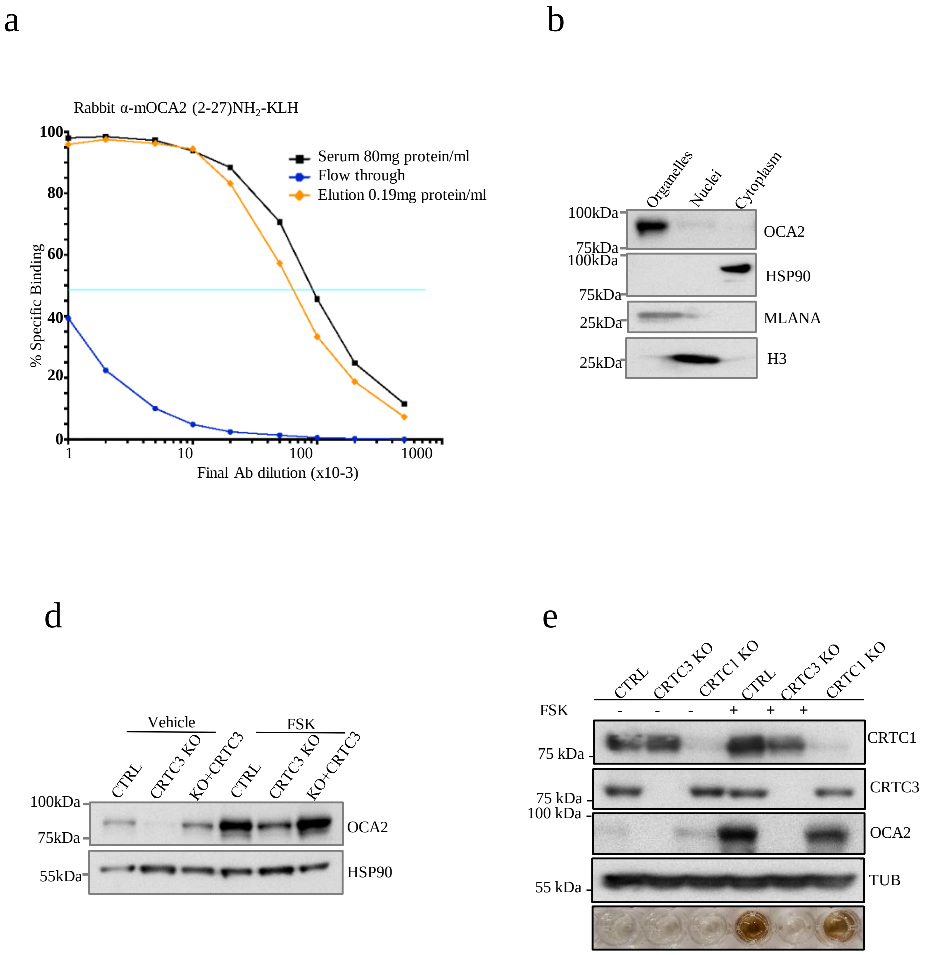

**Supplementary Fig. 3:** CRTC3 is regulated by cAMP and MAPK cascades in melanoma cells.

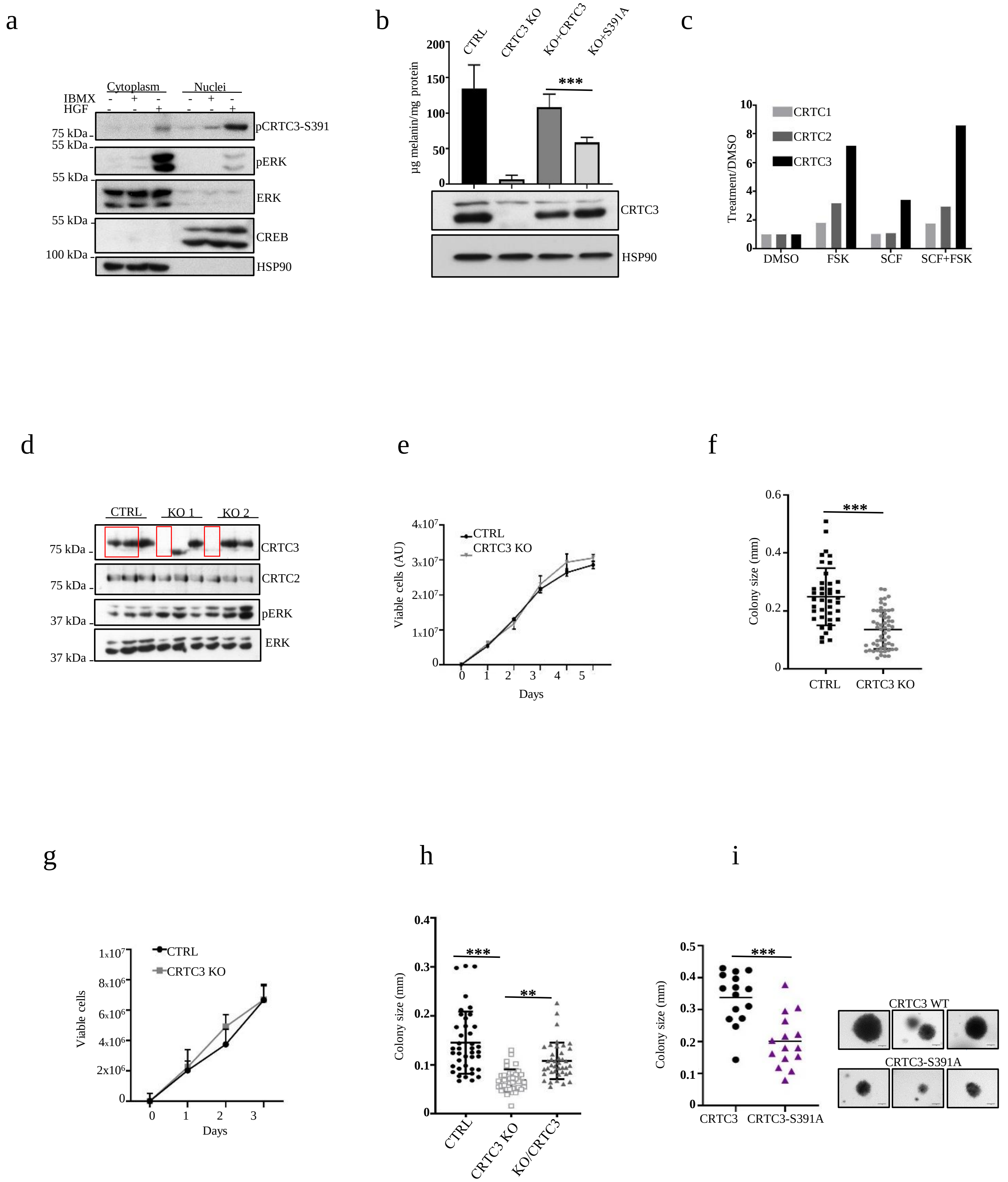

**Supplementary Fig. 4:** Modulation of PKA and CRTC3 activity in human melanoma.

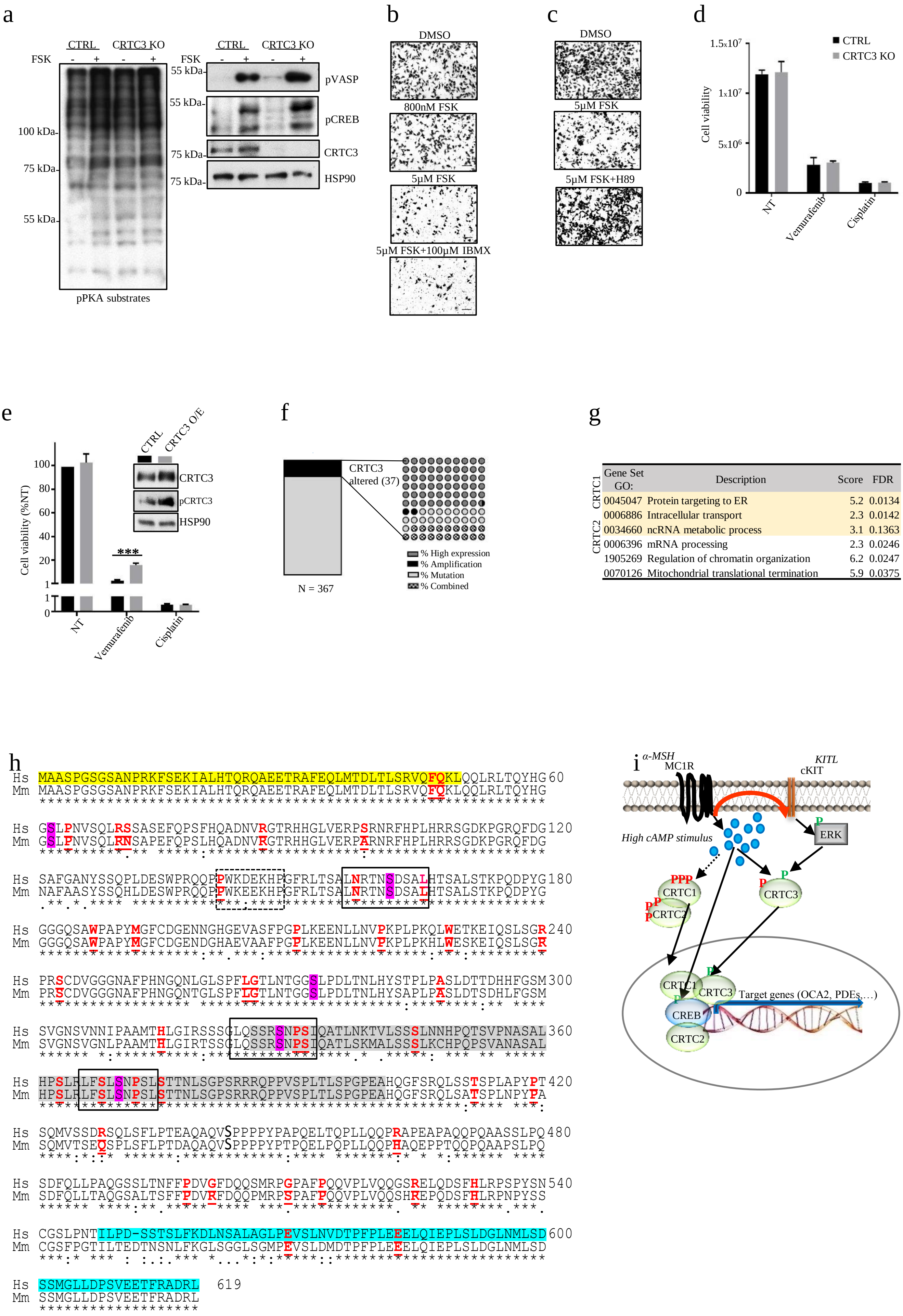

### Supplementary Tables

**Suppl. Table 1. Differentially expressed genes in WT vs. CRTC3 KO whole skin ($Log_2$+/-1.3, adj.p value<=0.05)**

| Transcript ID | Gene name | WT counts | KO counts | KO vs. WT Log2 Fold Change | KO vs. WT adj. p-value |
|---|---|---|---|---|---|
| NM_021879 | Oca2 | 142.9325 | 6.5225 | -4.53846 | 4.10E-19 |
| NM_027828 | Fam110c | 214.395 | 66.275 | -1.71051 | 2.34E-15 |
| NM_021882 | Pmel | 2559.2925 | 941.23 | -1.48433 | 5.45E-11 |
| NM_032400 | Sucnr1 | 106.7175 | 12.7975 | -3.0712 | 1.88E-09 |
| NM_028696 | Nabp1 | 995.1525 | 377.45 | -1.39958 | 8.93E-08 |
| NR_102304 | D830026I12Rik | 6.2725 | 70.115 | 3.499586 | 1.71E-07 |
| NM_144823 | Acsl6 | 235.3325 | 45.6375 | -2.35045 | 6.62E-07 |
| NM_080458 | Bsnd | 101.635 | 13.1825 | -2.92954 | 9.92E-07 |
| NM_144557 | Myrip | 26.66 | 84.8625 | 1.659308 | 9.92E-07 |
| NM_145363 | Tspan10 | 96.0625 | 26.7525 | -1.91508 | 4.06E-06 |
| NM_145963 | Kcnj14 | 47.105 | 6.73 | -2.8838 | 1.04E-05 |
| NM_001101531 | Gm5538 | 173.69 | 27.9125 | -2.67854 | 1.05E-05 |
| NM_177596 | Zfp947 | 78.485 | 25.94 | -1.5876 | 1.08E-05 |
| NM_026178 | Mmd | 4180.9025 | 1517.7125 | -1.49371 | 1.74E-05 |
| NM_153803 | Glb1l2 | 1827.39 | 635.455 | -1.52695 | 2.83E-05 |
| NM_144512 | Slc6a13 | 226.795 | 80.71 | -1.50558 | 4.04E-05 |
| NM_001001445 | Trpv1 | 35.645 | 6.39 | -2.48343 | 6.41E-05 |
| NM_029541 | Arxes1 | 473.68 | 151.17 | -1.68435 | 7.12E-05 |
| NM_026189 | Eepd1 | 811.055 | 332.175 | -1.31774 | 7.36E-05 |
| NM_172152 | Slc24a4 | 61.25 | 14.3725 | -2.18345 | 9.98E-05 |
| NM_001145403 | Kcnt1 | 17.5675 | 76.92 | 2.108982 | 0.000163 |
| NM_008759 | Sebox | 203.865 | 62.7075 | -1.68048 | 0.000252 |
| NM_146187 | Ffar2 | 534.2075 | 115.3675 | -2.29556 | 0.000263 |
| NM_144936 | Tmem45b | 487.6 | 99.9525 | -2.28557 | 0.000263 |
| NM_017394 | Slc7a10 | 1252.195 | 514.3125 | -1.31344 | 0.000299 |
| NM_172610 | Mpped1 | 37.92 | 107.0625 | 1.465559 | 0.000299 |
| NM_030599 | Klrb1b | 27.5325 | 77.0675 | 1.489472 | 0.000309 |
| NM_028235 | Ttc30b | 651.3 | 233.935 | -1.47953 | 0.000333 |
| NM_011395 | Slc22a3 | 233.7025 | 62.1375 | -1.955 | 0.000361 |
| NM_026956 | Cd209f | 224.395 | 54.725 | -2.06114 | 0.000542 |
| NM_008340 | Igfals | 110.63 | 31.435 | -1.86749 | 0.000674 |

| | | | | | |
|---|---|---|---|---|---|
| NM_016685 | Comp | 23.1775 | 60.945 | 1.384731 | 0.000674 |
| NM_030166 | Galnt15 | 125.3425 | 473.39 | 1.923421 | 0.000885 |
| NM_001204959 | Retn | 4916.08 | 1566.8625 | -1.66775 | 0.00101 |
| NM_007606 | Car3 | 79246.328 | 26081.193 | -1.61828 | 0.001486 |
| NM_207281 | Mettl21e | 45.0075 | 196.6125 | 2.144766 | 0.001726 |
| NM_011648 | Tshr | 908.165 | 290.205 | -1.69422 | 0.001756 |
| NM_029803 | Ifi27l2a | 1245.66 | 481.6525 | -1.37821 | 0.001842 |
| NM_018732 | Scn3a | 88.365 | 231.275 | 1.374242 | 0.001842 |
| NM_018752 | Trpm1 | 732.115 | 196.77 | -1.9804 | 0.001957 |
| NM_001085536 | Gm13178 | 166.2125 | 43.5425 | -1.92159 | 0.002142 |
| NM_008812 | Padi2 | 50.4825 | 164.7725 | 1.70792 | 0.003108 |
| NM_177746 | Awat2 | 215.4125 | 76.8675 | -1.47874 | 0.003545 |
| NM_183296 | Krtap16-3 | 343.8525 | 13.765 | -4.62918 | 0.003728 |
| NM_052994 | Spock2 | 137.435 | 451.4025 | 1.713977 | 0.004152 |
| NM_028048 | Slc25a35 | 183.3 | 71.2175 | -1.36726 | 0.004298 |
| NM_175370 | Als2cr12 | 58.69 | 22.1275 | -1.39172 | 0.004438 |
| NM_028765 | Acoxl | 45.78 | 18.73 | -1.34264 | 0.004541 |
| NM_207539 | Mrgprb8 | 40.2775 | 124.2725 | 1.605188 | 0.0046 |
| NM_007873 | Doc2b | 16.1575 | 58.36 | 1.803372 | 0.004647 |
| NM_008100 | Gcg | 17.7875 | 2.995 | -2.55592 | 0.004851 |
| NM_010571 | Irs3 | 123.98 | 35.125 | -1.8507 | 0.00497 |
| NM_007639 | Cd1d1 | 802.85 | 316.4775 | -1.38049 | 0.005231 |
| NR_130996 | 4930511A08Rik | 22.78 | 5.985 | -1.96497 | 0.00533 |
| NM_133675 | Sptssb | 31.945 | 7.08 | -2.13312 | 0.005635 |
| NR_131249 | Linc-md1 | 5.9125 | 24.28 | 2.022583 | 0.005635 |
| NM_010923 | Nnat | 1396.03 | 198.845 | -2.83201 | 0.005815 |
| NM_001013767 | Capn11 | 4.7125 | 51.9675 | 3.515012 | 0.005946 |
| NM_008365 | Il18r1 | 15.0775 | 52.1425 | 1.764076 | 0.006514 |
| NM_079835 | Btnl2 | 34.235 | 7.3725 | -2.23977 | 0.006767 |
| NM_139142 | Slc6a20a | 8.6975 | 35.07 | 2.011288 | 0.006979 |
| NM_013809 | Cyp2g1 | 1089.03 | 375.475 | -1.58484 | 0.00804 |
| NM_026212 | Agpat2 | 2863.8025 | 1099.69 | -1.39193 | 0.009028 |
| NM_133681 | Tspan1 | 28.4775 | 7.4675 | -1.93103 | 0.009053 |
| NM_017371 | Hpx | 49.15 | 16.3025 | -1.61365 | 0.009372 |
| NM_001085504 | Gm436 | 56.8525 | 14.6625 | -1.93046 | 0.010046 |
| NM_001114084 | Dgat2l6 | 79.22 | 22.365 | -1.78635 | 0.011468 |
| NM_028010 | Apol6 | 440.55 | 119.8575 | -1.93011 | 0.011652 |
| NM_001085376 | Pappa2 | 720.3 | 1829.955 | 1.34277 | 0.011907 |
| NM_001168590 | 2010106E10Rik | 6.4 | 32.0525 | 2.323899 | 0.01218 |
| NM_017370 | Hp | 7543.495 | 1277.175 | -2.56861 | 0.012669 |
| NR_003960 | Gm5478 | 9.695 | 114.095 | 3.598682 | 0.013659 |

| | | | | | |
|---|---|---|---|---|---|
| NM_013479 | Bcl2l10 | 28.4325 | 7.5625 | -1.87708 | 0.01388 |
| NM_029008 | Lvrn | 190.4025 | 502.0275 | 1.396776 | 0.01388 |
| NM_178258 | Il22ra2 | 19.225 | 99.88 | 2.410792 | 0.01388 |
| NM_198414 | Paqr9 | 64.8075 | 21.3575 | -1.64241 | 0.014439 |
| NM_008424 | Kcne1 | 49.4825 | 19.9 | -1.35108 | 0.014577 |
| NM_001048176 | Cerkl | 3.5925 | 15.9925 | 2.09607 | 0.015275 |
| NR_045153 | E130008D07Rik | 11.355 | 37.045 | 1.709522 | 0.015359 |
| NM_010484 | Slc6a4 | 413.325 | 1207.595 | 1.543159 | 0.015574 |
| NM_181748 | Ffar4 | 54.29 | 20.5275 | -1.45516 | 0.019157 |
| NM_008205 | H2-M9 | 3.1475 | 14.8175 | 2.261366 | 0.020056 |
| NR_015388 | Dlx6os1 | 3.755 | 31.195 | 3.078546 | 0.020862 |
| NM_175510 | Unc80 | 4.945 | 21.0625 | 2.136928 | 0.022752 |
| NM_027853 | Mettl7b | 49.665 | 16.45 | -1.55559 | 0.024475 |
| NM_011470 | Sprr2d | 72.205 | 1152.2425 | 4.022831 | 0.024505 |
| NM_012044 | Pla2g2e | 64.31 | 21.8025 | -1.57862 | 0.024853 |
| NM_011066 | Per2 | 280.6925 | 860.9675 | 1.616997 | 0.025501 |
| NM_001160370 | A830018L16Rik | 23.305 | 8.0075 | -1.60301 | 0.027062 |
| NM_001037294 | Alpk2 | 57.7725 | 154.9325 | 1.41974 | 0.027859 |
| NM_029006 | Kcnk16 | 23.9175 | 5.9875 | -2.0364 | 0.028571 |
| NR_037994 | 2310043L19Rik | 11.9625 | 36.2825 | 1.630536 | 0.029993 |
| NM_013821 | Hsd3b6 | 196.63 | 69.7775 | -1.54758 | 0.030683 |
| NM_001159367 | Per1 | 707.6575 | 2009.5175 | 1.508842 | 0.031127 |
| NM_146241 | Trhde | 80.44 | 32.5525 | -1.33191 | 0.031593 |
| NM_011990 | Slc7a11 | 178.35 | 567.5625 | 1.696168 | 0.0322 |
| NM_011430 | Sncg | 799.36 | 127.7275 | -2.61424 | 0.032411 |
| NM_021351 | Cryba4 | 248.52 | 85.2925 | -1.55881 | 0.032411 |
| NM_011994 | Abcd2 | 2803.9575 | 1052.3075 | -1.45212 | 0.032737 |
| NM_018790 | Arc | 29.905 | 91.5825 | 1.608313 | 0.037091 |
| NM_001302356 | Cbln2 | 14.25 | 51.635 | 1.817889 | 0.037091 |
| NM_144943 | Cd207 | 85.7225 | 8.1075 | -3.40254 | 0.037285 |
| NM_010115 | Egfbp2 | 10.2275 | 54.825 | 2.457697 | 0.038588 |
| NM_010644 | Klk1b26 | 10.2275 | 54.57 | 2.450885 | 0.038884 |
| NM_183187 | Fam107a | 26.455 | 226.2925 | 3.122422 | 0.039726 |
| NR_040624 | D030045P18Rik | 4.0925 | 19.715 | 2.313564 | 0.040675 |
| NM_172708 | Dok7 | 25.045 | 77.1475 | 1.617558 | 0.041101 |
| NM_001166625 | Ccr9 | 9.865 | 42.325 | 2.099585 | 0.041718 |
| NM_021443 | Ccl8 | 111.5525 | 35.31 | -1.69271 | 0.042144 |
| NM_177793 | Mettl24 | 58.12 | 19.35 | -1.59369 | 0.042177 |
| NM_001201323 | Krt83 | 1430.0525 | 487.15 | -1.57837 | 0.042895 |
| NM_010450 | Hoxa11 | 4.215 | 18.995 | 2.088094 | 0.04314 |
| NM_010672 | Krtap6-1 | 373.7025 | 43.0675 | -3.09517 | 0.043316 |

| | | | | | |
|---|---|---|---|---|---|
| NR_033520 | Tmem181b-ps | 55.025 | 218.4 | 1.864671 | 0.043316 |
| NM_013713 | Krtap15 | 1064.955 | 61.0225 | -4.09804 | 0.044173 |
| NM_009265 | Sprr1b | 375.9325 | 2733.635 | 2.888567 | 0.046257 |
| NM_183264 | Tespa1 | 21.5325 | 58.1725 | 1.418316 | 0.046333 |
| NM_011724 | Xirp1 | 352.4225 | 1016.5375 | 1.542709 | 0.047232 |

**Suppl. Table 2. FSK responsive genes in CTRL vs. CRTC3 KO B16F1 cells ($Log_2$+/-1, adj.p value<=0.05)**

| Transcript ID | Gene name | CTRL_FSK vs. CTRL_DMSO Log2 Fold Change | CTRL_FSK vs. CTRL_DMSO adj. p-value | KO_FSK vs. CTRL_FSK Log2 Fold Change | KO_FSK vs. CTRL_FSK adj. p-value |
|---|---|---|---|---|---|
| NM_015743 | Nr4a3 | 3.204768 | 1.06E-55 | -1.95131 | 2.16E-32 |
| NM_008601 | Mitf | 2.441533 | 3.89E-51 | -0.70662 | 3.54E-05 |
| NM_008037 | Fosl2 | 4.361325 | 6.36E-46 | -1.68476 | 3.53E-07 |
| NM_010831 | Sik1 | 3.8861 | 1.86E-20 | -1.634 | 1.43E-09 |
| NM_020507 | Tob2 | 1.90416 | 2.91E-18 | -1.0687 | 2.58E-08 |
| NM_013613 | Nr4a2 | 4.47046 | 3.31E-18 | -2.45009 | 0.010839 |
| NM_010444 | Nr4a1 | 2.57457 | 1.46E-10 | -2.15197 | 5.61E-10 |
| NM_001042591 | Arrdc3 | 1.931189 | 2.21E-10 | -0.56714 | 0.22317 |
| NM_009621 | Adamts1 | -1.83734 | 3.18E-08 | 0.871405 | 4.49E-05 |
| NM_011267 | Rgs16 | -3.0385 | 1.40E-07 | 1.496431 | 0.164562 |
| NM_010234 | Fos | 3.030366 | 3.06E-06 | -0.42033 | 0.662456 |
| NM_001159367 | Per1 | 1.515418 | 0.000136 | -0.40977 | 0.477377 |
| NM_019764 | Amotl2 | -1.04575 | 0.004942 | 0.455256 | 0.306505 |
| NM_001039710 | Coq10b | 1.398869 | 0.023987 | -0.35693 | 0.526408 |
| NM_009061 | Rgs2 | 3.216122 | 0.029613 | 0.202597 | 0.940741 |
| NM_001290726 | Arid5a | 1.471639 | 0.033072 | -0.34939 | 0.687407 |

**Suppl. Table 3. Differentially expressed and rescued genes in CTRL vs. CRTC3 KO B16F1 cells ($Log_2$+/-1, adj.p value<=0.05)**

| Transcript ID | Gene name | KO_FSK vs. CTRL_FSK Log2 Fold Change | KO_FSK vs. CTRL_FSK adj. p-value | RESCUE_FSK vs. KO_FSK Log2 Fold Change | RESCUE_FSK vs. KO_FSK adj. p-value |
|---|---|---|---|---|---|
| NM_001253754 | Gpm6a | -3.3 | 1.71E-41 | 2.1 | 1.11E-06 |
| NM_023627 | Isyna1 | -7.9 | 9.88E-34 | 7.5 | 1.78E-06 |
| NM_015743 | Nr4a3 | -2.0 | 2.16E-32 | 2.3 | 4.84E-35 |
| NM_010288 | Gja1 | -3.7 | 4.08E-23 | 5.0 | 5.21E-44 |

| | | | | | |
|---|---|---|---|---|---|
| NM_018760 | Slc4a4 | -4.7 | 9.12E-19 | 2.6 | 0.037061 |
| NM_134160 | Mcoln3 | 1.9 | 9.12E-19 | -1.6 | 7.29E-11 |
| NM_026139 | Armcx2 | -1.9 | 4.14E-18 | 2.3 | 7.02E-17 |
| NM_015774 | Ero1l | 1.4 | 9.99E-15 | -0.8 | 0.000855 |
| NM_001001979 | Megf10 | 2.3 | 1.33E-14 | -1.1 | 0.002477 |
| NM_198111 | Akap6 | 2.2 | 3.73E-13 | -1.2 | 2.06E-05 |
| NM_021879 | Oca2 | -2.4 | 1.51E-11 | 1.4 | 0.00038 |
| NM_001040397 | Filip1l | 1.7 | 1.62E-11 | -1.3 | 1.11E-05 |
| NM_010444 | Nr4a1 | -2.2 | 5.61E-10 | 3.1 | 7.09E-46 |
| NM_010831 | Sik1 | -1.6 | 1.43E-09 | 2.0 | 7.95E-20 |
| NM_001122993 | B3galt5 | 2.7 | 2.24E-08 | -1.3 | 0.000187 |
| NM_020507 | Tob2 | -1.1 | 2.58E-08 | 0.9 | 0.000124 |
| NM_001190449 | Ddah2 | -5.0 | 3.22E-08 | 4.6 | 0.000103 |
| NM_010404 | Hap1 | -1.2 | 3.51E-08 | 0.8 | 0.017395 |
| NM_001081413 | Unc13b | -3.4 | 1.41E-07 | 4.3 | 1.74E-14 |
| NM_177606 | Plekhh2 | -4.8 | 1.65E-07 | 6.2 | 0.042895 |
| NM_008037 | Fosl2 | -1.7 | 3.53E-07 | 2.2 | 0.008113 |
| NM_010807 | Marcksl1 | -1.7 | 4.18E-07 | 1.7 | 3.89E-10 |
| NM_016851 | Irf6 | 3.6 | 4.90E-07 | -2.0 | 0.585652 |
| NM_001302205 | Lmo1 | -1.8 | 1.13E-06 | 0.9 | 0.256015 |
| NM_001243064 | Cav1 | -2.3 | 1.83E-06 | 3.2 | 3.65E-10 |
| NM_146168 | Vopp1 | -3.7 | 2.00E-06 | 4.9 | 8.82E-11 |
| NM_001252292 | Mest | -1.6 | 2.51E-06 | 2.1 | 3.42E-11 |
| NM_199029 | Zfp395 | 2.2 | 8.92E-06 | -1.4 | 0.038363 |
| NM_001102613 | Phldb3 | 1.2 | 1.00E-05 | -0.8 | 0.028897 |
| NM_028945 | Fam208a | 1.1 | 1.30E-05 | -1.0 | 0.000835 |
| NM_008726 | Nppb | 3.1 | 1.30E-05 | -1.7 | 0.002308 |
| NM_144923 | Blvrb | -1.5 | 1.36E-05 | 1.2 | 0.019878 |
| NM_013778 | Akr1c13 | -1.9 | 1.89E-05 | 0.7 | 0.641026 |
| NM_198249 | Arhgef40 | 1.1 | 1.89E-05 | -0.7 | 0.078709 |
| NM_011123 | Plp1 | 1.7 | 2.44E-05 | -2.7 | 2.99E-22 |
| NM_008478 | L1cam | 1.2 | 2.82E-05 | -1.7 | 9.48E-11 |
| NM_007868 | Dmd | -2.7 | 2.98E-05 | 3.0 | 0.00083 |
| NM_011350 | Sema4f | 1.4 | 3.98E-05 | -1.0 | 0.013616 |
| NM_001037221 | Samd4 | 1.2 | 4.71E-05 | -0.8 | 0.021819 |
| NM_134013 | Psme4 | 1.1 | 5.56E-05 | -1.3 | 5.27E-12 |
| NM_183162 | Helz2 | 1.2 | 6.44E-05 | -1.2 | 0.002824 |
| NM_010577 | Itga5 | -1.7 | 8.80E-05 | 1.9 | 0.001922 |
| NM_001033214 | E330034G19Rik | 2.1 | 0.000107 | -0.8 | 0.404693 |
| NM_010576 | Itga4 | 1.2 | 0.000139 | -0.7 | 0.002918 |
| NM_011789 | Apc2 | -4.5 | 0.000152 | 1.6 | 0.910553 |

| | | | | | |
|---|---|---|---|---|---|
| NM_025959 | Psmc6 | 1.0 | 0.000154 | -0.8 | 0.000656 |
| NM_001168333 | Tinagl1 | -2.8 | 0.000154 | 5.4 | 9.75E-08 |
| NM_007488 | Arnt2 | 3.3 | 0.000161 | -1.9 | 0.028997 |
| NM_008168 | Grik5 | -3.1 | 0.000256 | 2.3 | 0.228784 |
| NM_001025576 | Ccdc141 | -1.5 | 0.000264 | 0.7 | 0.528394 |
| NM_177354 | Vash1 | 1.6 | 0.000271 | -2.0 | 3.64E-06 |
| NM_011670 | Uchl1 | -1.6 | 0.000545 | 2.1 | 1.97E-09 |
| NM_183115 | Ccdc125 | -4.4 | 0.000771 | 3.8 | 0.148773 |
| NM_001347162 | Srl | -1.5 | 0.000906 | 1.3 | 0.031554 |
| NM_013749 | Tnfrsf12a | 1.2 | 0.000906 | -0.4 | 0.616455 |
| NM_175514 | Fam171b | -2.9 | 0.00121 | 3.2 | 0.004833 |
| NM_001310453 | Mapk8 | 1.0 | 0.001318 | -1.0 | 0.007148 |
| NM_008795 | Cdk18 | 1.1 | 0.001542 | -0.7 | 0.184849 |
| NM_028266 | Col16a1 | 1.3 | 0.00187 | -1.8 | 6.72E-06 |
| NM_027711 | Iqgap2 | -1.7 | 0.001892 | 2.3 | 5.36E-10 |
| NM_133922 | Krba1 | -1.9 | 0.002425 | 2.1 | 0.007032 |
| NM_001159417 | Irf9 | 1.4 | 0.002469 | -0.8 | 0.260903 |
| NM_010153 | Erbb3 | 1.2 | 0.002493 | -0.6 | 0.044699 |
| NM_029609 | Lhpp | -1.2 | 0.002598 | 1.1 | 0.039371 |
| NM_001005426 | Zcwpw1 | 1.8 | 0.002639 | -2.2 | 0.000706 |
| NM_001310664 | Art3 | 2.4 | 0.00337 | -1.1 | 0.062514 |
| NM_011864 | Papss2 | -4.5 | 0.003506 | 2.7 | 0.635449 |
| NM_027399 | Steap1 | -2.7 | 0.003854 | 3.4 | 0.000585 |
| NM_001113246 | Chn1 | -1.9 | 0.004059 | 1.3 | 0.215339 |
| NM_019421 | Cd320 | -1.1 | 0.004119 | 0.9 | 0.102279 |
| NM_175521 | Nyap1 | -2.5 | 0.004319 | 1.0 | 0.789155 |
| NM_019634 | Tspan7 | -1.0 | 0.004319 | 1.1 | 0.019251 |
| NM_177235 | Bend6 | 1.4 | 0.004416 | -1.1 | 3.30E-06 |
| NM_009177 | St3gal1 | 1.1 | 0.004694 | -1.1 | 4.09E-06 |
| NM_139269 | Pla2g16 | -3.1 | 0.004753 | 3.3 | 0.002139 |
| NM_013529 | Gfpt2 | 1.7 | 0.004989 | -0.8 | 0.488313 |
| NM_001316688 | Slc27a3 | 1.6 | 0.005903 | -1.9 | 0.004267 |
| NM_022995 | Pmepa1 | 1.5 | 0.00598 | -2.7 | 1.42E-05 |
| NM_008872 | Plat | 1.9 | 0.00598 | -3.1 | 4.81E-06 |
| NM_008047 | Fstl1 | -4.3 | 0.006613 | 2.6 | 0.589709 |
| NM_026910 | Tnik | 1.1 | 0.007599 | -0.8 | 0.050631 |
| NM_027406 | Aldh1l1 | -1.4 | 0.007599 | 0.9 | 0.22188 |
| NM_001159647 | Cntn1 | -3.2 | 0.008341 | 3.6 | 0.030394 |
| NM_148941 | Elovl4 | -1.2 | 0.008512 | 1.6 | 0.003196 |
| NM_001271623 | Gja3 | 1.1 | 0.008964 | -4.2 | 1.32E-17 |
| NM_025907 | Mettl6 | 1.0 | 0.010016 | -0.7 | 0.243316 |

| | | | | | |
|---|---|---|---|---|---|
| NM_013613 | Nr4a2 | -2.5 | 0.010839 | 2.9 | 0.005367 |
| NM_001025251 | Mbp | 1.8 | 0.011444 | -2.8 | 3.66E-05 |
| NM_001103156 | Steap2 | -1.6 | 0.011913 | 2.6 | 2.02E-07 |
| NM_008179 | Gspt2 | -2.4 | 0.012286 | 2.1 | 0.152692 |
| NM_023118 | Dab2 | -1.4 | 0.012607 | 1.5 | 0.003286 |
| NR_029382 | Mir17hg | 1.3 | 0.013033 | -1.1 | 0.079648 |
| NM_008548 | Man1a | -1.6 | 0.013444 | 2.7 | 3.48E-06 |
| NM_053201 | Magee1 | -1.3 | 0.014008 | 1.7 | 0.004622 |
| NM_026767 | Dpm3 | -1.3 | 0.014323 | 0.8 | 0.528896 |
| NM_010284 | Ghr | -1.5 | 0.014375 | 1.2 | 0.280922 |
| NM_010770 | Matn3 | 2.6 | 0.015007 | -2.7 | 0.053474 |
| NM_019410 | Pfn2 | -1.3 | 0.017911 | 1.6 | 6.68E-05 |
| NM_001281848 | Kcnk2 | 3.1 | 0.018321 | -1.1 | 0.693495 |
| NM_011348 | Sema3e | -2.3 | 0.019293 | 1.6 | 0.371509 |
| NM_013724 | Nrk | -3.2 | 0.020305 | 2.7 | 0.358513 |
| NM_001004365 | Actr3b | -2.2 | 0.020305 | 2.4 | 0.006756 |
| NM_008813 | Enpp1 | -3.8 | 0.021096 | 4.5 | 0.006389 |
| NR_030696 | 9930014A18 Rik | -2.4 | 0.021615 | 3.5 | 9.74E-06 |
| NM_134189 | Galnt10 | -1.4 | 0.021978 | 1.4 | 0.009153 |
| NM_178704 | Dpy19l3 | -1.7 | 0.022488 | 2.1 | 0.007148 |
| NM_026336 | 2310057J18 Rik | 1.7 | 0.024572 | -3.6 | 2.63E-05 |
| NM_134250 | Havcr2 | -2.2 | 0.024775 | 1.2 | 0.676478 |
| NM_145523 | Gca | -1.8 | 0.024904 | 2.0 | 0.042007 |
| NM_027641 | Spef1 | 1.4 | 0.027401 | -1.0 | 0.180902 |
| NM_010142 | Ephb2 | 2.4 | 0.029113 | -2.0 | 0.010429 |
| NM_053143 | Pcdhb18 | 2.1 | 0.031752 | -1.9 | 0.015801 |
| NM_175501 | Adamts12 | -1.1 | 0.032349 | 0.8 | 0.17033 |
| NM_029011 | Pyroxd2 | 1.3 | 0.032895 | -1.2 | 0.167539 |
| NM_023422 | Hist1h2bc | 1.1 | 0.033419 | -0.7 | 0.47695 |
| NM_145612 | Zfp810 | 1.4 | 0.03406 | -0.8 | 0.446213 |
| NM_183029 | Igf2bp2 | -3.3 | 0.034743 | 1.2 | 0.923358 |
| NM_171824 | Pgbd5 | 1.1 | 0.037872 | -1.7 | 0.005077 |
| NM_027185 | Def6 | -2.6 | 0.037875 | 1.2 | 0.81661 |
| NM_008609 | Mmp15 | 1.2 | 0.038863 | -0.9 | 0.180902 |
| NM_001162926 | Fam84b | -1.2 | 0.044037 | 1.9 | 6.36E-05 |
| NM_028716 | Phf19 | -1.8 | 0.045423 | 2.9 | 5.76E-05 |
| NM_009932 | Col4a2 | -2.0 | 0.045423 | 2.4 | 0.007334 |

**Suppl. Table 4. Differentially expressed genes in CTRL vs. CRTC3 KO A375 cells ($Log_2$+/-1, adj.p value<=0.05)**

| Transcript ID | Gene name | CTRL_ DMSO counts | CTRL_ FSK counts | CRTC3KO_ DMSO counts | CRTC3KO_ FSK counts | CRTC3KO_ FSK vs. CTRL_FSK Log2 Fold Change | CRTC3KO_ FSK vs. CTRL_FSK adj. p-value |
|---|---|---|---|---|---|---|---|
| NM_182899 | CREB5 | 628 | 598 | 45 | 38 | -4.0 | 1.33E-40 |
| NM_001823 | CKB | 95 | 137 | 896 | 966 | 2.8 | 8.84E-34 |
| NM_003246 | THBS1 | 44240 | 44474 | 21749 | 17210 | -1.4 | 3.65E-33 |
| NM_173200 | NR4A3 | 39 | 715 | 42 | 132 | -2.5 | 1.62E-25 |
| NM_021214 | ABHD17C | 1799 | 2438 | 715 | 863 | -1.5 | 9.74E-25 |
| NM_001037340 | PDE4B | 324 | 531 | 70 | 77 | -2.8 | 6.40E-22 |
| NM_001197222 | PDE4D | 1083 | 3418 | 742 | 1318 | -1.4 | 7.78E-21 |
| NM_017831 | RNF125 | 211 | 200 | 777 | 794 | 2.0 | 3.11E-20 |
| NM_201432 | GAS7 | 1182 | 1199 | 3964 | 4085 | 1.7 | 8.83E-19 |
| NM_022551 | RPS18 | 5650 | 5546 | 11359 | 11779 | 1.0 | 1.55E-17 |
| NM_001293298 | CEMIP | 5203 | 5668 | 1997 | 2052 | -1.5 | 2.93E-15 |
| NM_014268 | MAPRE2 | 690 | 720 | 1753 | 1763 | 1.3 | 3.00E-14 |
| NM_001103184 | FMN1 | 2209 | 2193 | 977 | 899 | -1.3 | 6.05E-14 |
| NM_022842 | CDCP1 | 240 | 246 | 10 | 15 | -4.1 | 6.22E-14 |
| NM_001005340 | GPNMB | 849 | 827 | 300 | 292 | -1.5 | 1.53E-13 |
| NM_024652 | LRRK1 | 325 | 363 | 67 | 72 | -2.4 | 1.53E-13 |
| NM_003155 | STC1 | 524 | 1157 | 286 | 405 | -1.6 | 1.53E-13 |
| NM_007085 | FSTL1 | 452 | 436 | 113 | 103 | -2.1 | 1.84E-13 |
| NM_002587 | PCDH1 | 392 | 375 | 1339 | 1525 | 2.0 | 1.90E-13 |
| NM_002923 | RGS2 | 1036 | 2006 | 693 | 917 | -1.2 | 1.90E-13 |
| NM_012428 | NPTN | 2337 | 2560 | 1283 | 1269 | -1.1 | 5.29E-13 |
| NM_006472 | TXNIP | 140 | 134 | 674 | 640 | 2.2 | 7.08E-12 |
| NR_104309 | ULK3 | 1687 | 1688 | 827 | 805 | -1.1 | 1.44E-11 |
| NM_004811 | LPXN | 334 | 372 | 991 | 999 | 1.4 | 1.60E-11 |
| NM_012304 | FBXL7 | 5 | 4 | 196 | 179 | 5.3 | 1.40E-10 |
| NM_015476 | TPGS2 | 986 | 952 | 2019 | 1975 | 1.0 | 1.77E-10 |
| NM_001320644 | DMBT1 | 40 | 224 | 16 | 33 | -2.8 | 2.86E-10 |
| NM_006293 | TYRO3 | 1167 | 1105 | 509 | 509 | -1.2 | 7.60E-10 |
| NM_173354 | SIK1 | 224 | 1085 | 207 | 497 | -1.2 | 1.63E-09 |
| NM_001320643 | LOC102724428 | 223 | 1078 | 205 | 493 | -1.2 | 1.67E-09 |
| NM_004055 | CAPN5 | 585 | 564 | 1154 | 1206 | 1.1 | 3.44E-09 |

| | | | | | | | |
|---|---|---|---|---|---|---|---|
| NM_001031804 | MAF | 159 | 180 | 496 | 534 | 1.5 | 1.66E-08 |
| NM_001167928 | IL1RAP | 678 | 788 | 308 | 297 | -1.4 | 2.05E-08 |
| NM_001943 | DSG2 | 665 | 637 | 1389 | 1317 | 1.0 | 3.69E-08 |
| NM_153334 | SCARF2 | 252 | 231 | 51 | 49 | -2.3 | 5.39E-08 |
| NM_001330183 | SLFN5 | 177 | 158 | 500 | 443 | 1.4 | 7.49E-08 |
| NM_033046 | RTKN | 523 | 529 | 1125 | 1088 | 1.0 | 7.82E-08 |
| NM_000599 | IGFBP5 | 173 | 206 | 505 | 608 | 1.5 | 1.22E-07 |
| NM_005233 | EPHA3 | 167 | 179 | 650 | 647 | 1.8 | 1.64E-07 |
| NM_017947 | MOCOS | 852 | 896 | 1885 | 1928 | 1.1 | 1.64E-07 |
| NM_005978 | S100A2 | 310 | 351 | 873 | 840 | 1.2 | 1.69E-07 |
| NM_002839 | PTPRD | 418 | 471 | 170 | 161 | -1.6 | 2.41E-07 |
| NM_001146276 | NCEH1 | 943 | 1038 | 552 | 511 | -1.1 | 2.65E-07 |
| NM_001320977 | MAN2A2 | 1880 | 1780 | 942 | 884 | -1.1 | 2.84E-07 |
| NM_152342 | CDYL2 | 202 | 203 | 504 | 509 | 1.3 | 8.00E-07 |
| NM_001306080 | LMO7 | 611 | 637 | 288 | 294 | -1.2 | 9.67E-07 |
| NM_001201401 | GALC | 21 | 26 | 124 | 152 | 2.5 | 1.09E-06 |
| NM_001127323 | GRM8 | 24 | 32 | 262 | 277 | 3.1 | 1.15E-06 |
| NM_052947 | ALPK2 | 494 | 443 | 262 | 188 | -1.3 | 2.45E-06 |
| NM_020947 | TLDC1 | 383 | 346 | 739 | 722 | 1.0 | 3.66E-06 |
| NM_002421 | MMP1 | 367 | 361 | 181 | 139 | -1.4 | 8.50E-06 |
| NM_001270508 | TNFAIP3 | 224 | 715 | 253 | 351 | -1.1 | 8.55E-06 |
| NR_135681 | LOC105370792 | 615 | 685 | 244 | 290 | -1.3 | 1.61E-05 |
| NM_153690 | FAM43A | 232 | 536 | 165 | 249 | -1.1 | 1.61E-05 |
| NM_031288 | INO80B | 363 | 321 | 656 | 661 | 1.0 | 1.68E-05 |
| NM_020747 | ZNF608 | 223 | 194 | 531 | 458 | 1.2 | 1.95E-05 |
| NM_004370 | COL12A1 | 689 | 671 | 391 | 332 | -1.1 | 2.43E-05 |
| NM_001040197 | AGTRAP | 344 | 262 | 573 | 565 | 1.1 | 2.61E-05 |
| NM_015036 | ENDOD1 | 1144 | 1104 | 572 | 558 | -1.0 | 2.70E-05 |
| NM_022062 | PKNOX2 | 65 | 58 | 296 | 317 | 2.4 | 2.95E-05 |
| NM_006218 | PIK3CA | 775 | 846 | 490 | 417 | -1.1 | 3.60E-05 |
| NR_037654 | MTHFS | 721 | 681 | 369 | 343 | -1.0 | 3.98E-05 |
| NM_001199054 | CDIP1 | 302 | 273 | 479 | 566 | 1.0 | 4.43E-05 |
| NM_024697 | ZNF385D | 81 | 71 | 6 | 6 | -3.6 | 4.68E-05 |
| NR_135199 | FAM78B | 139 | 163 | 419 | 432 | 1.4 | 4.68E-05 |
| NM_001199760 | ST20-MTHFS | 681 | 664 | 353 | 329 | -1.1 | 5.14E-05 |

| | | | | | | | |
|---|---|---|---|---|---|---|---|
| NM_015187 | SEL1L3 | 510 | 575 | 238 | 260 | -1.2 | 5.55E-05 |
| NM_001321027 | AKR1C2 | 19 | 23 | 141 | 146 | 2.6 | 5.82E-05 |
| NM_001286565 | KIAA0513 | 60 | 61 | 179 | 203 | 1.7 | 7.93E-05 |
| NM_025135 | FHOD3 | 100 | 93 | 294 | 307 | 1.7 | 8.85E-05 |
| NM_001127401 | YPEL5 | 187 | 196 | 332 | 430 | 1.1 | 9.56E-05 |
| NM_001964 | EGR1 | 153 | 191 | 424 | 450 | 1.2 | 0.000103 |
| NM_014615 | GSE1 | 194 | 218 | 422 | 460 | 1.0 | 0.000114 |
| NM_001347006 | TMEM62 | 662 | 668 | 330 | 318 | -1.1 | 0.000114 |
| NM_033544 | RCCD1 | 426 | 419 | 233 | 197 | -1.1 | 0.000121 |
| NM_002998 | SDC2 | 91 | 110 | 332 | 348 | 1.6 | 0.000137 |
| NM_018192 | P3H2 | 298 | 301 | 124 | 128 | -1.3 | 0.000139 |
| NM_004775 | B4GALT6 | 385 | 365 | 820 | 778 | 1.1 | 0.000188 |
| NM_004058 | CAPS | 316 | 309 | 2066 | 2436 | 2.9 | 0.000211 |
| NM_001318503 | DUSP9 | 126 | 136 | 25 | 33 | -2.1 | 0.000214 |
| NM_080680 | COL11A2 | 125 | 119 | 253 | 306 | 1.3 | 0.000268 |
| NM_015931 | SSUH2 | 40 | 29 | 133 | 120 | 2.0 | 0.000331 |
| NM_001080509 | TSPAN11 | 23 | 20 | 169 | 140 | 2.8 | 0.000332 |
| NM_005427 | TP73 | 101 | 97 | 275 | 261 | 1.4 | 0.000333 |
| NM_001242409 | GAREM1 | 61 | 52 | 176 | 169 | 1.7 | 0.000343 |
| NM_000922 | PDE3B | 53 | 67 | 14 | 9 | -2.9 | 0.000343 |
| NR_038278 | LINC00665 | 15 | 17 | 92 | 93 | 2.4 | 0.000361 |
| NM_001321731 | EXOC6B | 215 | 213 | 449 | 450 | 1.0 | 0.000384 |
| NM_021012 | KCNJ12 | 91 | 76 | 203 | 225 | 1.5 | 0.000393 |
| NM_006169 | NNMT | 1980 | 2407 | 1146 | 1188 | -1.1 | 0.000564 |
| NR_135044 | LOC100506691 | 10 | 5 | 91 | 74 | 3.7 | 0.0006 |
| NM_198964 | PTHLH | 119 | 163 | 288 | 363 | 1.1 | 0.000618 |
| NM_152243 | CDC42EP1 | 210 | 209 | 69 | 71 | -1.6 | 0.000658 |
| NM_052916 | RNF157 | 133 | 146 | 364 | 361 | 1.3 | 0.000715 |
| NM_001206572 | SORCS1 | 274 | 305 | 715 | 738 | 1.2 | 0.000715 |
| NM_198536 | TMEM205 | 348 | 325 | 679 | 683 | 1.0 | 0.000715 |
| NM_001030313 | CERKL | 227 | 286 | 150 | 133 | -1.1 | 0.000914 |
| NM_006914 | RORB | 39 | 33 | 133 | 126 | 1.9 | 0.000933 |
| NM_014271 | IL1RAPL1 | 68 | 78 | 232 | 230 | 1.5 | 0.000957 |
| NM_018159 | NUDT11 | 49 | 42 | 132 | 148 | 1.8 | 0.000992 |
| NM_006237 | POU4F1 | 4 | 5 | 60 | 57 | 3.6 | 0.001012 |
| NM_004864 | GDF15 | 164 | 200 | 26 | 46 | -2.1 | 0.0011 |

| | | | | | | | |
|---|---|---|---|---|---|---|---|
| NM_001199039 | SERINC2 | 200 | 146 | 381 | 385 | 1.4 | 0.001116 |
| NR_134605 | LOC101927809 | 51 | 86 | 200 | 221 | 1.3 | 0.001332 |
| NM_001098817 | INO80C | 100 | 136 | 289 | 301 | 1.1 | 0.001615 |
| NM_153456 | HS6ST3 | 41 | 26 | 94 | 112 | 2.1 | 0.002077 |
| NM_000713 | BLVRB | 84 | 87 | 224 | 240 | 1.4 | 0.002098 |
| NM_003466 | PAX8 | 34 | 36 | 113 | 128 | 1.8 | 0.002368 |
| NM_003598 | TEAD2 | 61 | 61 | 187 | 177 | 1.5 | 0.002656 |
| NM_001099294 | KIAA1644 | 21 | 27 | 156 | 166 | 2.6 | 0.003902 |
| NM_001635 | AMPH | 3 | 2 | 32 | 38 | 4.2 | 0.004022 |
| NM_053279 | FAM167A | 44 | 49 | 165 | 171 | 1.8 | 0.004022 |
| NM_054025 | B3GAT1 | 37 | 25 | 97 | 95 | 1.9 | 0.004379 |
| NM_033225 | CSMD1 | 12 | 10 | 69 | 65 | 2.7 | 0.00531 |
| NM_153813 | ZFPM1 | 97 | 75 | 163 | 191 | 1.3 | 0.005406 |
| NM_001322891 | ABLIM1 | 1049 | 1010 | 491 | 502 | -1.0 | 0.006399 |
| NM_001062 | TCN1 | 64 | 70 | 198 | 196 | 1.5 | 0.00642 |
| NM_001195470 | SATB1 | 99 | 78 | 229 | 213 | 1.4 | 0.006681 |
| NR_040027 | ZNF790-AS1 | 9 | 8 | 49 | 52 | 2.6 | 0.006708 |
| NM_021910 | FXYD3 | 342 | 339 | 727 | 801 | 1.2 | 0.007121 |
| NM_004385 | VCAN | 95 | 138 | 417 | 392 | 1.5 | 0.007236 |
| NM_001011554 | SLC13A3 | 39 | 27 | 78 | 103 | 1.9 | 0.007575 |
| NM_032307 | C9orf64 | 149 | 124 | 58 | 42 | -1.6 | 0.007687 |
| NM_001040272 | ADAMTSL1 | 105 | 104 | 54 | 33 | -1.7 | 0.009607 |
| NR_134982 | CYP2J2 | 17 | 10 | 73 | 77 | 2.9 | 0.009623 |
| NM_153374 | LYSMD2 | 287 | 253 | 123 | 116 | -1.2 | 0.010015 |
| NM_000185 | SERPIND1 | 35 | 30 | 109 | 107 | 1.8 | 0.010446 |
| NM_020163 | SEMA3G | 33 | 33 | 104 | 124 | 1.9 | 0.010446 |
| NM_138736 | GNAO1 | 22 | 17 | 87 | 90 | 2.4 | 0.012123 |
| NM_001159920 | FLT1 | 100 | 116 | 330 | 318 | 1.4 | 0.012828 |
| NM_001314077 | PROS1 | 199 | 198 | 100 | 94 | -1.1 | 0.012975 |
| NR_109862 | BRCAT54 | 8 | 6 | 68 | 66 | 3.3 | 0.012975 |
| NM_006158 | NEFL | 240 | 244 | 478 | 539 | 1.1 | 0.013259 |
| NM_001322988 | KALRN | 71 | 77 | 205 | 180 | 1.2 | 0.013279 |
| NM_031421 | TTC25 | 69 | 45 | 106 | 125 | 1.4 | 0.013386 |
| NM_031910 | C1QTNF6 | 40 | 48 | 11 | 8 | -2.7 | 0.013638 |
| NM_003597 | KLF11 | 93 | 135 | 217 | 287 | 1.1 | 0.013888 |

| | | | | | | | |
|---|---|---|---|---|---|---|---|
| NM_015617 | PYGO1 | 245 | 210 | 91 | 106 | -1.0 | 0.014896 |
| NM_198253 | TERT | 147 | 140 | 51 | 56 | -1.4 | 0.015592 |
| NM_005101 | ISG15 | 30 | 18 | 58 | 71 | 1.9 | 0.016414 |
| NM_004925 | AQP3 | 72 | 100 | 224 | 241 | 1.2 | 0.016869 |
| NR_033869 | LINC01060 | 26 | 26 | 68 | 85 | 1.7 | 0.017142 |
| NM_006945 | SPRR2D | 105 | 148 | 233 | 328 | 1.1 | 0.017335 |
| NM_001290046 | CECR2 | 112 | 119 | 37 | 41 | -1.6 | 0.017603 |
| NM_004369 | COL6A3 | 305 | 327 | 1404 | 1322 | 2.0 | 0.018033 |
| NM_005114 | HS3ST1 | 90 | 97 | 29 | 35 | -1.5 | 0.018348 |
| NM_018490 | LGR4 | 409 | 407 | 154 | 167 | -1.3 | 0.018403 |
| NM_018936 | PCDHB2 | 65 | 73 | 215 | 194 | 1.4 | 0.018409 |
| NM_152339 | SPATA2L | 51 | 85 | 153 | 199 | 1.2 | 0.018529 |
| NM_003719 | PDE8B | 67 | 66 | 17 | 17 | -2.0 | 0.019325 |
| NM_005429 | VEGFC | 254 | 242 | 96 | 123 | -1.0 | 0.020468 |
| NM_000071 | CBS | 194 | 197 | 98 | 89 | -1.2 | 0.020622 |
| NM_001321073 | CBSL | 176 | 178 | 88 | 80 | -1.2 | 0.020622 |
| NM_138420 | AHNAK2 | 175 | 172 | 393 | 361 | 1.0 | 0.020955 |
| NM_001104554 | PAQR5 | 40 | 39 | 8 | 6 | -2.8 | 0.020955 |
| NM_004913 | VPS9D1 | 121 | 96 | 225 | 205 | 1.1 | 0.021968 |
| NR_077247 | RPS18P9 | 91 | 92 | 180 | 196 | 1.1 | 0.02445 |
| NM_012213 | MLYCD | 61 | 72 | 159 | 172 | 1.2 | 0.024657 |
| NM_020179 | SMCO4 | 235 | 218 | 98 | 104 | -1.1 | 0.025156 |
| NM_001319137 | CTSH | 58 | 40 | 91 | 110 | 1.4 | 0.027539 |
| NM_012476 | VAX2 | 8 | 3 | 20 | 31 | 3.6 | 0.027968 |
| NM_057159 | LPAR1 | 35 | 43 | 5 | 8 | -2.5 | 0.030527 |
| NR_033312 | BDNF-AS | 72 | 130 | 47 | 40 | -1.7 | 0.031428 |
| NR_029193 | SH3RF3-AS1 | 120 | 181 | 93 | 88 | -1.1 | 0.03163 |
| NM_207363 | NCKAP5 | 98 | 105 | 46 | 34 | -1.7 | 0.03351 |
| NR_015379 | UCA1 | 30 | 45 | 166 | 159 | 1.8 | 0.033584 |
| NM_001127464 | ZNF469 | 49 | 64 | 128 | 166 | 1.3 | 0.035072 |
| NM_000499 | CYP1A1 | 115 | 103 | 248 | 224 | 1.1 | 0.038846 |
| NM_006633 | IQGAP2 | 27 | 20 | 103 | 103 | 2.3 | 0.038894 |
| NM_213647 | FGFR4 | 110 | 99 | 203 | 221 | 1.1 | 0.040176 |
| NM_000927 | ABCB1 | 86 | 70 | 150 | 154 | 1.1 | 0.04267 |
| NM_080826 | ISM1 | 12 | 8 | 44 | 52 | 2.7 | 0.044306 |
| NM_031476 | CRISPLD2 | 23 | 17 | 74 | 73 | 2.0 | 0.04572 |
| NM_002345 | LUM | 108 | 100 | 32 | 32 | -1.6 | 0.046431 |

| Transcript ID | Gene name | CTRL_ DMSO counts | CTRL_ FSK counts | CRTC3KO_ DMSO counts | CRTC3KO_ FSK counts | CRTC3KO_ DMSO vs. CTRL_ DMSO Log2 Fold Change | CRTC3KO_ DMSO vs. CTRL_ DMSO adj. p-value |
|---|---|---|---|---|---|---|---|
| NM_182899 | CREB5 | 628 | 598 | 45 | 38 | -3.8 | 2.44E-49 |
| NM_001823 | CKB | 95 | 137 | 896 | 966 | 3.2 | 4.18E-46 |
| NM_003246 | THBS1 | 44240 | 44474 | 21749 | 17210 | -1.1 | 1.81E-24 |
| NM_022842 | CDCP1 | 240 | 246 | 10 | 15 | -4.7 | 1.31E-21 |
| NM_021214 | ABHD17C | 1799 | 2438 | 715 | 863 | -1.4 | 1.69E-20 |
| NM_017831 | RNF125 | 211 | 200 | 777 | 794 | 1.9 | 1.69E-20 |
| NM_001103184 | FMN1 | 2209 | 2193 | 977 | 899 | -1.2 | 3.42E-18 |
| NM_014268 | MAPRE2 | 690 | 720 | 1753 | 1763 | 1.3 | 7.68E-18 |
| NM_004811 | LPXN | 334 | 372 | 991 | 999 | 1.5 | 1.05E-16 |
| NM_005233 | EPHA3 | 167 | 179 | 650 | 647 | 1.9 | 3.33E-16 |
| NM_201432 | GAS7 | 1182 | 1199 | 3964 | 4085 | 1.7 | 4.94E-16 |
| NM_001005340 | GPNMB | 849 | 827 | 300 | 292 | -1.5 | 1.71E-13 |
| NM_005978 | S100A2 | 310 | 351 | 873 | 840 | 1.5 | 2.06E-13 |
| NM_001293298 | CEMIP | 5203 | 5668 | 1997 | 2052 | -1.4 | 3.63E-12 |
| NM_007085 | FSTL1 | 452 | 436 | 113 | 103 | -2.0 | 5.63E-12 |
| NM_012304 | FBXL7 | 5 | 4 | 196 | 179 | 5.2 | 8.20E-12 |
| NM_006293 | TYRO3 | 1167 | 1105 | 509 | 509 | -1.2 | 1.01E-11 |
| NR_104309 | ULK3 | 1687 | 1688 | 827 | 805 | -1.1 | 2.17E-11 |
| NM_024652 | LRRK1 | 325 | 363 | 67 | 72 | -2.3 | 2.23E-11 |
| NM_006472 | TXNIP | 140 | 134 | 674 | 640 | 2.2 | 3.18E-11 |
| NM_002587 | PCDH1 | 392 | 375 | 1339 | 1525 | 1.7 | 3.56E-11 |
| NM_015476 | TPGS2 | 24 | 32 | 262 | 277 | 1.0 | 5.01E-11 |
| NM_001127323 | GRM8 | 986 | 952 | 2019 | 1975 | 3.4 | 5.01E-11 |
| NM_000599 | IGFBP5 | 173 | 206 | 505 | 608 | 1.5 | 6.40E-11 |

| | | | | | | | |
|---|---|---|---|---|---|---|---|
| NM_006965 | ZNF24 | 839 | 848 | 1779 | 1594 | 1.1 | 1.04E-10 |
| NM_001037340 | PDE4B | 324 | 531 | 70 | 77 | -2.2 | 7.24E-10 |
| NM_153334 | SCARF2 | 252 | 231 | 51 | 49 | -2.3 | 8.12E-10 |
| NM_033046 | RTKN | 523 | 529 | 1125 | 1088 | 1.1 | 2.73E-09 |
| NM_015036 | ENDOD1 | 1144 | 1104 | 572 | 558 | -1.0 | 8.42E-09 |
| NM_001330183 | SLFN5 | 177 | 158 | 500 | 443 | 1.5 | 8.80E-09 |
| NM_017947 | MOCOS | 852 | 896 | 1885 | 1928 | 1.1 | 9.27E-09 |
| NM_001320977 | MAN2A2 | 1880 | 1780 | 942 | 884 | -1.0 | 9.43E-09 |
| NM_005929 | MELTF | 3437 | 3369 | 1693 | 1813 | -1.0 | 1.98E-08 |
| NM_001943 | DSG2 | 665 | 637 | 1389 | 1317 | 1.0 | 4.18E-08 |
| NM_014939 | TRAPPC8 | 358 | 373 | 769 | 745 | 1.1 | 7.53E-08 |
| NM_004775 | B4GALT6 | 385 | 365 | 820 | 778 | 1.1 | 9.64E-08 |
| NM_052916 | RNF157 | 133 | 146 | 364 | 361 | 1.4 | 2.41E-07 |
| NM_020747 | ZNF608 | 223 | 194 | 531 | 458 | 1.2 | 2.71E-07 |
| NM_001286741 | UBL7 | 1971 | 1924 | 986 | 1047 | -1.0 | 3.41E-07 |
| NM_002839 | PTPRD | 418 | 471 | 170 | 161 | -1.3 | 6.53E-07 |
| NM_001031804 | MAF | 159 | 180 | 496 | 534 | 1.6 | 6.53E-07 |
| NR_134605 | LOC101927809 | 51 | 86 | 200 | 221 | 1.9 | 9.77E-07 |
| NM_001306080 | LMO7 | 611 | 637 | 288 | 294 | -1.1 | 1.20E-06 |
| NM_024697 | ZNF385D | 81 | 71 | 6 | 6 | -3.8 | 1.25E-06 |
| NM_001098817 | INO80C | 100 | 136 | 289 | 301 | 1.5 | 2.01E-06 |
| NM_001080509 | TSPAN11 | 23 | 20 | 169 | 140 | 2.9 | 2.19E-06 |
| NM_001964 | EGR1 | 153 | 191 | 424 | 450 | 1.4 | 2.28E-06 |
| NM_152342 | CDYL2 | 202 | 203 | 504 | 509 | 1.3 | 2.33E-06 |
| NM_153840 | ADGRF1 | 49 | 50 | 188 | 116 | 1.9 | 4.02E-06 |
| NM_145658 | SPESP1 | 146 | 136 | 0 | 0 | -8.3 | 4.03E-06 |
| NM_004058 | CAPS | 316 | 309 | 2066 | 2436 | 2.7 | 4.17E-06 |

| | | | | | | | |
|---|---|---|---|---|---|---|---|
| NM_001201401 | GALC | 21 | 26 | 124 | 152 | 2.6 | 6.42E-06 |
| NR_135199 | FAM78B | 139 | 163 | 419 | 432 | 1.6 | 6.92E-06 |
| NM_014271 | IL1RAPL1 | 68 | 78 | 232 | 230 | 1.7 | 8.29E-06 |
| NM_006183 | NTS | 64 | 76 | 2 | 7 | -5.1 | 1.04E-05 |
| NM_015617 | PYGO1 | 245 | 210 | 91 | 106 | -1.5 | 1.11E-05 |
| NM_015187 | SEL1L3 | 510 | 575 | 238 | 260 | -1.1 | 1.54E-05 |
| NM_001318503 | DUSP9 | 126 | 136 | 25 | 33 | -2.3 | 1.54E-05 |
| NM_001122679 | TENM2 | 3 | 15 | 64 | 57 | 4.2 | 2.03E-05 |
| NR_135681 | LOC105370792 | 615 | 685 | 244 | 290 | -1.4 | 2.13E-05 |
| NM_005427 | TP73 | 101 | 97 | 275 | 261 | 1.4 | 2.14E-05 |
| NM_001167928 | IL1RAP | 678 | 788 | 308 | 297 | -1.2 | 2.27E-05 |
| NM_014615 | GSE1 | 194 | 218 | 422 | 460 | 1.1 | 2.27E-05 |
| NM_001321027 | AKR1C2 | 19 | 23 | 141 | 146 | 2.9 | 2.80E-05 |
| NM_002998 | SDC2 | 91 | 110 | 332 | 348 | 1.8 | 3.20E-05 |
| NM_001206572 | SORCS1 | 274 | 305 | 715 | 738 | 1.4 | 4.26E-05 |
| NM_018490 | LGR4 | 95 | 138 | 417 | 392 | -1.4 | 4.43E-05 |
| NM_004385 | VCAN | 409 | 407 | 154 | 167 | 2.1 | 4.43E-05 |
| NM_004864 | GDF15 | 164 | 200 | 26 | 46 | -2.7 | 4.58E-05 |
| NM_018192 | P3H2 | 298 | 301 | 124 | 128 | -1.3 | 4.81E-05 |
| NM_001347006 | TMEM62 | 662 | 668 | 330 | 318 | -1.0 | 5.36E-05 |
| NM_153374 | LYSMD2 | 287 | 253 | 123 | 116 | -1.2 | 6.16E-05 |
| NM_198964 | PTHLH | 119 | 163 | 288 | 363 | 1.2 | 6.16E-05 |
| NM_005429 | VEGFC | 254 | 242 | 96 | 123 | -1.4 | 6.41E-05 |
| NM_018936 | PCDHB2 | 65 | 73 | 215 | 194 | 1.7 | 6.53E-05 |
| NM_004003 | CRAT | 201 | 201 | 430 | 400 | 1.1 | 6.70E-05 |
| NR_038278 | LINC00665 | 15 | 17 | 92 | 93 | 2.6 | 7.41E-05 |
| NM_025135 | FHOD3 | 100 | 93 | 294 | 307 | 1.5 | 8.02E-05 |

| | | | | | | | |
|---|---|---|---|---|---|---|---|
| NM_001322891 | ABLIM1 | 1049 | 1010 | 491 | 502 | -1.1 | 9.54E-05 |
| NM_022062 | PKNOX2 | 65 | 58 | 296 | 317 | 2.1 | 9.87E-05 |
| NM_001322988 | KALRN | 71 | 77 | 205 | 180 | 1.5 | 0.00023 |
| NM_175634 | RUNX1T1 | 6 | 8 | 70 | 50 | 3.6 | 0.000251 |
| NM_001242409 | GAREM1 | 61 | 52 | 176 | 169 | 1.5 | 0.0004 |
| NM_005366 | MAGEA11 | 801 | 838 | 328 | 333 | -1.3 | 0.000475 |
| NM_001321731 | EXOC6B | 215 | 213 | 449 | 450 | 1.0 | 0.000562 |
| NM_006914 | RORB | 39 | 33 | 133 | 126 | 1.7 | 0.000579 |
| NM_015931 | SSUH2 | 40 | 29 | 133 | 120 | 1.7 | 0.000592 |
| NM_152339 | SPATA2L | 51 | 85 | 153 | 199 | 1.5 | 0.000705 |
| NM_006237 | POU4F1 | 4 | 5 | 60 | 57 | 3.8 | 0.000771 |
| NM_000713 | BLVRB | 84 | 87 | 224 | 240 | 1.4 | 0.000908 |
| NM_004925 | AQP3 | 72 | 100 | 224 | 241 | 1.6 | 0.001035 |
| NM_001159920 | FLT1 | 100 | 116 | 330 | 318 | 1.7 | 0.001062 |
| NM_033225 | CSMD1 | 12 | 10 | 69 | 65 | 2.5 | 0.001216 |
| NM_002421 | MMP1 | 367 | 361 | 181 | 139 | -1.0 | 0.001217 |
| NM_003597 | KLF11 | 93 | 135 | 217 | 287 | 1.2 | 0.001509 |
| NM_000526 | KRT14 | 119 | 148 | 268 | 303 | 1.1 | 0.001746 |
| NM_152243 | CDC42EP1 | 210 | 209 | 69 | 71 | -1.6 | 0.001808 |
| NM_001286565 | KIAA0513 | 60 | 61 | 179 | 203 | 1.5 | 0.002228 |
| NM_001297651 | MAST4 | 78 | 91 | 183 | 171 | 1.2 | 0.002812 |
| NM_024522 | NKAIN1 | 16 | 27 | 73 | 74 | 2.2 | 0.002917 |
| NM_138420 | AHNAK2 | 175 | 172 | 393 | 361 | 1.1 | 0.003629 |
| NM_001290046 | CECR2 | 112 | 119 | 37 | 41 | -1.6 | 0.003955 |
| NM_003466 | PAX8 | 34 | 36 | 113 | 128 | 1.7 | 0.004111 |
| NR_024063 | ZSCAN12P1 | 53 | 53 | 9 | 11 | -2.6 | 0.004695 |
| NM_001062 | TCN1 | 64 | 70 | 198 | 196 | 1.6 | 0.004953 |

| | | | | | | | |
|---|---|---|---|---|---|---|---|
| NM_001409 | MEGF6 | 38 | 28 | 4 | 7 | -3.1 | 0.00515 |
| NM_021012 | KCNJ12 | 91 | 76 | 203 | 225 | 1.1 | 0.005178 |
| NM_001301036 | TMCC3 | 147 | 183 | 305 | 296 | 1.0 | 0.00554 |
| NM_012213 | MLYCD | 61 | 72 | 159 | 172 | 1.3 | 0.00555 |
| NM_003598 | TEAD2 | 61 | 61 | 187 | 177 | 1.6 | 0.005602 |
| NM_020179 | SMCO4 | 235 | 218 | 98 | 104 | -1.3 | 0.006148 |
| NM_032679 | ZNF577 | 18 | 19 | 74 | 63 | 2.0 | 0.006238 |
| NM_005114 | HS3ST1 | 57 | 64 | 152 | 127 | -1.7 | 0.006868 |
| NM_016260 | IKZF2 | 90 | 97 | 29 | 35 | 1.4 | 0.006868 |
| NM_002546 | TNFRSF11B | 116 | 97 | 45 | 47 | -1.4 | 0.007316 |
| NM_000185 | SERPIND1 | 35 | 30 | 109 | 107 | 1.6 | 0.008312 |
| NR_038285 | MELTF-AS1 | 192 | 189 | 88 | 106 | -1.2 | 0.008683 |
| NM_003719 | PDE8B | 67 | 66 | 17 | 17 | -2.0 | 0.009043 |
| NM_018159 | NUDT11 | 49 | 42 | 132 | 148 | 1.4 | 0.009605 |
| NM_003059 | SLC22A4 | 53 | 67 | 134 | 126 | 1.3 | 0.010028 |
| NM_001314077 | PROS1 | 199 | 198 | 100 | 94 | -1.0 | 0.010068 |
| NR_038977 | LINC01239 | 81 | 85 | 183 | 173 | 1.1 | 0.01043 |
| NM_001321866 | ZNF600 | 14 | 16 | 64 | 60 | 2.1 | 0.010798 |
| NM_032812 | PLXDC2 | 27 | 23 | 1 | 3 | -4.8 | 0.011048 |
| NR_040027 | ZNF790-AS1 | 9 | 8 | 49 | 52 | 2.5 | 0.011221 |
| NM_001099280 | MROH1 | 38 | 49 | 107 | 77 | 1.5 | 0.012191 |
| NM_198253 | TERT | 147 | 140 | 51 | 56 | -1.6 | 0.012219 |
| NM_004572 | PKP2 | 260 | 196 | 118 | 134 | -1.2 | 0.012258 |
| NM_004845 | PCYT1B | 38 | 38 | 106 | 94 | 1.5 | 0.012357 |
| NM_021910 | FXYD3 | 342 | 339 | 727 | 801 | 1.1 | 0.014001 |
| NM_018015 | CXorf57 | 11 | 13 | 51 | 51 | 2.2 | 0.015392 |
| NM_000641 | IL11 | 182 | 605 | 398 | 629 | 1.1 | 0.015559 |

| | | | | | | | |
|---|---|---|---|---|---|---|---|
| NM_000499 | CYP1A1 | 115 | 103 | 248 | 224 | 1.1 | 0.01607 |
| NM_021205 | RHOU | 16 | 26 | 65 | 66 | 2.0 | 0.016403 |
| NM_003182 | TAC1 | 2 | 8 | 34 | 23 | 3.8 | 0.016764 |
| NM_053279 | FAM167A | 44 | 49 | 165 | 171 | 1.9 | 0.016972 |
| NM_002345 | LUM | 108 | 100 | 32 | 32 | -1.8 | 0.017545 |
| NM_015852 | ZNF117 | 36 | 36 | 98 | 85 | 1.4 | 0.017663 |
| NM_018849 | ABCB4 | 4 | 8 | 34 | 32 | 3.0 | 0.018758 |
| NM_025208 | PDGFD | 127 | 150 | 48 | 85 | -1.4 | 0.018847 |
| NM_001290040 | ROBO2 | 3 | 4 | 29 | 14 | 3.3 | 0.02126 |
| NM_172362 | KCNH1 | 162 | 190 | 76 | 103 | -1.1 | 0.021736 |
| NM_004644 | AP3B2 | 23 | 32 | 74 | 90 | 1.6 | 0.022678 |
| NM_031476 | CRISPLD2 | 23 | 17 | 74 | 73 | 1.7 | 0.022715 |
| NM_054025 | B3GAT1 | 37 | 25 | 97 | 95 | 1.4 | 0.023041 |
| NM_001104554 | PAQR5 | 68 | 82 | 170 | 171 | -2.4 | 0.023411 |
| NM_013259 | TAGLN3 | 40 | 39 | 8 | 6 | 1.3 | 0.023411 |
| NM_002425 | MMP10 | 42 | 36 | 9 | 8 | -2.3 | 0.02408 |
| NM_002982 | CCL2 | 24 | 11 | 0 | 3 | -5.6 | 0.024599 |
| NM_000071 | CBS | 194 | 197 | 98 | 89 | -1.0 | 0.025606 |
| NM_001127464 | ZNF469 | 49 | 64 | 128 | 166 | 1.4 | 0.025893 |
| NM_006729 | DIAPH2 | 151 | 163 | 325 | 321 | 1.1 | 0.026636 |
| NM_004984 | KIF5A | 4 | 6 | 35 | 26 | 3.0 | 0.026804 |
| NM_001195470 | SATB1 | 99 | 78 | 229 | 213 | 1.2 | 0.027033 |
| NM_004369 | COL6A3 | 305 | 327 | 1404 | 1322 | 2.2 | 0.027298 |
| NM_001369 | DNAH5 | 59 | 69 | 141 | 148 | 1.2 | 0.027907 |
| NM_032307 | C9orf64 | 149 | 124 | 58 | 42 | -1.4 | 0.028715 |
| NM_001321544 | SLC25A42 | 30 | 37 | 85 | 85 | 1.5 | 0.029152 |
| NM_002910 | RENBP | 27 | 25 | 3 | 2 | -3.3 | 0.030134 |

| | | | | | | | |
|---|---|---|---|---|---|---|---|
| NM_020335 | VANGL2 | 23 | 31 | 83 | 88 | 1.8 | 0.032628 |
| NM_020163 | SEMA3G | 33 | 33 | 104 | 124 | 1.6 | 0.033867 |
| NM_001321073 | CBSL | 176 | 178 | 88 | 80 | -1.0 | 0.035995 |
| NR_109862 | BRCAT54 | 8 | 6 | 68 | 66 | 3.1 | 0.03703 |
| NM_001318880 | STK32C | 65 | 58 | 141 | 126 | 1.1 | 0.038909 |
| NR_134982 | CYP2J2 | 17 | 10 | 73 | 77 | 2.1 | 0.039315 |
| NM_006945 | SPRR2D | 105 | 148 | 233 | 328 | 1.1 | 0.039547 |
| NM_138736 | GNAO1 | 22 | 17 | 87 | 90 | 2.0 | 0.04024 |
| NM_002397 | MEF2C | 61 | 80 | 131 | 134 | 1.1 | 0.045481 |
| NM_057159 | LPAR1 | 35 | 43 | 5 | 8 | -2.7 | 0.047693 |
| NM_001135178 | ZNF397 | 56 | 69 | 129 | 154 | 1.2 | 0.048648 |
| NM_001400 | S1PR1 | 76 | 93 | 17 | 19 | -2.2 | 0.049077 |

**Suppl. Table 5. CRTC1/2/3 positive correlations in melanoma patients (from TCGA Firehose Legacy at cbioportal.org).**

| CRTC3 | | | CRTC2 | | | CRTC1 | | |
|---|---|---|---|---|---|---|---|---|
| Gene | Spearman's Correlation | q-Value | Gene | Spearman's Correlation | q-Value | Gene | Spearman's Correlation | q-Value |
| ZNF592 | 0.599568 | 6.86E-33 | PYGO2 | 0.712994 | 6.25E-54 | ZNF414 | 0.559993 | 2.24E-27 |
| TBC1D2B | 0.545345 | 5.40E-26 | SF3B4 | 0.674477 | 4.98E-46 | MED26 | 0.534758 | 1.57E-24 |
| POLG | 0.540593 | 1.55E-25 | VPS72 | 0.673141 | 6.07E-46 | SUGP1 | 0.510524 | 6.35E-22 |
| UNC45A | 0.531508 | 1.53E-24 | DEDD | 0.637414 | 1.56E-39 | MYO9B | 0.50575 | 1.59E-21 |
| RAPGEF1 | 0.524894 | 7.57E-24 | SCNM1 | 0.633141 | 6.61E-39 | CCDC97 | 0.504814 | 1.60E-21 |
| VPS33B | 0.5229 | 1.10E-23 | PSMD4 | 0.61274 | 1.12E-35 | ZBTB45 | 0.497669 | 7.76E-21 |
| ZNF710 | 0.489843 | 3.05E-20 | PRCC | 0.597656 | 1.89E-33 | PACS2 | 0.495256 | 1.05E-20 |
| ITPKB | 0.475011 | 7.29E-19 | KRTCAP2 | 0.595954 | 2.94E-33 | MAP2K7 | 0.495242 | 1.05E-20 |
| OCA2 | 0.472084 | 1.22E-18 | HAX1 | 0.591729 | 1.08E-32 | SWSAP1 | 0.49172 | 2.17E-20 |

| | | | | | | | | |
|---|---|---|---|---|---|---|---|---|
| ZBED3 | 0.462802 | 7.86E-18 | FLAD1 | 0.584542 | 1.03E-31 | PIN1 | 0.490532 | 2.59E-20 |
| HEXA-AS1 | 0.460979 | 1.08E-17 | SCAMP3 | 0.584307 | 1.03E-31 | ALKBH7 | 0.484091 | 1.05E-19 |
| ARHGAP8 | 0.445237 | 1.98E-16 | MEF2D | 0.574013 | 2.53E-30 | BTBD2 | 0.483831 | 1.05E-19 |
| LIMS2 | 0.441363 | 4.22E-16 | ZNF672 | 0.572971 | 3.24E-30 | ZNF653 | 0.479886 | 2.40E-19 |
| PMEL | 0.439593 | 5.42E-16 | MTX1 | 0.5668 | 2.03E-29 | XAB2 | 0.474644 | 7.35E-19 |
| SNX33 | 0.436691 | 8.74E-16 | ZNF687 | 0.566373 | 2.16E-29 | TRAPPC6A | 0.473928 | 8.06E-19 |
| ARPIN | 0.432084 | 2.06E-15 | YY1AP1 | 0.565751 | 2.45E-29 | BRF1 | 0.473178 | 8.95E-19 |
| ZNF687 | 0.431119 | 2.37E-15 | DENND4B | 0.565447 | 2.53E-29 | PGLS | 0.469426 | 1.95E-18 |
| SLC45A2 | 0.429536 | 3.01E-15 | UBE2Q1 | 0.564584 | 3.11E-29 | MAP1S | 0.467793 | 2.64E-18 |
| FAM53B | 0.426431 | 4.92E-15 | FAM71D | 0.559108 | 1.54E-28 | PIK3R2 | 0.466061 | 3.66E-18 |
| RUNX3 | 0.421993 | 1.04E-14 | TRIM11 | 0.557565 | 2.31E-28 | RANBP3 | 0.463066 | 6.68E-18 |
| KCNAB2 | 0.420345 | 1.40E-14 | ARL8A | 0.553654 | 6.98E-28 | WIZ | 0.462263 | 7.58E-18 |
| PIK3CD | 0.418271 | 1.96E-14 | APH1A | 0.534294 | 1.62E-25 | RFXANK | 0.460305 | 1.10E-17 |
| MYO9B | 0.416829 | 2.52E-14 | SLC39A1 | 0.533194 | 2.10E-25 | RABGGTA | 0.457744 | 1.83E-17 |
| RAB32 | 0.416401 | 2.68E-14 | USP21 | 0.528831 | 6.58E-25 | ILVBL | 0.456594 | 2.24E-17 |
| SNN | 0.414639 | 3.55E-14 | B4GALT3 | 0.526942 | 1.05E-24 | ARHGEF18 | 0.455702 | 2.60E-17 |
| CYB561A3 | 0.41261 | 4.78E-14 | PMF1 | 0.523739 | 2.29E-24 | YJU2 | 0.455259 | 2.74E-17 |
| TRPM1 | 0.411404 | 5.79E-14 | RUSC1 | 0.521206 | 4.30E-24 | USF2 | 0.454846 | 2.88E-17 |
| MCF2L | 0.408836 | 8.63E-14 | MRPL9 | 0.519741 | 6.10E-24 | FKBP8 | 0.453787 | 3.48E-17 |
| SIRPB1 | 0.40676 | 1.15E-13 | JTB | 0.517697 | 1.01E-23 | SIN3B | 0.453193 | 3.80E-17 |
| UBAP1L | 0.40631 | 1.23E-13 | MAPKAPK2 | 0.514899 | 2.01E-23 | HDAC6 | 0.447862 | 1.12E-16 |
| CSK | 0.406193 | 1.24E-13 | PBXIP1 | 0.512202 | 3.88E-23 | UPF1 | 0.446841 | 1.33E-16 |
| SP2 | 0.40484 | 1.53E-13 | PMVK | 0.508637 | 9.31E-23 | ZNF575 | 0.446312 | 1.44E-16 |
| SCAMP2 | 0.404406 | 1.63E-13 | RRNAD1 | 0.50057 | 6.36E-22 | CCDC124 | 0.446001 | 1.46E-16 |
| DET1 | 0.404363 | 1.63E-13 | ZBTB7B | 0.495018 | 2.33E-21 | TYK2 | 0.445952 | 1.46E-16 |
| NEO1 | 0.404217 | 1.65E-13 | SLC25A44 | 0.490954 | 6.02E-21 | PWWP3A | 0.445303 | 1.62E-16 |
| RREB1 | 0.404162 | 1.65E-13 | UBAP2L | 0.490597 | 6.39E-21 | TECR | 0.444254 | 1.95E-16 |

| | | | | | | | | |
|---|---|---|---|---|---|---|---|---|
| OR7A5 | 0.403964 | 1.69E-13 | FAM189B | 0.486769 | 1.47E-20 | ENDOV | 0.442901 | 2.50E-16 |
| PCIF1 | 0.401748 | 2.40E-13 | SLC50A1 | 0.485227 | 2.06E-20 | OCEL1 | 0.441991 | 2.87E-16 |
| PLEKHG3 | 0.401634 | 2.40E-13 | PPOX | 0.483225 | 3.14E-20 | ASPSCR1 | 0.441906 | 2.87E-16 |
| TPPP | 0.400962 | 2.65E-13 | USF1 | 0.482339 | 3.77E-20 | TPRN | 0.441849 | 2.87E-16 |
| NMRK2 | 0.398423 | 3.90E-13 | DPM3 | 0.481701 | 4.28E-20 | FN3K | 0.441445 | 3.04E-16 |
| DIPK1B | 0.397808 | 4.28E-13 | LIN37 | 0.48026 | 5.73E-20 | C19ORF44 | 0.440576 | 3.53E-16 |
| NUP214 | 0.397396 | 4.56E-13 | LAMTOR2 | 0.479804 | 6.24E-20 | THAP8 | 0.440176 | 3.74E-16 |
| PRODH | 0.39713 | 4.69E-13 | TPM3 | 0.478755 | 7.78E-20 | DAPK3 | 0.439286 | 4.37E-16 |
| GNAL | 0.396561 | 5.04E-13 | NAXE | 0.477205 | 1.09E-19 | ZNF837 | 0.439069 | 4.47E-16 |
| TFAP2A | 0.395645 | 5.86E-13 | DUSP23 | 0.472374 | 3.12E-19 | TRMT61A | 0.438894 | 4.53E-16 |
| ZNF362 | 0.394716 | 6.82E-13 | MSTO1 | 0.469815 | 5.32E-19 | FLYWCH1 | 0.438531 | 4.76E-16 |
| MECP2 | 0.393356 | 8.48E-13 | MRPL55 | 0.468708 | 6.69E-19 | TRAPPC5 | 0.436194 | 6.86E-16 |
| MGAT5B | 0.393129 | 8.75E-13 | ARF1 | 0.463879 | 1.84E-18 | CFAP410 | 0.435953 | 7.07E-16 |
| IP6K1 | 0.392811 | 9.08E-13 | TARS2 | 0.463194 | 2.10E-18 | LRFN3 | 0.43297 | 1.25E-15 |
| SCARB1 | 0.392736 | 9.13E-13 | GATAD2B | 0.462905 | 2.16E-18 | SGTA | 0.429881 | 2.21E-15 |
| GPR143 | 0.39207 | 1.02E-12 | PI4KB | 0.46256 | 2.29E-18 | FZR1 | 0.429713 | 2.24E-15 |
| AGPAT3 | 0.391702 | 1.07E-12 | SUPT5H | 0.462478 | 2.30E-18 | MAP2K2 | 0.429534 | 2.28E-15 |
| AP5B1 | 0.391392 | 1.11E-12 | GUK1 | 0.458074 | 5.30E-18 | FAM98C | 0.428969 | 2.46E-15 |
| CDH3 | 0.389326 | 1.51E-12 | CXXC1 | 0.457865 | 5.42E-18 | KLHL26 | 0.427989 | 2.88E-15 |
| SIRPA | 0.389227 | 1.53E-12 | CERS2 | 0.457543 | 5.63E-18 | ZNF358 | 0.42661 | 3.70E-15 |
| EDC3 | 0.388593 | 1.67E-12 | CHTOP | 0.457234 | 5.93E-18 | ZER1 | 0.42399 | 5.93E-15 |
| PEPD | 0.388367 | 1.70E-12 | C1ORF35 | 0.457142 | 5.98E-18 | ZNF574 | 0.423981 | 5.93E-15 |
| FAM124A | 0.38685 | 2.14E-12 | PSMB4 | 0.454084 | 1.13E-17 | SYDE1 | 0.42295 | 7.11E-15 |
| GGA3 | 0.386316 | 2.32E-12 | HGS | 0.453512 | 1.23E-17 | EPS15L1 | 0.42128 | 9.62E-15 |
| SLC1A4 | 0.384955 | 2.82E-12 | CEP131 | 0.450905 | 2.10E-17 | RPTOR | 0.421088 | 9.83E-15 |
| VPS18 | 0.384115 | 3.17E-12 | RNF220 | 0.44964 | 2.61E-17 | TMEM205 | 0.420181 | 1.13E-14 |
| RASSF2 | 0.383099 | 3.69E-12 | PPP1R9B | 0.446732 | 4.63E-17 | RILPL1 | 0.420103 | 1.13E-14 |

| | | | | | | | | |
|---|---|---|---|---|---|---|---|---|
| ZFYVE26 | 0.382861 | 3.80E-12 | MAP1S | 0.445992 | 5.30E-17 | ZC3H4 | 0.418878 | 1.41E-14 |
| RIN3 | 0.382449 | 3.98E-12 | MRPL24 | 0.445623 | 5.65E-17 | RABL6 | 0.418264 | 1.56E-14 |
| IL16 | 0.382203 | 4.11E-12 | GBAP1 | 0.443972 | 7.67E-17 | RNF220 | 0.41811 | 1.58E-14 |
| WDR81 | 0.381916 | 4.22E-12 | ATXN2L | 0.442355 | 1.03E-16 | EXD3 | 0.417123 | 1.85E-14 |
| ZNF777 | 0.38131 | 4.58E-12 | RPP21 | 0.442081 | 1.06E-16 | ANAPC2 | 0.416541 | 2.04E-14 |
| REPS2 | 0.381178 | 4.65E-12 | MIIP | 0.441248 | 1.25E-16 | TMEM161A | 0.414958 | 2.67E-14 |
| CDK2 | 0.380775 | 4.94E-12 | PRR14 | 0.440489 | 1.40E-16 | NDUFB7 | 0.414878 | 2.67E-14 |
| IRF4 | 0.380375 | 5.20E-12 | SSR2 | 0.438192 | 2.10E-16 | BTBD6 | 0.414675 | 2.74E-14 |
| VPS53 | 0.380363 | 5.20E-12 | SIRT7 | 0.438021 | 2.16E-16 | CCDC130 | 0.41447 | 2.81E-14 |
| ARHGEF18 | 0.380183 | 5.33E-12 | NCSTN | 0.436862 | 2.60E-16 | RAD23A | 0.414051 | 3.00E-14 |
| DAB2 | 0.379619 | 5.74E-12 | SNAPIN | 0.436727 | 2.65E-16 | AP1M1 | 0.413858 | 3.06E-14 |
| HERC1 | 0.379048 | 6.20E-12 | GPATCH3 | 0.436195 | 2.93E-16 | ZNF526 | 0.413816 | 3.06E-14 |
| KLHL25 | 0.378654 | 6.51E-12 | UBQLN4 | 0.435648 | 3.21E-16 | CACTIN | 0.413734 | 3.07E-14 |
| AATBC | 0.378544 | 6.59E-12 | SUGP1 | 0.43452 | 3.86E-16 | TBC1D17 | 0.411547 | 4.54E-14 |
| RFLNB | 0.376393 | 9.06E-12 | NIT1 | 0.430237 | 8.09E-16 | ZNF784 | 0.411392 | 4.62E-14 |
| PPARD | 0.375991 | 9.53E-12 | CPTP | 0.430212 | 8.09E-16 | KXD1 | 0.410795 | 5.09E-14 |
| ATP11A | 0.3755 | 1.02E-11 | SURF2 | 0.429993 | 8.34E-16 | BICRA | 0.410549 | 5.27E-14 |
| CLN6 | 0.374996 | 1.10E-11 | CNPY3 | 0.429711 | 8.75E-16 | CHERP | 0.410422 | 5.33E-14 |
| NOP9 | 0.374141 | 1.25E-11 | PIPSL | 0.427123 | 1.40E-15 | PRR12 | 0.410125 | 5.56E-14 |
| RTN4R | 0.373775 | 1.30E-11 | DTX2 | 0.426885 | 1.44E-15 | MIB2 | 0.410027 | 5.60E-14 |
| DYNC1H1 | 0.372611 | 1.51E-11 | TUBGCP2 | 0.426456 | 1.56E-15 | SCAF1 | 0.408564 | 7.21E-14 |
| ZNF703 | 0.372591 | 1.51E-11 | RNPEP | 0.426404 | 1.56E-15 | LOC407835 | 0.407956 | 7.87E-14 |
| BRPF1 | 0.372575 | 1.51E-11 | TPRA1 | 0.426251 | 1.60E-15 | PEX10 | 0.407925 | 7.87E-14 |
| MED12 | 0.372562 | 1.51E-11 | C1ORF56 | 0.426038 | 1.65E-15 | NDUFA11 | 0.40749 | 8.43E-14 |
| GM2A | 0.372213 | 1.59E-11 | F8A1 | 0.425931 | 1.66E-15 | ZNF628 | 0.407416 | 8.46E-14 |
| USP36 | 0.372008 | 1.63E-11 | THAP3 | 0.425609 | 1.74E-15 | JUND | 0.407175 | 8.66E-14 |
| CEP68 | 0.371254 | 1.81E-11 | TSEN54 | 0.425525 | 1.76E-15 | BRD4 | 0.407067 | 8.74E-14 |

| | | | | | | | | |
|---|---|---|---|---|---|---|---|---|
| AKT1 | 0.370859 | 1.89E-11 | MRPS21 | 0.42459 | 2.07E-15 | RBM42 | 0.405857 | 1.08E-13 |
| DIPK1C | 0.370472 | 1.98E-11 | YPEL3 | 0.424169 | 2.21E-15 | REX1BD | 0.405119 | 1.22E-13 |
| CELSR2 | 0.369958 | 2.12E-11 | ZNF341 | 0.421653 | 3.34E-15 | TECPR2 | 0.404891 | 1.26E-13 |
| SEC16A | 0.369633 | 2.23E-11 | BOLA1 | 0.421327 | 3.47E-15 | UBXN6 | 0.404738 | 1.28E-13 |
| TMEM86A | 0.36831 | 2.69E-11 | FBXL15 | 0.421307 | 3.47E-15 | DOHH | 0.404248 | 1.38E-13 |
| PARP6 | 0.368119 | 2.76E-11 | RNF187 | 0.420913 | 3.69E-15 | PNPLA6 | 0.404076 | 1.41E-13 |
| TMEM268 | 0.367954 | 2.82E-11 | NDUFS2 | 0.419593 | 4.64E-15 | PDK2 | 0.40379 | 1.47E-13 |
| MFHAS1 | 0.367741 | 2.90E-11 | ZNF574 | 0.419495 | 4.70E-15 | LRSAM1 | 0.403183 | 1.61E-13 |
| SLC24A4 | 0.366508 | 3.41E-11 | SNAP47 | 0.418629 | 5.48E-15 | DHRS12 | 0.402743 | 1.73E-13 |
| MLXIP | 0.365609 | 3.91E-11 | S100A11 | 0.418016 | 6.11E-15 | ITPK1 | 0.401409 | 2.13E-13 |
| HCFC1 | 0.365294 | 4.09E-11 | PHLDB3 | 0.417431 | 6.71E-15 | KCNAB2 | 0.401327 | 2.13E-13 |
| TRIM32 | 0.365149 | 4.17E-11 | C6ORF136 | 0.416529 | 7.56E-15 | FEM1A | 0.401136 | 2.18E-13 |
| WNK2 | 0.364192 | 4.76E-11 | PEX11B | 0.415243 | 9.47E-15 | DNM2 | 0.400991 | 2.22E-13 |
| ATG2A | 0.362949 | 5.71E-11 | MRPL38 | 0.412838 | 1.42E-14 | CCDC22 | 0.400553 | 2.38E-13 |
| PATL1 | 0.362487 | 6.09E-11 | MED26 | 0.412574 | 1.48E-14 | TRMT1 | 0.400203 | 2.51E-13 |
| PML | 0.362137 | 6.38E-11 | EFNA4 | 0.412481 | 1.49E-14 | RPS15 | 0.399492 | 2.77E-13 |
| BRD4 | 0.361989 | 6.45E-11 | ACD | 0.412049 | 1.58E-14 | MBD3 | 0.399256 | 2.87E-13 |
| GDPD5 | 0.361987 | 6.45E-11 | TBC1D10B | 0.411291 | 1.78E-14 | AKAP8L | 0.39908 | 2.93E-13 |
| ULK3 | 0.361261 | 7.03E-11 | SETDB1 | 0.411061 | 1.84E-14 | RTN4R | 0.397971 | 3.43E-13 |
| SMPD2 | 0.361038 | 7.25E-11 | REXO1 | 0.41024 | 2.10E-14 | ECSIT | 0.397826 | 3.49E-13 |
| AP3S2 | 0.360672 | 7.58E-11 | LYPLA2 | 0.409154 | 2.50E-14 | GATAD2A | 0.397351 | 3.76E-13 |
| NCOA6 | 0.3604 | 7.88E-11 | PPP1R11 | 0.409077 | 2.53E-14 | RFX1 | 0.39708 | 3.92E-13 |
| SPTAN1 | 0.360148 | 8.16E-11 | HDGF | 0.408216 | 2.92E-14 | MAST3 | 0.396728 | 4.12E-13 |
| PLEKHO2 | 0.359863 | 8.43E-11 | POLR3C | 0.408172 | 2.92E-14 | MCRIP1 | 0.39671 | 4.12E-13 |
| ITPK1 | 0.359525 | 8.73E-11 | C19ORF47 | 0.407359 | 3.33E-14 | NDUFS7 | 0.396433 | 4.29E-13 |
| TANGO2 | 0.358708 | 9.82E-11 | NFKB2 | 0.407234 | 3.39E-14 | SLC27A1 | 0.396249 | 4.36E-13 |
| PNMA6A | 0.358352 | 1.03E-10 | CLK2 | 0.405699 | 4.32E-14 | CIAO3 | 0.396247 | 4.36E-13 |

| | | | | | | | | |
|---|---|---|---|---|---|---|---|---|
| CMIP | 0.35823 | 1.05E-10 | NFKBIL1 | 0.405055 | 4.76E-14 | SOX10 | 0.396137 | 4.42E-13 |
| FCMR | 0.358083 | 1.07E-10 | NOC4L | 0.404831 | 4.94E-14 | C19ORF71 | 0.395894 | 4.57E-13 |
| SLC7A5 | 0.358055 | 1.07E-10 | PHF1 | 0.403019 | 6.49E-14 | NUDT14 | 0.395427 | 4.89E-13 |
| ZNF609 | 0.357533 | 1.14E-10 | IGSF8 | 0.402778 | 6.75E-14 | RRP1 | 0.395143 | 5.10E-13 |
| TRIM26 | 0.357473 | 1.15E-10 | DAP3 | 0.402412 | 7.18E-14 | DMAP1 | 0.393756 | 6.36E-13 |
| NAV2 | 0.35707 | 1.21E-10 | TICAM1 | 0.40168 | 8.09E-14 | REEP6 | 0.393685 | 6.36E-13 |
| GDPGP1 | 0.356941 | 1.23E-10 | LAGE3 | 0.401675 | 8.09E-14 | SAFB2 | 0.393678 | 6.36E-13 |
| NDST2 | 0.356497 | 1.31E-10 | ARMC7 | 0.401173 | 8.78E-14 | TPGS1 | 0.393459 | 6.56E-13 |
| CTSH | 0.355574 | 1.47E-10 | DDX41 | 0.400993 | 9.04E-14 | KLHL22 | 0.392372 | 7.80E-13 |
| HPS1 | 0.355512 | 1.48E-10 | FAAP20 | 0.400216 | 1.02E-13 | MAP3K10 | 0.391937 | 8.29E-13 |
| HS3ST2 | 0.355324 | 1.51E-10 | CCDC9 | 0.400154 | 1.02E-13 | FAM53B | 0.391313 | 9.17E-13 |
| BCOR | 0.354461 | 1.70E-10 | ZGPAT | 0.39967 | 1.10E-13 | IRF2BP1 | 0.390459 | 1.05E-12 |
| OTUD7A | 0.354303 | 1.73E-10 | TJAP1 | 0.399665 | 1.10E-13 | TTLL1 | 0.390137 | 1.09E-12 |
| RAB5B | 0.353855 | 1.84E-10 | TMUB1 | 0.399374 | 1.14E-13 | RPL18A | 0.389709 | 1.15E-12 |
| SEMA6A | 0.353709 | 1.87E-10 | GPS2 | 0.399193 | 1.17E-13 | HMG20B | 0.389352 | 1.22E-12 |
| PPP1R26 | 0.35353 | 1.92E-10 | SDHAF1 | 0.398717 | 1.25E-13 | WDR13 | 0.389143 | 1.25E-12 |
| TMPRSS13 | 0.352722 | 2.15E-10 | TNIP2 | 0.398627 | 1.26E-13 | PIP5K1C | 0.388943 | 1.28E-12 |
| MAN2A2 | 0.350662 | 2.83E-10 | RFX1 | 0.397429 | 1.53E-13 | RAB11B | 0.388785 | 1.31E-12 |
| RASSF3 | 0.350266 | 2.97E-10 | CDK11B | 0.397256 | 1.57E-13 | CIC | 0.388637 | 1.33E-12 |
| SEPTIN4 | 0.35004 | 3.05E-10 | MRPL2 | 0.396155 | 1.84E-13 | SGF29 | 0.387225 | 1.67E-12 |
| GAB2 | 0.350039 | 3.05E-10 | ZBTB48 | 0.396022 | 1.87E-13 | AGPAT2 | 0.386867 | 1.74E-12 |
| LRCH3 | 0.349936 | 3.09E-10 | GBA | 0.395561 | 2.02E-13 | TBCB | 0.386845 | 1.74E-12 |
| PDE9A | 0.349883 | 3.11E-10 | ZNF628 | 0.395394 | 2.08E-13 | IPO13 | 0.386753 | 1.76E-12 |
| TLNRD1 | 0.34924 | 3.41E-10 | NUDC | 0.394346 | 2.46E-13 | RAVER1 | 0.386554 | 1.81E-12 |
| EPM2A | 0.349166 | 3.44E-10 | TRIM26 | 0.393385 | 2.85E-13 | NDUFA7 | 0.386222 | 1.89E-12 |
| C15ORF39 | 0.348303 | 3.82E-10 | SCAF1 | 0.39265 | 3.18E-13 | DEXI | 0.386139 | 1.91E-12 |
| ZBTB42 | 0.348298 | 3.82E-10 | POLL | 0.392598 | 3.20E-13 | UNC45A | 0.385504 | 2.11E-12 |

| | | | | | | | | |
|---|---|---|---|---|---|---|---|---|
| RCCD1 | 0.347555 | 4.17E-10 | PFDN2 | 0.392188 | 3.41E-13 | PTPRS | 0.385438 | 2.11E-12 |
| CDS2 | 0.347433 | 4.22E-10 | TESK1 | 0.392079 | 3.46E-13 | EVI5L | 0.385012 | 2.26E-12 |
| NAT8 | 0.347166 | 4.37E-10 | MCRIP1 | 0.392016 | 3.49E-13 | PPP1R26 | 0.384247 | 2.51E-12 |
| NCKAP5L | 0.346939 | 4.50E-10 | SLC66A1 | 0.391682 | 3.68E-13 | ARMC5 | 0.383898 | 2.63E-12 |
| SPG21 | 0.346634 | 4.67E-10 | GLMP | 0.391436 | 3.82E-13 | CTDP1 | 0.383659 | 2.72E-12 |
| EP400 | 0.346374 | 4.84E-10 | ZNF768 | 0.391133 | 3.99E-13 | ZNF205 | 0.383644 | 2.72E-12 |
| POU3F1 | 0.345951 | 5.11E-10 | DEDD2 | 0.39078 | 4.21E-13 | PLEKHM2 | 0.382994 | 3.01E-12 |
| QPCT | 0.344642 | 6.03E-10 | ADIPOR1 | 0.390259 | 4.53E-13 | TMEM259 | 0.382846 | 3.07E-12 |
| NHSL1 | 0.344312 | 6.26E-10 | PRPF3 | 0.39025 | 4.53E-13 | ARMC7 | 0.382738 | 3.11E-12 |
| ST20-AS1 | 0.343368 | 7.05E-10 | ZBTB17 | 0.390129 | 4.60E-13 | PCIF1 | 0.381888 | 3.55E-12 |
| DTNB | 0.343234 | 7.17E-10 | GPX4 | 0.389381 | 5.16E-13 | GADD45GIP1 | 0.381485 | 3.75E-12 |
| GATAD2A | 0.343003 | 7.36E-10 | ZBTB45 | 0.388695 | 5.77E-13 | HPS6 | 0.381117 | 3.97E-12 |
| USF2 | 0.342985 | 7.36E-10 | DMAP1 | 0.388414 | 5.99E-13 | PAK4 | 0.380943 | 4.06E-12 |
| CREBBP | 0.34279 | 7.55E-10 | ZNF513 | 0.388216 | 6.16E-13 | EIF3G | 0.380276 | 4.51E-12 |
| IQSEC1 | 0.342712 | 7.60E-10 | RAD23A | 0.387878 | 6.47E-13 | USP20 | 0.380171 | 4.55E-12 |
| MICAL1 | 0.342285 | 8.04E-10 | PTPN23 | 0.387781 | 6.55E-13 | ZNF524 | 0.380107 | 4.57E-12 |
| CCDC97 | 0.342259 | 8.05E-10 | WBP2 | 0.387071 | 7.34E-13 | ZNF341 | 0.380015 | 4.62E-12 |
| DNM2 | 0.341628 | 8.73E-10 | CYBC1 | 0.386965 | 7.46E-13 | DDA1 | 0.379496 | 4.98E-12 |
| KAZN | 0.34152 | 8.85E-10 | LRRC45 | 0.386482 | 8.03E-13 | MPND | 0.378123 | 6.15E-12 |
| SLC3A2 | 0.341325 | 9.05E-10 | IRF2BP1 | 0.38635 | 8.15E-13 | FAAP100 | 0.377347 | 6.95E-12 |
| FXYD6 | 0.341106 | 9.32E-10 | UFC1 | 0.38541 | 9.31E-13 | POLR2E | 0.377183 | 7.11E-12 |
| FLYWCH1 | 0.340847 | 9.65E-10 | GPS1 | 0.385043 | 9.81E-13 | PFKL | 0.376881 | 7.40E-12 |
| TYRP1 | 0.340757 | 9.75E-10 | SETD1A | 0.384427 | 1.07E-12 | ATP5F1D | 0.376541 | 7.77E-12 |
| ATP10A | 0.340391 | 1.01E-09 | ARHGDIA | 0.383731 | 1.19E-12 | MVB12A | 0.376519 | 7.77E-12 |
| LDB1 | 0.340128 | 1.04E-09 | EHMT2 | 0.383611 | 1.21E-12 | ATG4D | 0.376474 | 7.79E-12 |
| MYH9 | 0.339374 | 1.16E-09 | OGFR | 0.383322 | 1.26E-12 | REXO1 | 0.376009 | 8.33E-12 |
| INSM1 | 0.3393 | 1.17E-09 | NSUN5 | 0.383318 | 1.26E-12 | PHLDB3 | 0.375846 | 8.50E-12 |

| | | | | | | | | |
|---|---|---|---|---|---|---|---|---|
| ILRUN | 0.339216 | 1.18E-09 | BRAT1 | 0.383257 | 1.27E-12 | ASIC3 | 0.375718 | 8.61E-12 |
| TECPR2 | 0.339168 | 1.18E-09 | EIF3G | 0.382633 | 1.38E-12 | BLOC1S3 | 0.374226 | 1.07E-11 |
| PRR5-ARHGAP8 | 0.338953 | 1.22E-09 | BCL7C | 0.382615 | 1.38E-12 | GTF2F1 | 0.373945 | 1.12E-11 |
| DENND1A | 0.338727 | 1.25E-09 | PSENEN | 0.382579 | 1.39E-12 | SAFB | 0.373817 | 1.14E-11 |
| SPSB1 | 0.338157 | 1.34E-09 | GAK | 0.382573 | 1.39E-12 | MECP2 | 0.372582 | 1.36E-11 |
| CIZ1 | 0.337999 | 1.36E-09 | INTS3 | 0.382533 | 1.39E-12 | COPE | 0.37254 | 1.36E-11 |
| TATDN2 | 0.337443 | 1.46E-09 | FBXW5 | 0.382457 | 1.41E-12 | CLCN7 | 0.37243 | 1.38E-11 |
| SMAD3 | 0.337163 | 1.52E-09 | SH3BP5L | 0.382357 | 1.43E-12 | STK32C | 0.371616 | 1.55E-11 |
| FAT2 | 0.337025 | 1.54E-09 | ZDHHC18 | 0.381855 | 1.54E-12 | ZNF213 | 0.371541 | 1.56E-11 |
| RRAGD | 0.336884 | 1.56E-09 | PRPF6 | 0.381822 | 1.54E-12 | AKT2 | 0.371414 | 1.59E-11 |
| IRX3 | 0.336691 | 1.60E-09 | HCFC1R1 | 0.381456 | 1.62E-12 | MIER2 | 0.371199 | 1.63E-11 |
| LRRC3 | 0.336471 | 1.65E-09 | ADRM1 | 0.380313 | 1.93E-12 | KLF16 | 0.371096 | 1.65E-11 |
| DCAF11 | 0.33639 | 1.67E-09 | TRIR | 0.379768 | 2.09E-12 | SUPT5H | 0.370252 | 1.86E-11 |
| HOMEZ | 0.336252 | 1.69E-09 | ARHGEF11 | 0.379646 | 2.12E-12 | HOMER3 | 0.370121 | 1.88E-11 |
| USP20 | 0.336181 | 1.70E-09 | PCBP1 | 0.37961 | 2.13E-12 | MCOLN1 | 0.369414 | 2.07E-11 |
| SLC9A3R1 | 0.336015 | 1.73E-09 | SAFB2 | 0.379534 | 2.15E-12 | ARMC6 | 0.36927 | 2.10E-11 |
| TRIM63 | 0.335369 | 1.85E-09 | HCN3 | 0.37938 | 2.19E-12 | HDGFL2 | 0.368733 | 2.27E-11 |
| CABLES1 | 0.334815 | 1.99E-09 | RFNG | 0.379126 | 2.28E-12 | SCAF4 | 0.368062 | 2.52E-11 |
| NATD1 | 0.334338 | 2.11E-09 | LRCH4 | 0.378622 | 2.45E-12 | MED24 | 0.368014 | 2.52E-11 |
| RUBCN | 0.334054 | 2.18E-09 | HPS6 | 0.378448 | 2.51E-12 | BSG | 0.368007 | 2.52E-11 |
| EPG5 | 0.333792 | 2.23E-09 | C7ORF26 | 0.377749 | 2.76E-12 | UBA52 | 0.367928 | 2.52E-11 |
| CCNK | 0.333518 | 2.31E-09 | SSU72 | 0.377475 | 2.86E-12 | IP6K1 | 0.367801 | 2.56E-11 |
| AMBRA1 | 0.333092 | 2.45E-09 | CSNK2B | 0.377302 | 2.93E-12 | RPL28 | 0.36779 | 2.56E-11 |
| SETD1B | 0.33244 | 2.65E-09 | OTUD5 | 0.377247 | 2.95E-12 | BORCS8-MEF2B | 0.367674 | 2.57E-11 |
| CTSD | 0.332068 | 2.78E-09 | FBRS | 0.376916 | 3.09E-12 | SDR39U1 | 0.367657 | 2.57E-11 |
| TFF3 | 0.331946 | 2.82E-09 | SF3A2 | 0.376869 | 3.11E-12 | RENBP | 0.367657 | 2.57E-11 |
| SRCAP | 0.33169 | 2.92E-09 | TMC6 | 0.376739 | 3.16E-12 | POLR2I | 0.367247 | 2.72E-11 |

| | | | | | | | | |
|---|---|---|---|---|---|---|---|---|
| RNF31 | 0.331342 | 3.04E-09 | ISG20L2 | 0.376348 | 3.35E-12 | SF3A2 | 0.367009 | 2.81E-11 |
| UBE2O | 0.331023 | 3.17E-09 | ZNF668 | 0.375811 | 3.60E-12 | KCTD13 | 0.366429 | 3.06E-11 |
| CTSF | 0.330987 | 3.18E-09 | PTPN18 | 0.375761 | 3.61E-12 | KMT2B | 0.366397 | 3.07E-11 |
| PIK3C2B | 0.330849 | 3.23E-09 | CNOT3 | 0.375536 | 3.74E-12 | SLC3A2 | 0.366292 | 3.11E-11 |
| SLC37A1 | 0.33062 | 3.32E-09 | CTDP1 | 0.375161 | 3.93E-12 | PPAN | 0.36624 | 3.12E-11 |
| PGM5 | 0.33022 | 3.50E-09 | TIMM17B | 0.374099 | 4.57E-12 | ZSWIM1 | 0.365516 | 3.42E-11 |
| OLFM1 | 0.330153 | 3.52E-09 | KMT2B | 0.374043 | 4.59E-12 | AXIN1 | 0.365456 | 3.44E-11 |
| KDM2B | 0.330072 | 3.55E-09 | CDC42SE1 | 0.373839 | 4.74E-12 | ELL | 0.36543 | 3.45E-11 |
| TBC1D16 | 0.329801 | 3.65E-09 | RAVER1 | 0.3738 | 4.76E-12 | IFT140 | 0.365201 | 3.55E-11 |
| MAP2K5 | 0.329603 | 3.74E-09 | NENF | 0.373365 | 5.07E-12 | ARHGDIA | 0.365128 | 3.57E-11 |
| FSTL4 | 0.3294 | 3.82E-09 | TRAF2 | 0.373107 | 5.26E-12 | EEF2 | 0.365111 | 3.57E-11 |
| ADCY2 | 0.32898 | 4.03E-09 | GDI1 | 0.373019 | 5.31E-12 | KDM4B | 0.364245 | 4.08E-11 |
| UCK1 | 0.328811 | 4.10E-09 | AKAP8L | 0.372983 | 5.33E-12 | LHPP | 0.363986 | 4.21E-11 |
| LINC00906 | 0.328462 | 4.27E-09 | RNF25 | 0.372951 | 5.34E-12 | TMEM143 | 0.363893 | 4.26E-11 |
| PTPN23 | 0.328024 | 4.50E-09 | IMP4 | 0.372676 | 5.54E-12 | CARM1 | 0.363714 | 4.37E-11 |
| SERPINA12 | 0.32756 | 4.77E-09 | ZNF593 | 0.372024 | 6.10E-12 | LONP1 | 0.363622 | 4.41E-11 |
| RAI1 | 0.327395 | 4.87E-09 | CUTA | 0.372023 | 6.10E-12 | ABCD1 | 0.3635 | 4.47E-11 |
| BEST1 | 0.327064 | 5.06E-09 | ZNF511 | 0.371397 | 6.68E-12 | DCAF11 | 0.362175 | 5.43E-11 |
| CDH1 | 0.326851 | 5.20E-09 | C8ORF58 | 0.370222 | 7.94E-12 | SLC25A23 | 0.362051 | 5.50E-11 |
| TBC1D9B | 0.326586 | 5.36E-09 | ATP6V0B | 0.370203 | 7.94E-12 | SMARCA4 | 0.362047 | 5.50E-11 |
| RABL6 | 0.326497 | 5.41E-09 | NR1H2 | 0.369707 | 8.55E-12 | MFSD12 | 0.362031 | 5.50E-11 |
| MRTFA | 0.326391 | 5.46E-09 | UBXN11 | 0.369395 | 8.94E-12 | DPP7 | 0.361529 | 5.89E-11 |
| MAL | 0.326158 | 5.59E-09 | GSK3A | 0.369237 | 9.13E-12 | DBP | 0.361377 | 6.01E-11 |
| ZER1 | 0.326003 | 5.69E-09 | GRAMD1A | 0.369231 | 9.13E-12 | GDF1 | 0.361105 | 6.21E-11 |
| PPM1H | 0.325943 | 5.72E-09 | PAXX | 0.368695 | 9.81E-12 | HPS1 | 0.36099 | 6.28E-11 |
| STARD10 | 0.325816 | 5.81E-09 | COG1 | 0.368633 | 9.85E-12 | FDX2 | 0.360713 | 6.53E-11 |
| CEP250 | 0.325685 | 5.90E-09 | CSK | 0.368308 | 1.03E-11 | PNMA6A | 0.360655 | 6.57E-11 |

| | | | | | | | | |
|---|---|---|---|---|---|---|---|---|
| PELI3 | 0.325291 | 6.20E-09 | THAP4 | 0.368199 | 1.04E-11 | ZNF581 | 0.360065 | 7.17E-11 |
| MYO1D | 0.324754 | 6.63E-09 | TREX1 | 0.368196 | 1.04E-11 | OTUD7A | 0.359919 | 7.31E-11 |
| KIT | 0.324572 | 6.73E-09 | RGS14 | 0.368173 | 1.05E-11 | ST3GAL3 | 0.359518 | 7.75E-11 |
| EPB41 | 0.324322 | 6.95E-09 | SMG7 | 0.368111 | 1.05E-11 | BAP1 | 0.359492 | 7.75E-11 |
| AMER2 | 0.324294 | 6.96E-09 | SNX8 | 0.368017 | 1.07E-11 | BAIAP2 | 0.359118 | 8.11E-11 |
| LOC148709 | 0.324213 | 7.02E-09 | DUS1L | 0.367767 | 1.10E-11 | SETD1A | 0.359087 | 8.12E-11 |
| EVC | 0.324207 | 7.02E-09 | INTS11 | 0.367551 | 1.13E-11 | SARS2 | 0.358959 | 8.26E-11 |
| CLCN7 | 0.323147 | 8.01E-09 | PYCR2 | 0.367529 | 1.13E-11 | AATBC | 0.35863 | 8.63E-11 |
| CNPPD1 | 0.323117 | 8.03E-09 | FKBPL | 0.367425 | 1.15E-11 | SNAPC2 | 0.358102 | 9.27E-11 |
| ARSG | 0.322918 | 8.21E-09 | MZT2B | 0.367259 | 1.17E-11 | TMEM187 | 0.358054 | 9.31E-11 |
| TTC7B | 0.322801 | 8.31E-09 | TNK2 | 0.36708 | 1.20E-11 | QTRT1 | 0.357969 | 9.40E-11 |
| KCNIP3 | 0.322296 | 8.82E-09 | EXOSC1 | 0.366775 | 1.26E-11 | PQBP1 | 0.357828 | 9.58E-11 |
| KMT2B | 0.322123 | 8.99E-09 | ADPRHL2 | 0.36657 | 1.30E-11 | TMEM191A | 0.357362 | 1.02E-10 |
| WDTC1 | 0.322031 | 9.09E-09 | SAP30BP | 0.366037 | 1.38E-11 | RECQL5 | 0.357351 | 1.02E-10 |
| SLC16A6 | 0.321998 | 9.12E-09 | RHBDF2 | 0.36566 | 1.46E-11 | SYMPK | 0.357203 | 1.04E-10 |
| PACS2 | 0.321778 | 9.37E-09 | ENSA | 0.365145 | 1.58E-11 | CEP170B | 0.356841 | 1.10E-10 |
| DUSP8 | 0.321491 | 9.68E-09 | PEA15 | 0.365076 | 1.59E-11 | AKT1 | 0.356836 | 1.10E-10 |
| OTX1 | 0.321475 | 9.68E-09 | NOSIP | 0.36488 | 1.63E-11 | POLRMT | 0.356679 | 1.12E-10 |
| MBP | 0.320953 | 1.03E-08 | FDPS | 0.364375 | 1.76E-11 | CEP131 | 0.356651 | 1.12E-10 |
| AFF3 | 0.320824 | 1.04E-08 | ZNF787 | 0.364186 | 1.80E-11 | ZNF362 | 0.356637 | 1.12E-10 |
| SIK1 | 0.320694 | 1.05E-08 | RNPEPL1 | 0.36411 | 1.82E-11 | LIN37 | 0.356608 | 1.12E-10 |
| CHRFAM7A | 0.3205 | 1.08E-08 | MYPOP | 0.364101 | 1.82E-11 | CSNK1D | 0.356601 | 1.12E-10 |
| SIGLEC8 | 0.320109 | 1.13E-08 | EDC4 | 0.363506 | 1.97E-11 | TBCD | 0.356266 | 1.17E-10 |
| HIP1R | 0.319899 | 1.16E-08 | NCKAP5L | 0.363124 | 2.07E-11 | NELFB | 0.355978 | 1.21E-10 |
| LRRK1 | 0.319742 | 1.17E-08 | MAP7D1 | 0.363099 | 2.08E-11 | DGCR6L | 0.35596 | 1.21E-10 |
| RPS6KA4 | 0.319728 | 1.17E-08 | MRPL43 | 0.362999 | 2.10E-11 | NKPD1 | 0.35594 | 1.21E-10 |
| ABCC2 | 0.319693 | 1.18E-08 | TMEM115 | 0.362982 | 2.10E-11 | TYSND1 | 0.355932 | 1.21E-10 |

| | | | | | | | | |
|---|---|---|---|---|---|---|---|---|
| TGFBRAP1 | 0.319534 | 1.20E-08 | LIX1L | 0.362289 | 2.32E-11 | SSBP4 | 0.355749 | 1.24E-10 |
| ZNF142 | 0.318792 | 1.31E-08 | ACTB | 0.362002 | 2.39E-11 | WDTC1 | 0.355639 | 1.26E-10 |
| BIK | 0.318787 | 1.31E-08 | C1ORF43 | 0.361816 | 2.44E-11 | ZNF646 | 0.355635 | 1.26E-10 |
| ZNF672 | 0.318557 | 1.35E-08 | PWWP2B | 0.361812 | 2.44E-11 | COQ4 | 0.355527 | 1.27E-10 |
| SMG1P3 | 0.317977 | 1.45E-08 | ACBD6 | 0.361456 | 2.56E-11 | USE1 | 0.355393 | 1.29E-10 |
| GOLGA7B | 0.317941 | 1.45E-08 | RHOG | 0.361238 | 2.64E-11 | TAF6L | 0.355349 | 1.29E-10 |
| ABCD1 | 0.31766 | 1.50E-08 | XAB2 | 0.361153 | 2.67E-11 | MAP3K6 | 0.355337 | 1.29E-10 |
| TMEM94 | 0.317613 | 1.51E-08 | USF2 | 0.36093 | 2.76E-11 | ZNF768 | 0.3549 | 1.37E-10 |
| KCTD21 | 0.317592 | 1.51E-08 | SCO2 | 0.360869 | 2.78E-11 | MIEF2 | 0.354852 | 1.38E-10 |
| SUV39H1 | 0.3175 | 1.52E-08 | TAGLN2 | 0.360775 | 2.80E-11 | MLST8 | 0.354465 | 1.45E-10 |
| MGRN1 | 0.31747 | 1.53E-08 | MRPL41 | 0.360615 | 2.86E-11 | STRADA | 0.354305 | 1.48E-10 |
| TECPR1 | 0.31745 | 1.53E-08 | RAB40C | 0.360527 | 2.89E-11 | TNPO2 | 0.353861 | 1.57E-10 |
| SNX29 | 0.31733 | 1.55E-08 | ASPSCR1 | 0.360495 | 2.90E-11 | APBA3 | 0.353408 | 1.68E-10 |
| KIF26A | 0.317211 | 1.57E-08 | LMNA | 0.360437 | 2.91E-11 | HDAC11 | 0.353357 | 1.69E-10 |
| ASB13 | 0.316823 | 1.64E-08 | MTMR14 | 0.359678 | 3.24E-11 | ZBTB42 | 0.35312 | 1.73E-10 |
| DCAF7 | 0.315903 | 1.84E-08 | ANXA11 | 0.35963 | 3.26E-11 | TOLLIP | 0.353093 | 1.74E-10 |
| KRTAP5-AS1 | 0.315811 | 1.85E-08 | INO80E | 0.359603 | 3.27E-11 | LSS | 0.352918 | 1.78E-10 |
| IL6R | 0.315467 | 1.93E-08 | BAG6 | 0.359292 | 3.42E-11 | ABHD4 | 0.352169 | 1.97E-10 |
| FGD1 | 0.315022 | 2.03E-08 | MVB12A | 0.359223 | 3.45E-11 | CHMP6 | 0.352116 | 1.98E-10 |
| IMP3 | 0.314654 | 2.12E-08 | C6ORF47 | 0.359139 | 3.49E-11 | TBL3 | 0.352069 | 1.99E-10 |
| DVL3 | 0.31461 | 2.13E-08 | CCDC71 | 0.358794 | 3.67E-11 | ASNA1 | 0.351916 | 2.03E-10 |
| ZFP92 | 0.314404 | 2.18E-08 | LRFN3 | 0.358597 | 3.77E-11 | ABHD17A | 0.351849 | 2.04E-10 |
| ST6GALNAC1 | 0.314196 | 2.22E-08 | PIK3CD | 0.358304 | 3.93E-11 | CYB561A3 | 0.350978 | 2.30E-10 |
| RBPMS | 0.313457 | 2.41E-08 | CCDC107 | 0.35828 | 3.94E-11 | CNPPD1 | 0.350774 | 2.37E-10 |
| SYDE1 | 0.313218 | 2.48E-08 | PXDC1 | 0.358093 | 4.03E-11 | ZNF687 | 0.350692 | 2.39E-10 |
| FBRSL1 | 0.312758 | 2.61E-08 | ARHGEF1 | 0.357812 | 4.20E-11 | RAB3A | 0.350554 | 2.41E-10 |
| HPS6 | 0.312601 | 2.65E-08 | BICRA | 0.357623 | 4.31E-11 | EEFSEC | 0.350363 | 2.46E-10 |

| | | | | | | | | |
|---|---|---|---|---|---|---|---|---|
| MAD1L1 | 0.312557 | 2.66E-08 | MGST3 | 0.357333 | 4.48E-11 | LZTS1 | 0.350307 | 2.48E-10 |
| ZMYM3 | 0.312267 | 2.76E-08 | RAB13 | 0.357194 | 4.57E-11 | TRIR | 0.350119 | 2.54E-10 |
| ZFYVE1 | 0.312185 | 2.78E-08 | C2CD4D | 0.357138 | 4.60E-11 | SLC66A2 | 0.349818 | 2.64E-10 |
| DCLK1 | 0.311819 | 2.90E-08 | ZNF653 | 0.356956 | 4.72E-11 | JOSD2 | 0.349272 | 2.85E-10 |
| ANKRD9 | 0.311729 | 2.93E-08 | RBM38 | 0.356796 | 4.83E-11 | GTPBP3 | 0.349202 | 2.87E-10 |
| FLNA | 0.311664 | 2.94E-08 | DALRD3 | 0.356716 | 4.87E-11 | F8A1 | 0.349152 | 2.88E-10 |
| DUSP16 | 0.311207 | 3.09E-08 | ADAM15 | 0.356449 | 5.04E-11 | RPL36 | 0.34909 | 2.90E-10 |
| TINCR | 0.310765 | 3.26E-08 | SAMD10 | 0.355899 | 5.43E-11 | MGRN1 | 0.348859 | 2.99E-10 |
| GPR157 | 0.310546 | 3.35E-08 | IER5 | 0.355324 | 5.85E-11 | LIPE | 0.347961 | 3.39E-10 |
| CASZ1 | 0.310338 | 3.43E-08 | MAD1L1 | 0.355084 | 6.05E-11 | PGAP3 | 0.347742 | 3.48E-10 |
| TSPAN10 | 0.310319 | 3.43E-08 | STK32C | 0.355054 | 6.07E-11 | TMEM160 | 0.347622 | 3.53E-10 |
| COX5A | 0.31028 | 3.44E-08 | SLX1B | 0.354986 | 6.12E-11 | BABAM1 | 0.347538 | 3.56E-10 |
| TLE3 | 0.310276 | 3.44E-08 | AURKAIP1 | 0.354888 | 6.19E-11 | DYRK1B | 0.347097 | 3.76E-10 |
| HES6 | 0.310243 | 3.45E-08 | HLX | 0.354828 | 6.24E-11 | UCK1 | 0.346852 | 3.87E-10 |
| PRRC2B | 0.309968 | 3.56E-08 | ARNT | 0.354708 | 6.34E-11 | DEAF1 | 0.346811 | 3.88E-10 |
| CA14 | 0.309286 | 3.84E-08 | RBM42 | 0.354368 | 6.64E-11 | NRL | 0.346566 | 4.01E-10 |
| TTYH2 | 0.309189 | 3.88E-08 | PPP1R35 | 0.354319 | 6.68E-11 | PARS2 | 0.346552 | 4.01E-10 |
| MAP3K3 | 0.308477 | 4.18E-08 | TBCC | 0.353702 | 7.25E-11 | SDHAF1 | 0.346357 | 4.10E-10 |
| FBXW5 | 0.308247 | 4.29E-08 | PRUNE1 | 0.353546 | 7.39E-11 | FBXW5 | 0.346344 | 4.10E-10 |
| KMT2D | 0.308101 | 4.35E-08 | ZFAND2B | 0.35328 | 7.66E-11 | PRKACA | 0.346306 | 4.11E-10 |
| PIP4K2A | 0.30795 | 4.40E-08 | RNF40 | 0.353192 | 7.74E-11 | CPTP | 0.345709 | 4.46E-10 |
| NR4A3 | 0.307704 | 4.52E-08 | SH2B2 | 0.35299 | 7.95E-11 | MRPL4 | 0.345649 | 4.47E-10 |
| RETREG2 | 0.307676 | 4.53E-08 | LEMD2 | 0.352851 | 8.06E-11 | NDUFA13 | 0.34562 | 4.47E-10 |
| SLC6A8 | 0.307621 | 4.55E-08 | MUTYH | 0.352725 | 8.19E-11 | PPP1R12C | 0.345531 | 4.51E-10 |
| PRR12 | 0.307017 | 4.88E-08 | XYLT2 | 0.352707 | 8.20E-11 | MGAT5B | 0.345318 | 4.62E-10 |
| EPHX1 | 0.306901 | 4.95E-08 | TMEM79 | 0.352622 | 8.29E-11 | TTYH2 | 0.345243 | 4.66E-10 |
| TTYH3 | 0.306341 | 5.28E-08 | NELFE | 0.352581 | 8.33E-11 | OR7A5 | 0.345133 | 4.72E-10 |

| | | | | | | | | |
|---|---|---|---|---|---|---|---|---|
| ARRB1 | 0.306282 | 5.31E-08 | ZNF691 | 0.352107 | 8.88E-11 | PEX11G | 0.3451 | 4.73E-10 |
| ELMSAN1 | 0.306219 | 5.34E-08 | ABT1 | 0.352089 | 8.88E-11 | RPS9 | 0.344692 | 4.97E-10 |
| NUMA1 | 0.305339 | 5.90E-08 | KIAA2013 | 0.351623 | 9.45E-11 | ITPKB | 0.34446 | 5.13E-10 |
| SPATA41 | 0.304742 | 6.31E-08 | RITA1 | 0.351327 | 9.83E-11 | NAB2 | 0.343817 | 5.59E-10 |
| MYO18A | 0.304082 | 6.80E-08 | SNAPC4 | 0.350631 | 1.08E-10 | RRNAD1 | 0.343678 | 5.68E-10 |
| GATAD2B | 0.304034 | 6.83E-08 | LYPLA2P1 | 0.350626 | 1.08E-10 | PEMT | 0.343668 | 5.68E-10 |
| FNBP1 | 0.303258 | 7.43E-08 | PSMC5 | 0.350572 | 1.09E-10 | FBXL15 | 0.343368 | 5.91E-10 |
| RPTOR | 0.302909 | 7.73E-08 | GALK1 | 0.349734 | 1.22E-10 | PRG2 | 0.343218 | 6.02E-10 |
| PITRM1 | 0.302559 | 8.02E-08 | ADAR | 0.349395 | 1.27E-10 | PEPD | 0.343208 | 6.02E-10 |
| HLA-F-AS1 | 0.302495 | 8.08E-08 | FBRSL1 | 0.349356 | 1.28E-10 | CLPP | 0.342776 | 6.40E-10 |
| RNF220 | 0.302294 | 8.26E-08 | APEX2 | 0.34881 | 1.38E-10 | 2-Mar | 0.342524 | 6.59E-10 |
| PACS1 | 0.302075 | 8.48E-08 | UBE2J2 | 0.34876 | 1.39E-10 | PRODH | 0.34239 | 6.67E-10 |
| TBC1D13 | 0.30182 | 8.68E-08 | DDA1 | 0.348425 | 1.44E-10 | POMT1 | 0.342172 | 6.83E-10 |
| DYNLL2 | 0.301806 | 8.68E-08 | RNF5 | 0.348416 | 1.44E-10 | PEX14 | 0.342165 | 6.83E-10 |
| NEDD4L | 0.301694 | 8.76E-08 | TMEM39B | 0.348141 | 1.49E-10 | AVPI1 | 0.342131 | 6.85E-10 |
| CHRNA7 | 0.301595 | 8.84E-08 | EMD | 0.34782 | 1.56E-10 | TFF3 | 0.342081 | 6.88E-10 |
| WHAMM | 0.301488 | 8.95E-08 | RPUSD2 | 0.347583 | 1.61E-10 | WDR83 | 0.342003 | 6.95E-10 |
| UBOX5 | 0.301257 | 9.17E-08 | MZT2A | 0.347535 | 1.62E-10 | UBALD1 | 0.341655 | 7.27E-10 |
| HEXA | 0.301196 | 9.23E-08 | SSNA1 | 0.347367 | 1.66E-10 | TRAPPC12 | 0.341605 | 7.31E-10 |
| THRAP3 | 0.300306 | 1.02E-07 | PPP4C | 0.347239 | 1.69E-10 | PIP4P1 | 0.34147 | 7.44E-10 |
| TKT | 0.30025 | 1.03E-07 | EIF2B4 | 0.347162 | 1.70E-10 | LZTS2 | 0.341433 | 7.46E-10 |
| GMPR | 0.300107 | 1.05E-07 | RAB3A | 0.347001 | 1.74E-10 | FBRSL1 | 0.341391 | 7.48E-10 |
| PRXL2B | 0.300095 | 1.05E-07 | EPN1 | 0.346642 | 1.83E-10 | TMEM250 | 0.34117 | 7.68E-10 |
| SNX12 | 0.300011 | 1.05E-07 | ZFYVE19 | 0.346302 | 1.92E-10 | XRCC1 | 0.341164 | 7.68E-10 |
| | | | EEFSEC | 0.346271 | 1.92E-10 | GPATCH3 | 0.341032 | 7.78E-10 |
| | | | TSSC4 | 0.346245 | 1.93E-10 | STK11 | 0.340968 | 7.83E-10 |
| | | | ZNF444 | 0.346218 | 1.93E-10 | NOP53 | 0.340905 | 7.89E-10 |

| | | | | | | | | |
|---|---|---|---|---|---|---|---|---|
| | | | BLOC1S3 | 0.346023 | 1.98E-10 | TIMM44 | 0.340533 | 8.30E-10 |
| | | | UBL7 | 0.345761 | 2.06E-10 | ARSE | 0.340157 | 8.68E-10 |
| | | | MEA1 | 0.34538 | 2.16E-10 | THRA | 0.339749 | 9.12E-10 |
| | | | TPRN | 0.345303 | 2.18E-10 | MICOS13 | 0.339017 | 1.01E-09 |
| | | | FAM50A | 0.345278 | 2.18E-10 | INF2 | 0.338521 | 1.07E-09 |
| | | | ACTR1A | 0.345124 | 2.22E-10 | MNT | 0.338514 | 1.07E-09 |
| | | | OCEL1 | 0.34498 | 2.27E-10 | HSPBP1 | 0.338319 | 1.09E-09 |
| | | | ZNF296 | 0.344945 | 2.27E-10 | GALK1 | 0.338319 | 1.09E-09 |
| | | | UBALD2 | 0.344746 | 2.34E-10 | DUS3L | 0.33827 | 1.10E-09 |
| | | | EGLN2 | 0.344626 | 2.37E-10 | GCDH | 0.338171 | 1.11E-09 |
| | | | LENG1 | 0.344512 | 2.40E-10 | WDR81 | 0.338157 | 1.11E-09 |
| | | | CLN3 | 0.344497 | 2.41E-10 | LINC00999 | 0.337977 | 1.14E-09 |
| | | | SIRT6 | 0.344302 | 2.46E-10 | TMED1 | 0.337792 | 1.16E-09 |
| | | | ZC3H3 | 0.344088 | 2.54E-10 | KAT8 | 0.337573 | 1.20E-09 |
| | | | MED25 | 0.343987 | 2.57E-10 | BGLAP | 0.337296 | 1.24E-09 |
| | | | RBM10 | 0.343318 | 2.81E-10 | PIP4K2B | 0.337208 | 1.25E-09 |
| | | | EFNA3 | 0.343024 | 2.92E-10 | DIPK1B | 0.337102 | 1.26E-09 |
| | | | EDF1 | 0.342935 | 2.96E-10 | FAM71E1 | 0.337083 | 1.27E-09 |
| | | | POGZ | 0.342782 | 3.02E-10 | RPUSD2 | 0.336807 | 1.31E-09 |
| | | | DNAJC4 | 0.342718 | 3.03E-10 | ASB6 | 0.336069 | 1.44E-09 |
| | | | PLCD1 | 0.342502 | 3.10E-10 | ALDH16A1 | 0.336064 | 1.44E-09 |
| | | | APBA3 | 0.342232 | 3.21E-10 | ATG2A | 0.335963 | 1.46E-09 |
| | | | KLF16 | 0.342195 | 3.22E-10 | CCDC159 | 0.335657 | 1.51E-09 |
| | | | ITPKB | 0.341969 | 3.32E-10 | MED16 | 0.335262 | 1.59E-09 |
| | | | ABCF3 | 0.341826 | 3.38E-10 | UBAP1L | 0.334708 | 1.71E-09 |
| | | | NDOR1 | 0.341534 | 3.52E-10 | CELSR2 | 0.334379 | 1.78E-09 |
| | | | TAZ | 0.341424 | 3.57E-10 | GTPBP6 | 0.33433 | 1.79E-09 |

| | | | | | | | | |
|---|---|---|---|---|---|---|---|---|
| | | | ELOF1 | 0.341093 | 3.73E-10 | KCNIP3 | 0.334283 | 1.80E-09 |
| | | | RETREG2 | 0.341073 | 3.74E-10 | LENG1 | 0.334272 | 1.80E-09 |
| | | | AP5Z1 | 0.341007 | 3.77E-10 | HGS | 0.334127 | 1.83E-09 |
| | | | DCAF8 | 0.341005 | 3.77E-10 | MAD1L1 | 0.333896 | 1.88E-09 |
| | | | MFSD13A | 0.340285 | 4.14E-10 | C11ORF68 | 0.333816 | 1.90E-09 |
| | | | E4F1 | 0.340176 | 4.18E-10 | UNK | 0.333615 | 1.94E-09 |
| | | | PHKG2 | 0.339973 | 4.29E-10 | PIAS4 | 0.333507 | 1.96E-09 |
| | | | VPS16 | 0.339968 | 4.29E-10 | VAT1 | 0.33345 | 1.97E-09 |
| | | | PAF1 | 0.339597 | 4.50E-10 | ZNF579 | 0.333186 | 2.04E-09 |
| | | | ESS2 | 0.339144 | 4.78E-10 | PAF1 | 0.333046 | 2.08E-09 |
| | | | SCAND1 | 0.338944 | 4.90E-10 | ERI3 | 0.332955 | 2.10E-09 |
| | | | AGPAT1 | 0.338842 | 4.96E-10 | ZNF688 | 0.33286 | 2.12E-09 |
| | | | ZNF213 | 0.338835 | 4.96E-10 | NCKAP5L | 0.332535 | 2.21E-09 |
| | | | GRK6 | 0.338634 | 5.09E-10 | RABAC1 | 0.332248 | 2.28E-09 |
| | | | GPN2 | 0.338589 | 5.12E-10 | RPL10 | 0.332091 | 2.32E-09 |
| | | | ZNF579 | 0.338578 | 5.12E-10 | MRPL38 | 0.332052 | 2.33E-09 |
| | | | TMCC2 | 0.33854 | 5.15E-10 | TAOK2 | 0.331692 | 2.44E-09 |
| | | | CCM2 | 0.33841 | 5.21E-10 | SNAPC4 | 0.331619 | 2.46E-09 |
| | | | CUEDC2 | 0.337963 | 5.52E-10 | AP5B1 | 0.331605 | 2.46E-09 |
| | | | MAPK3 | 0.337902 | 5.57E-10 | MECR | 0.331572 | 2.46E-09 |
| | | | BANP | 0.337876 | 5.58E-10 | RPL18 | 0.331457 | 2.49E-09 |
| | | | IRF3 | 0.337771 | 5.65E-10 | MXD4 | 0.331314 | 2.53E-09 |
| | | | FBXO46 | 0.337635 | 5.74E-10 | STRN4 | 0.330949 | 2.65E-09 |
| | | | H1FX | 0.337628 | 5.74E-10 | NMRAL1 | 0.330487 | 2.81E-09 |
| | | | SAFB | 0.33758 | 5.77E-10 | CIZ1 | 0.330411 | 2.83E-09 |
| | | | TRAPPC12 | 0.33752 | 5.81E-10 | | | |
| | | | NELFA | 0.337111 | 6.10E-10 | | | |

| | | | | | |
|---|---|---|---|---|---|
| | | | NAB2 | 0.336883 | 6.28E-10 |
| | | | NDUFB11 | 0.336638 | 6.47E-10 |
| | | | GEMIN7 | 0.336435 | 6.63E-10 |
| | | | ZNF335 | 0.336398 | 6.65E-10 |
| | | | SNRPA | 0.336393 | 6.65E-10 |
| | | | CLK3 | 0.336355 | 6.68E-10 |
| | | | SLC25A28 | 0.336343 | 6.69E-10 |
| | | | SPSB3 | 0.336298 | 6.71E-10 |
| | | | LZTS2 | 0.335966 | 7.01E-10 |
| | | | RNF31 | 0.335962 | 7.01E-10 |
| | | | DAPK3 | 0.335906 | 7.05E-10 |
| | | | ZNF446 | 0.335866 | 7.08E-10 |
| | | | TRIM8 | 0.335756 | 7.18E-10 |
| | | | DPP7 | 0.335568 | 7.34E-10 |
| | | | CC2D1B | 0.33543 | 7.47E-10 |
| | | | DAGLB | 0.335317 | 7.57E-10 |
| | | | UBA52 | 0.335155 | 7.73E-10 |
| | | | UNC93B1 | 0.335132 | 7.75E-10 |
| | | | SLC35C2 | 0.334984 | 7.90E-10 |
| | | | MARK4 | 0.334927 | 7.96E-10 |
| | | | PRRC2A | 0.33479 | 8.09E-10 |
| | | | ARPC4 | 0.334603 | 8.28E-10 |
| | | | PQBP1 | 0.334486 | 8.40E-10 |
| | | | PPP6R1 | 0.334471 | 8.41E-10 |
| | | | MGAT5B | 0.334321 | 8.58E-10 |
| | | | AP1M1 | 0.334021 | 8.89E-10 |
| | | | TFEB | 0.333988 | 8.92E-10 |

| | | | | | |
|---|---|---|---|---|---|
| | | | HSD3B7 | 0.33392 | 8.99E-10 |
| | | | ZNF205 | 0.333819 | 9.11E-10 |
| | | | ABCD1 | 0.333706 | 9.24E-10 |
| | | | PEX14 | 0.3336 | 9.37E-10 |
| | | | MOB2 | 0.333457 | 9.55E-10 |
| | | | JUND | 0.33304 | 1.01E-09 |
| | | | CREB3L4 | 0.333024 | 1.01E-09 |
| | | | DYNLRB1 | 0.332146 | 1.12E-09 |
| | | | HMG20B | 0.331918 | 1.16E-09 |
| | | | USP36 | 0.331777 | 1.18E-09 |
| | | | GIT1 | 0.331498 | 1.22E-09 |
| | | | ZNF362 | 0.331356 | 1.24E-09 |
| | | | OCA2 | 0.330935 | 1.31E-09 |
| | | | R3HDM4 | 0.330365 | 1.41E-09 |
| | | | GOLGA7B | 0.330298 | 1.42E-09 |
| | | | HS1BP3 | 0.330209 | 1.44E-09 |

**Suppl. Table 6. List of CRTC3 mutations in melanoma patients.**

| Mutation type | Aminoacid change | Mutation type | Aminoacid change |
|---|---|---|---|
| Missense | ***S368F*** | Missense | ***S363L*** |
| Missense | ***P142S*** | Synonymous | ***L337-*** |
| Missense | ***P142L*** | Missense | ***P331H*** |
| Missense | ***S99F*** | Synonymous | ***V306-*** |
| Missense | ***R428Q*** | Synonymous | ***S283-*** |
| Missense | ***S332F*** | Missense | ***G266W*** |
| Missense | ***P419S*** | Missense | ***L265F*** |
| Missense | ***S345F*** | Synonymous | ***R242-*** |
| Missense | ***L166F*** | Missense | ***R240H*** |
| Missense | ***P514S*** | Missense | ***P221S*** |

| | | | |
|---|---|---|---|
| Missense | ***G510D*** | Missense | ***W187R*** |
| Missense | ***P64S*** | Missense | ***N158K*** |
| Missense | ***F47L*** | Synonymous | ***L153-*** |
| Missense | ***A288V*** | Missense | ***P142F*** |
| Missense | ***S375F*** | Missense | ***R87W*** |
| Missense | ***W229L*** | Missense | ***S71N*** |
| Missense | ***E583D*** | Missense | ***R70Q*** |
| Missense | ***M192I*** | Synonymous | ***L69-*** |
| Missense | ***G501C*** | Missense | ***Q48R*** |
| Missense | ***R525L*** | Missense | ***P372L*** |
| Fusion | ***CPEB1-CRTC3*** | Missense | ***H315N*** |
| Missense | ***P211S*** | Synonymous | ***S99-*** |
| Missense | ***S243Y*** | Synonymous | ***L395-*** |
| Missense | ***P64L*** | Synonymous | ***G604-*** |
| Truncation | **X551_splice** | Synonymous | ***L364-*** |
| Missense | ***E570D*** | Missense | ***R463C*** |
| Synonymous | ***L619-*** | Synonymous | ***S443-*** |
| Synonymous | ***C541-*** | Synonymous | ***S425-*** |
| Missense | ***P535L*** | Missense | ***T412N*** |
| Missense | ***H532N*** | Synonymous | ***S380-*** |
